# 3D single molecule localization microscopy reveals the topography of the immunological synapse at isotropic precision below 15 nm

**DOI:** 10.1101/2021.08.09.455230

**Authors:** Lukas Velas, Mario Brameshuber, Johannes B. Huppa, Elke Kurz, Michael L. Dustin, Philipp Zelger, Alexander Jesacher, Gerhard J. Schütz

**Affiliations:** Institute of Applied Physics, TU Wien, Vienna, Austria; Institute for Hygiene and Applied Immunology, Center for Pathophysiology, Infectiology and Immunology, Medical University of Vienna, Vienna, Austria; Kennedy Institute of Rheumatology, University of Oxford, Oxford, UK; Division for Biomedical Physics, Medical University of Innsbruck, Müllerstraße 44, 6020 Innsbruck, Austria

## Abstract

T-cells engage with antigen-presenting cells in search for antigenic peptides and form transient interfaces termed immunological synapses. A variety of protein-protein interactions in trans-configuration defines the topography of the synapse and orchestrates the antigen-recognition process. In turn, the synapse topography affects receptor binding rates and the mutual segregation of proteins due to size exclusion effects. For better understanding it is hence critical to map the 3D topography of the immunological synapse at high precision. Current methods, however, provide only rather coarse images of the protein distribution within the synapse, which do not reach the dimension of the protein ectodomains. Here, we applied supercritical angle fluorescence microscopy combined with defocused imaging, which allows 3-dimensional single molecule localization microscopy (3D-SMLM) at an isotropic localization precision below 15 nm. Experiments were performed on hybrid synapses between primary T-cells and functionalized glass-supported lipid bilayers. We used 3D-SMLM to quantify the cleft size within the synapse by mapping the position of the T-cell receptor (TCR) with respect to the supported lipid bilayer, yielding average distances of 18 nm up to 31 nm for activating and non-activating bilayers, respectively.

## INTRODUCTION

Understanding the topography of cellular interfaces is central for addressing many cell biological questions. The distance between the two juxtaposing cell surfaces not only regulates the affinity of protein-protein trans-interactions ^1, 2^, but the extension of the intercellular cleft also affects the spatial distribution of membrane proteins with differently-sized ectodomains ^3^. A prominent example is the size-exclusion of the large phosphatase CD45 upon contact formation between a T-cell and an antigen-presenting cell (APC), which is suspected to represent an important regulatory mechanism for the phosphorylation of the T-cell receptor (TCR) ^4^; according to this model, shifting the balance between Lck-mediated phosphorylation and CD45-mediated dephosphorylation induces downstream signaling.

A common way to study antigen-specific T-cell activation involves the use of functionalized glass-supported lipid bilayers (SLBs) as surrogates of APCs ^5, 6, 7^. This experimental design has the advantage of enabling the application of high-resolution microscopy techniques, while still preserving the essential hallmarks of T-cell signaling, including the formation of an immunological synapse, the recruitment of the kinase ZAP-70 and other downstream signaling effectors, the increase in intracellular calcium and the release of cytokines.

First, interfacing cells with a glass coverslip further allows for exploiting the interference of light reflected from the glass-water interface and light reflected from the cell membrane for imaging purposes ^8, 9^. This technique, termed interference reflection microscopy (IRM), yields high precision information on the separation of the cell surface from the surface of the glass coverslip, eventually limited only by the signal to noise ratio of the data and by the knowledge of the interference model ^9^. IRM has been frequently applied to qualitatively assess the homogeneity of T-cell adhesion to activating or inert surfaces ^5, 10, 11, 12^, often yielding patches of close contact next to areas of substantial elevation of the T-cell surface. Quantitative interpretation of the data, however, is often hampered by unknowns of the interference model ^9^. As a second advantage, studying hybrid synapses facilitates the application of total internal reflection (TIR) excitation to accentuate the signal of dyes proximal to the glass surface over intracellular background ^13^. Using TIR excitation, researchers discovered the formation of TCR microclusters upon T-cell activation ^14, 15, 16^.

More recently, single molecule localization microscopy (SMLM) has been applied to study the organization of various signaling molecules in the course of T-cell activation ^17, 18, 19, 20, 21, 22^. Briefly, SMLM achieves superior spatial resolution by the high precision to localize well separated single molecule signals that can be obtained from blinking chromophores ^23^. While a two-dimensional localization precision below 20 nm was frequently reported, it is difficult to achieve similar precision along the optical axis ^24^. In this context, studying hybrid synapses offers a third advantage: the presence of the glass coverslip in the vicinity of the fluorophores of interest allows for using the supercritical angle fluorescence as a parameter for determining the distance between the dye molecule and the glass surface ^25, 26^. Of note, the dye’s distance from the glass surface affects the shape of the recorded point spread functions due to different supercritical angle contributions. We have recently demonstrated that, when combined with defocused imaging, supercritical angle three-dimensional SMLM (3D-SMLM) achieves isotropic localization precision down to ~10 nm in all three dimensions ^27^.

Here, we used 3D-SMLM based on supercritical angle microscopy to study the topography of the immunological synapse formed between primary murine CD4^+^ T-cells and functionalized SLBs at isotropic localization precision below 15 nm. The obtained TCR localization maps were correlated with IRM images of the same synapse, thereby allowing not only cross-validation of the two approaches, but also the identification of artifacts inherent to IRM images. From the TCR z-coordinates we quantified the roughness of the T-cell surface within the synapse, as well as its separation from the SLB: Both for activating and non-activating conditions we observed multiple TCR-proximal spots of close contact, which would qualify for CD45 exclusion. Our data hence suggest that CD45 exclusion *per se* is not sufficient to explain T-cell activation. We finally quantitatively compared 3D-SMLM images with diffraction-limited TIR fluorescence microscopy of T-cell synapses to disentangle different contributions to the appearance of TCR microclusters. Our data reveal clear enrichment of TCR in microclusters, and only marginal effects due to the increased brightness of the signals of proximal TCR.

## RESULTS

### Correlative 3D-Single Molecule Localization Microscopy and Interference Reflection Microscopy

In order to evaluate the correlation between 3D-SMLM and IRM data we sought for a system with known separation of the detected single dye molecules from the glass surface. We opted for an AlexaF647-coated glass sphere of 1mm radius adhered to a glass coverslip, which yielded z-distances of up to 150 nm within the field of view of 22 × 22 µm^2^ (**Fig. 1a**). **Fig. 1b** shows an IRM image recorded next to the contact point between the sphere and the glass surface. Concentric interference fringes are clearly visible. When plotting the recorded IRM intensity values versus the distance of the according pixels from the glass surface, *z*_0_, we observed the characteristic cosine-dependence (**Fig. 1c**). In this case, three branches of the IRM intensity can be distinguished, corresponding to different orders of the interference pattern. A slight decrease in amplitude and wavelength of the recorded IRM curve is noted for increasing IRM interference orders, which is a consequence of reflections on the curved surface ^9^. In **Fig. 1b** we also included the recorded 3D-SMLM data, with the color-code indicating the calculated displacement from the glass surface, *z*_*SMLM*_. When plotted against the radial distance from the sphere’s contact point, *r*_0_, the measured single molecule displacements *z*_*SMLM*_ follow closely the surface of the sphere. (**Fig. 1d**). We further correlated *z*_*SMLM*_ with the IRM intensity values recorded on the corresponding pixels (**Fig. 1e**), yielding very good agreement of the two data sets. The black line shows the calibration curve obtained in **Fig. 1c**, the dashed lines indicate the expected Cramér-Rao lower bound (CRLB) ^27, 28^. To obtain a quantitative measure of the method’s z-precision we calculated for each localization the difference between *z*_*SMLM*_ and the theoretical *z*_0_ (**Fig. 1f**). The determined standard deviation of 21nm agrees well with the Cramér-Rao lower bound of 17nm.

**Figure 1:**
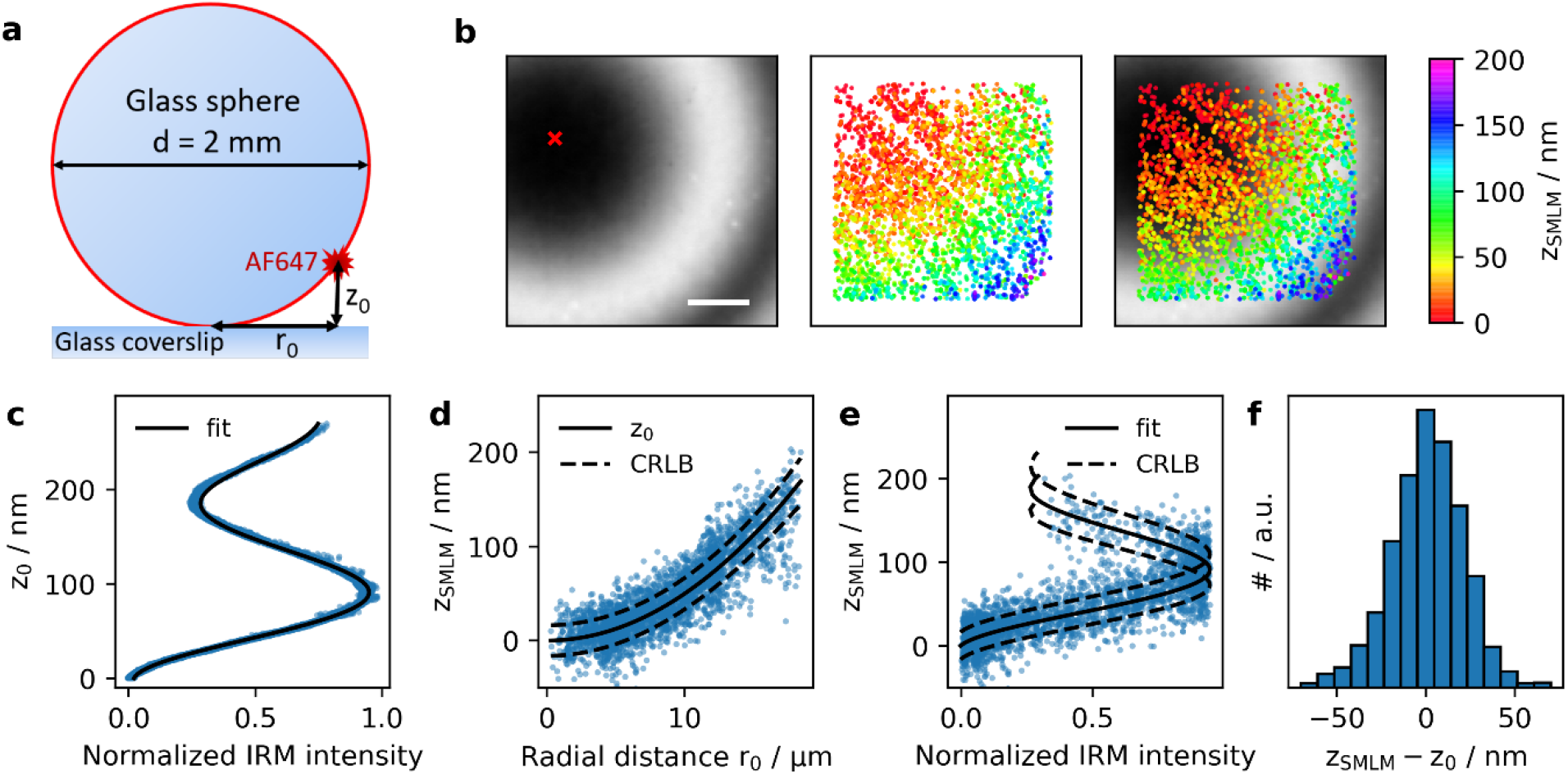
Control experiment on a fluorescently labeled glass sphere. A glass sphere of 2 mm diameter was labeled with BSA-Biotin-Streptavidin-AF647. Based on the known size of the sphere, the distance of the AF647 to the coverslip z_0_ was calculated from the measured radial distance r_0_ from the contact point of the sphere with the coverslip (a) (see Methods Eq. 1). (b) IRM and SMLM images were recorded next to the contact point of the sphere with the coverslip (red cross). (c) We plotted the z_0_ corresponding to the center of the respective pixel against the normalized IRM intensity and fitted with Eq. 2. (d) The measured distances z_SMLM_ of all localizations were plotted against their radial distance r_0_. The full line indicates the expected behavior for a spherical surface (Eq. 1), dashed lines indicate the square root of the Cramér-Rao lower bound. (e) Dependence of z_SMLM_ on the normalized IRM intensity. Full line indicates fit results from panel (c), dashed lines the square root of the Cramér-Rao lower bound. (f) The difference between the measured values z_SMLM_ and the calculated values z_0_ showed a standard deviation of 21 nm. Scale bar 5 μm.

### Correlative IRM, TIR and 3D-SMLM of the immunological synapse

Next, we applied the method to image the 3-dimensional topography of the immunological synapse formed between CD4^+^ murine 5c.c7 TCR-transgenic T-cells and stimulatory or inert surfaces. To visualize the position of the TCR, T-cells were labeled with an AlexaF647-conjugated single chain antibody fragment ^6^ against the TCR β subunit. Before analysis, all 3D-SMLM images were corrected for overcounts, which eventually allowed us to obtain a valid estimation of the surface roughness (see Methods). A mean precision of 12.7 nm, 12.0 nm, and 14.6 nm was determined for single molecule localization along the x, y, and z axis, respectively (**Supplementary Fig. 1**).

We first addressed the three-dimensional organization of the TCR in activated T-cells. To this end, T-cells were seeded onto fluid SLBs functionalized with MCC-loaded I-E^k^ at a surface density of 100 ± 30 molecules/μm^2^, which is known to stimulate intracellular calcium release ^29^; in addition, bilayers contained the adhesion molecule ICAM-1 and the costimulatory molecule B7-1 (**Fig. 2a**). For all applied conditions, T-cell activation was controlled via ratiometric calcium imaging (**Supplementary Fig. 3**). Prior to conducting imaging experiments, T-cells were fixed at specific time points (here 5 minutes) after their seeding onto the SLBs, and imaged both via IRM and 3D-SMLM. The IRM images show substantial contrast fluctuations (**Fig. 2b**), indicating corresponding fluctuations in the distance of the T-cell membrane from the SLB. This is supported by the 3D-SMLM data, where the determined TCR z-positions spread between 0 nm and 300 nm. Plotting *z*_*SMLM*_ versus the IRM intensity revealed a good correlation between the two data sets for the first branch of the IRM signal (**Fig. 2c**, black line). This correlation vanished, however, for higher order IRM branches. We attribute this lack of correlation to additional parameters affecting the measured IRM contrast such as unknown angles of the reflecting surfaces or the occurrence of multiple interferences, which particularly disturbs IRM signals originating from reflections at larger distances from the glass surface. We therefore did not fit those regions of the IRM curves. In addition, different resolutions of the two methods impede direct comparison of the two data sets: While 3D-SMLM data report on z-distances specific for 2D coordinates that can be determined with a precision below the diffraction-limit, IRM images are limited by diffraction and hence provide average values over areas given by the size of the 2D point spread function.

**Figure 2:**
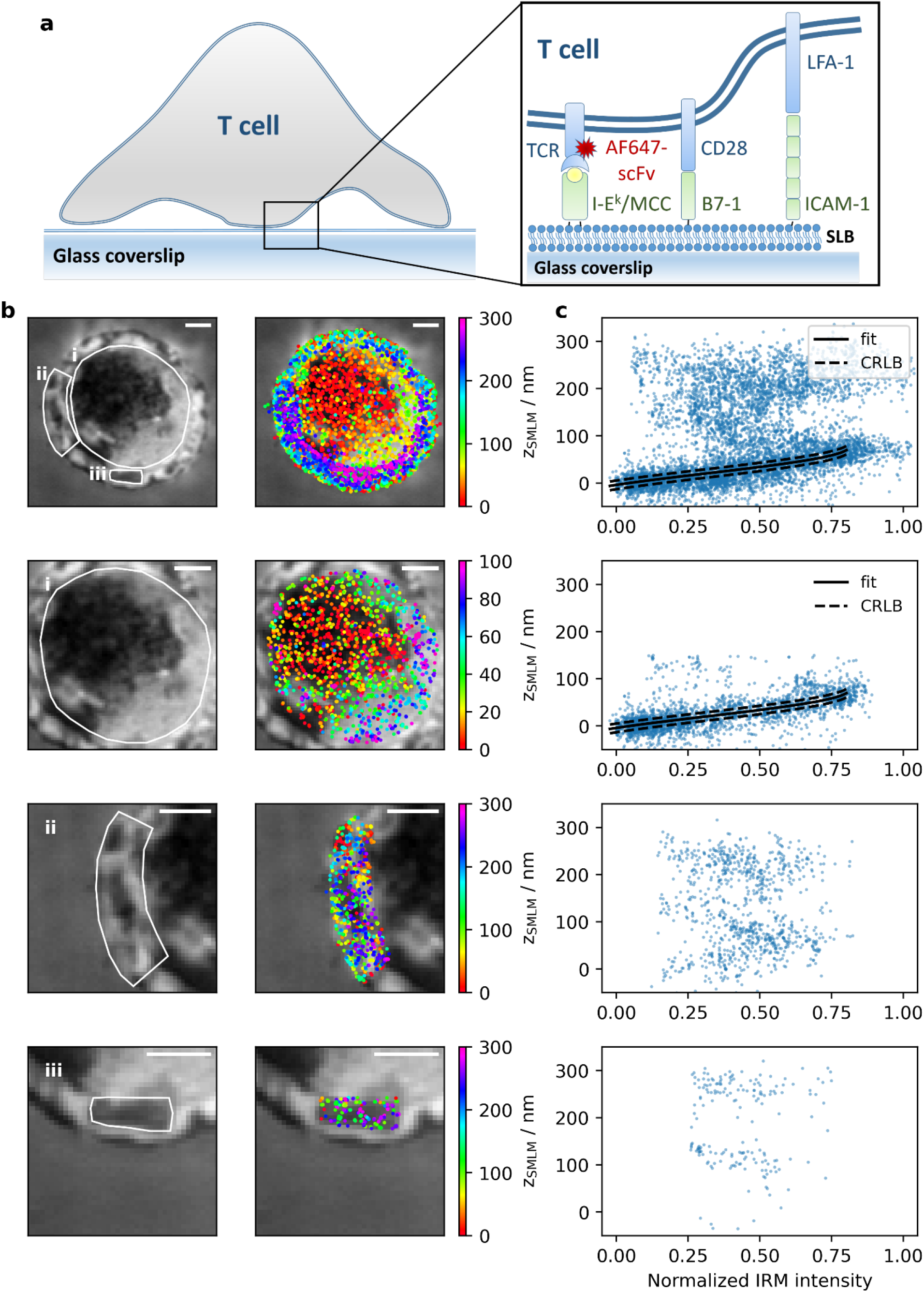
Correlative 3D-SMLM and IRM within the immunological synapse. (a) Experiments were performed on a T-cell adhering to an SLB functionalized with I-E^k^/MCC, B7-1 and ICAM-1; the TCR was labeled via AF647-conjugated H57-scFv. (b) IRM (left) and SMLM/IRM overlay images (right) of the immunological synapse. The areas i, ii and iii are shown in magnification below. (c) Correlation plots between z_SMLM_ and the normalized IRM intensity for the areas indicated in panel (b). Data points with z_SMLM_ < 100 nm were fitted with Eq. 3 (full line). The dashed lines indicate the square root of the Cramér-Rao lower bound. Scale bars 2 μm.

For detailed assessment we selected in region i) of **Fig. 2b** the cell center, in which the left half of the region was well adhered to the SLB surface while the right half featured a separation of ~60 nm from the SLB. Here, results obtained with the two imaging modalities are in very good agreement. In region ii) showing the cell edge, where the T-cell formed a narrow lamellipodium, we observed two distinct clusters in the *z*_*SMLM*_ data, one reflecting the bottom, the other the top membrane of the lamellipodium. The two clusters were separated by ~ 160 nm (**Supplementary Fig. 2**), which corresponded to the thickness of the lamellipodium ^30^. Interestingly, the z-positions of this particular region hardly correlated with the IRM patterns, likely due to high inclinations of the reflecting membrane at the lamellipodium edge possibly causing additional interferences in the IRM image. In region iii), another feature of membrane topography became apparent: the central dark IRM area did not indicate an area of close contact, but instead showed rather distal lamellipodium regions reflected by *z*_*SMLM*_ coordinates more than 100 nm away from the glass surface. Also in this region, the separation of the two lamellipodia membranes of ~160 nm became apparent from two well-separated localization clusters in the *z*_*SMLM*_ data.

It is also instructive to compare the obtained 3D SMLM and IRM images with conventional diffraction-limited TIR fluorescence microscopy images (**Fig. 3**; see also **Supplementary gallery figure 1 and 2** for additional examples). In this particular example, the cell had been fixed just before the formation of the central supramolecular activation cluster (cSMAC), when TCR microclusters were observed in a ring-like structure around the cell center (i). This image hence presumably reflects a snapshot of the directional microcluster transport towards the cSMAC ^31^. While TCRs in the ring itself showed tight contact with the glass surface – likely due to binding to the cognate pMHC –, 3D-SMLM revealed that the engulfed circular membrane patch contained TCRs substantially elevated by ~100nm (iii & ix), in agreement with the observation that the cSMAC is a site of TCR endocytosis ^32^. In addition, some of the SLB engaged TCR in the cSMAC reside in ~ 100 nm extracellular microvesicles that elevate the non-engaged TCR bearing plasma membrane by ~100 nm above the SLB ^33^, potentially contributing to the two layers of TCR in the cSMAC.

**Figure 3:**
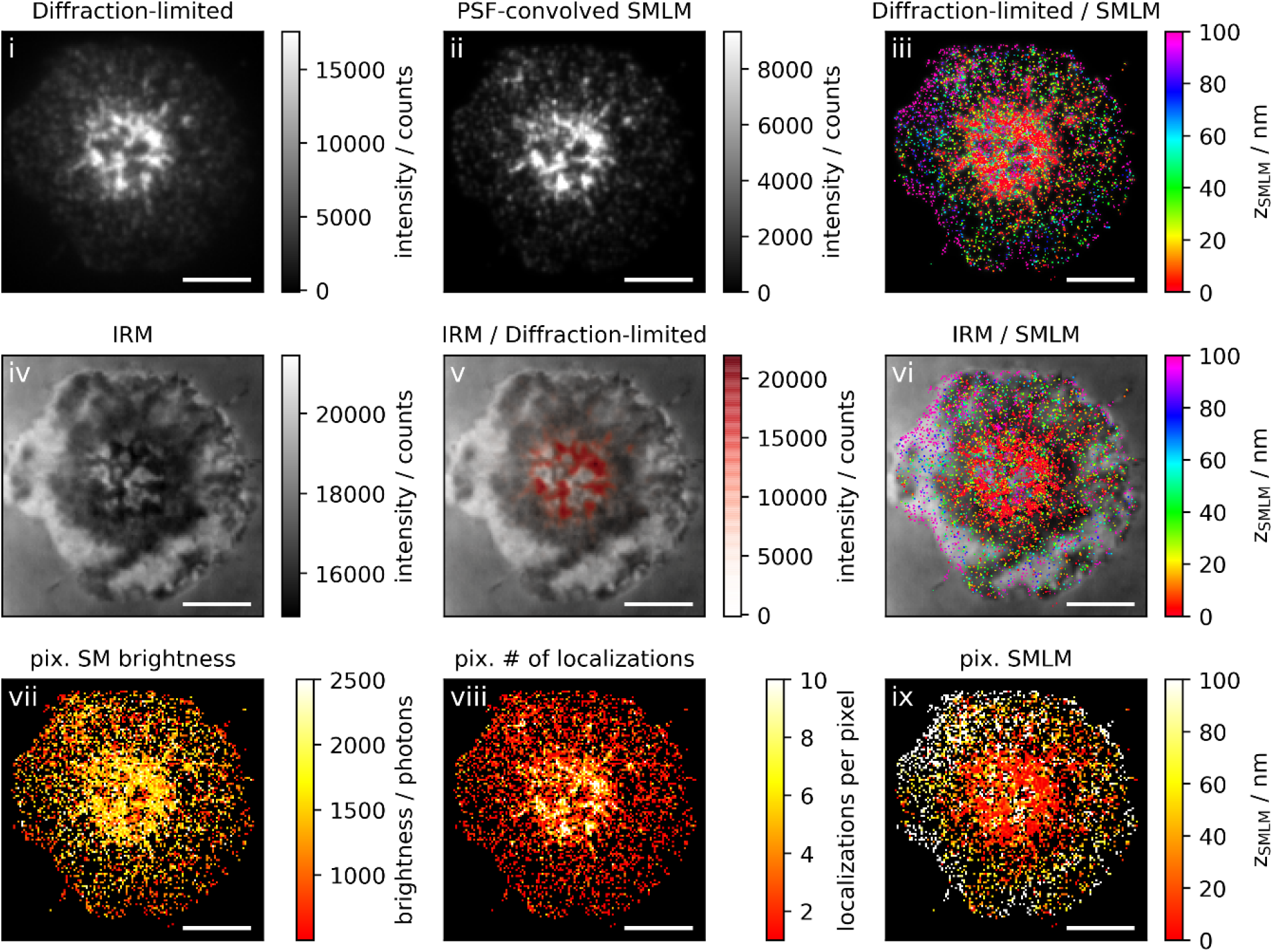
Correlative 3D-SMLM, IRM, and diffraction-limited TIR microscopy of the immunological synapse. **T-cell**s were activated on an SLB functionalized with I-E^k^/MCC, B7-1 and high densities of ICAM-1, and fixed 10 minutes post seeding. The T-cell was imaged with IRM and fluorescence microscopy: (i) Diffraction-limited TIR image of the T-cell. (ii) Reconstruction of the diffraction-limited image by convolving the 3D-SMLM image with the corresponding psf. (iii) Overlay of the diffraction-limited TIR image with the 3D-SMLM image. The color-code indicates distances to the coverslip z_SMLM_. (iv) IRM image. (v) Overlay of the IRM image with the diffraction-limited image. (vi) Overlay of the IRM and the 3D-SMLM image. Bottom row images were generated by calculating the pixel-wise average of the 3D-SMLM images (pixel size of 146 nm is consistent with diffraction-limited image) according to pixelated mean single molecule (SM) brightness (vii), pixelated number of localizations (viii) and pixelated mean z_SMLM_ (ix). Scale bars 5 μm.

We next analyzed TCR microclusters in more detail. In principle, TCR microcluster contrast is not only determined by protein enrichment, but also by the closer proximity of the fluorophores to the glass surface, yielding both increased excitation intensity in the evanescent field as well as increased detection efficiency due to the collection of supercritical angle fluorescence. Since our method allows for disentangling single molecule fluorescence brightness, z-position, and local clustering, we addressed the different contributions to the diffraction-limited TIR images. For this, we compared the diffraction-limited TIR images (**Fig. 3i**) with data obtained from 3D-SMLM. The single molecule brightness indeed showed some correlation with the positions of microclusters in the diffraction-limited image **(Fig. 3vii)**, which was also reflected in a map of the single molecule z-positions **(Fig. 3ix)**. Furthermore, single molecule localizations were strongly clustered at the positions of the microclusters **(Fig. 3viii)**. We sought to reconstruct the diffraction-limited image by convolving the single molecule localization map with the brightness-weighted point spread function (psf) **(Fig. 3ii)**. The reconstructed image agreed well with the original diffraction-limited image, down to the level of the individual TCR microclusters.

To quantitatively disentangle the different contributions, we identified TCR microclusters by intensity-thresholding the diffraction-limited images (**Supplementary Fig. 6**), and analyzed the single molecule properties separately for localizations coinciding with the TCR microcluster regions versus localizations outside of TCR microclusters. **Fig. 4** shows the ratios of single molecule brightness and number of single molecule localizations per pixel, which were obtained from multiple cells. While the brightness levels in microclusters increased only by 20%, we observed about 3-fold enrichment of localizations. The product of the two ratios quantitatively matched the average brightness ratios of pixels corresponding to microclusters versus pixels of outside regions obtained from the diffraction-limited images. Importantly, as this analysis was based on pixel-wise discrimination of microclusters in the diffraction-limited images, it was not affected by residual overcounts in the SMLM images. Taken together, we conclude that the increased brightness of TCR microclusters in diffraction-limited images is mainly explained by enrichment of TCR molecules. Of note, we observed the expected increased brightness of microclusters fixed at late time-points (red data points) compared to those fixed at early time-points (orange data points) ^16^; also this effect is explained by an increase in the number of single molecule localizations.

**Figure 4:**
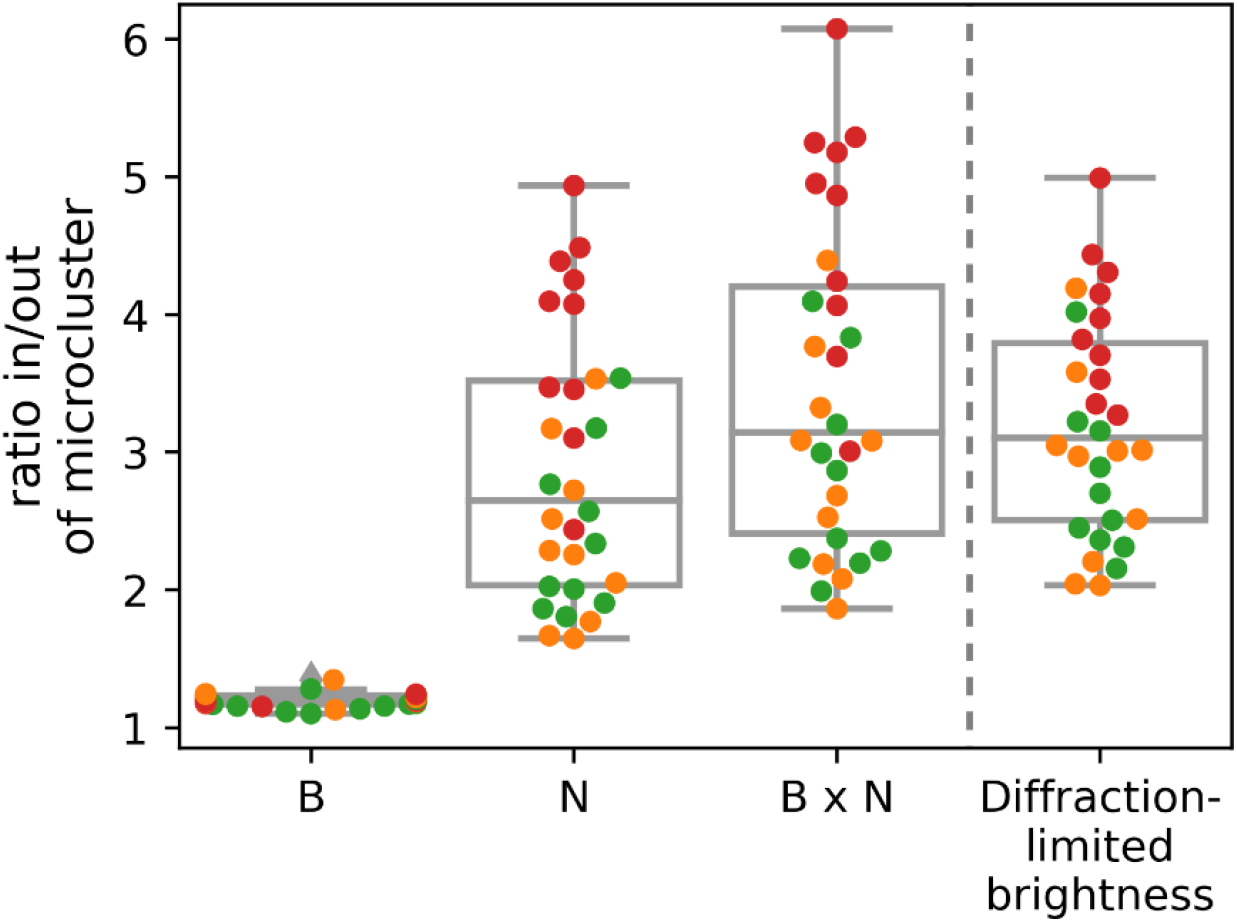
Disentangling single molecule brightness and molecular enrichment in TCR microclusters: We quantified the average single molecule brightness, B, and the number of localizations, N, in pixels corresponding to TCR microclusters (“in”) and the complementary regions of the synapse (“out”). Plotted are the ratios *in/out* for B, N, the product B x N, and the average diffraction-limited brightness per pixel. Each data point corresponds to the average ratio per cell. Colors indicate the time point of fixation post seeding (orange: 5-10 minutes, green: 10 minutes, red: 10-15 minutes). (n=30 cells).

We further studied T-cells contacting SLBs that were functionalized with the adhesion molecule ICAM-1 only, so that no calcium signal was triggered (see **Supplementary Fig. 3** for ratiometric calcium analysis and **Supplementary gallery figure 3 and 4** for exemplary images). As expected, we did not observe the formation of TCR microclusters. While in general the cells spread well on such substrates, areas of close contact appeared more fragmented than in the activated situation. This effect was more pronounced when we reduced the density of the adhesion molecule ICAM-1 in the SLB from 125 ± 22 molecules/μm^2^ to 4 ± 2 molecules/μm^2^.

### Quantitative Analysis of T-cell surface topography within the immunological synapse

For quantification, we determined the roughness of the T-cell surface within the immunological synapse. To prevent the inclusion of data originating from the top membrane of lamellipodia we only considered localizations with *z*_*SMLM*_ ≤ 100 nm. At low ICAM-1 densities, we observed substantial fluctuations of the recorded z-positions (**Fig. 5a**). For quantitative determination of the surface roughness, we compared the obtained variances 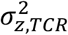 with values obtained for fluorescently labeled SLB-anchored I-E^k^/MCC, 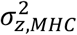, according to 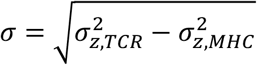, yielding a standard deviation of *σ* = 37 nm. Fluctuations decreased to 29 nm when we increased the density of the adhesion molecule ICAM-1 in the lipid bilayer. Upon activation via higher densities of I-E^k^/MCC, T-cells adhered more smoothly to the surface, as indicated by reduced overall z-fluctuations of 19 nm. Also, in case of activation we observed the T-cell surface flattening out to a considerable extent with ICAM-1 present at higher densities.

**Figure 5:**
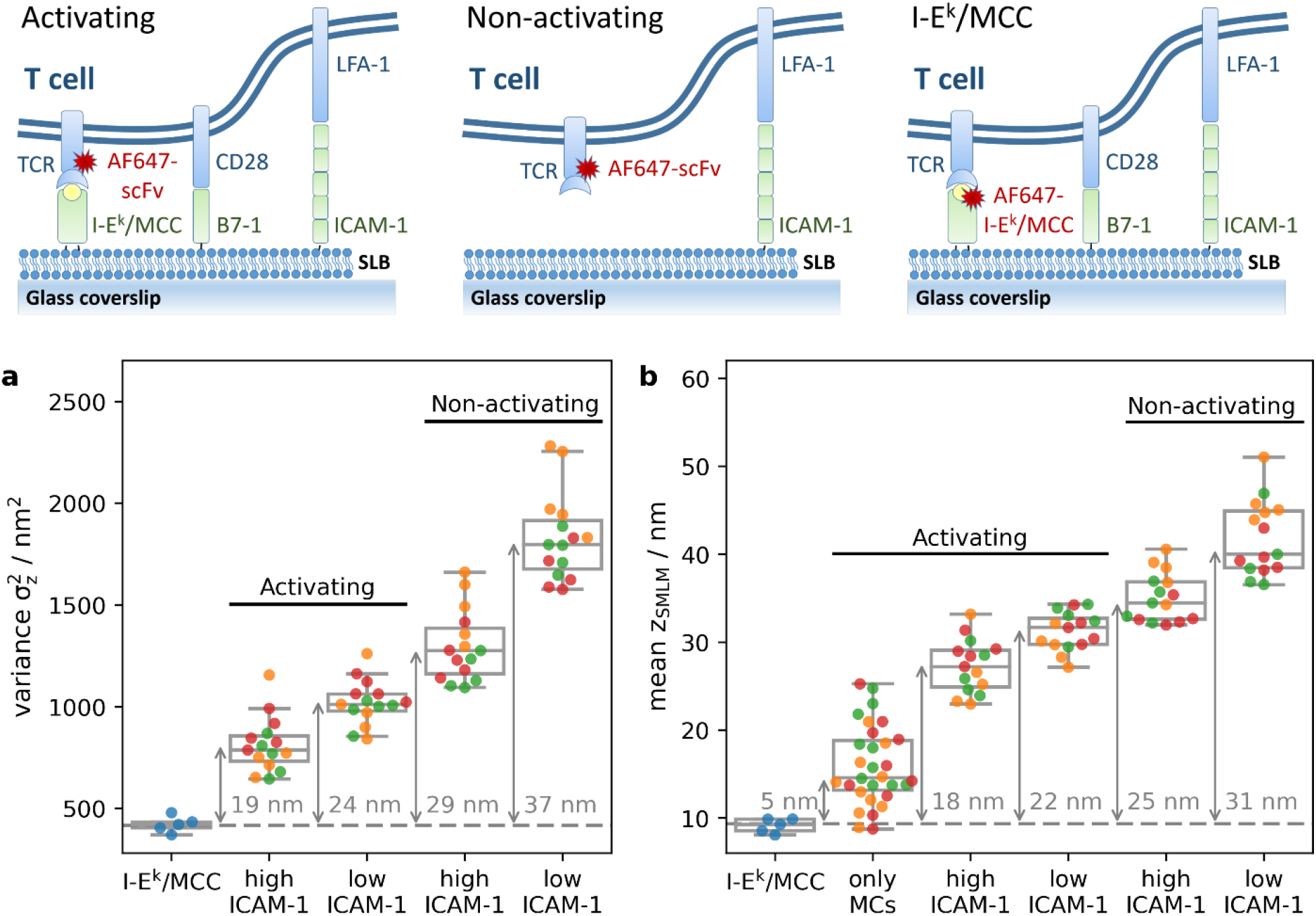
Contact analysis for T-cells recorded under activating and non-activating conditions: **T-cell**s were seeded on SLBs either functionalized with I-E^k^/MCC, B7-1 and ICAM-1 (termed “activating”) or with ICAM-1 only (termed “non-activating”). To vary adhesive strength, we used 125 ICAM-1 molecules per µm^2^ (termed “high ICAM-1”) or 4 ICAM-1 molecules per µm^2^ (termed “low ICAM-1”). T-cells were allowed to spread on SLBs and were fixed after 5-10 (orange), 10 (green) or 10-15 (red) minutes post seeding. As a control, unlabeled T-cells were seeded on SLBs containing I-E^k^/MCC labeled with AF647. The recorded localizations were filtered for z_SMLM_ < 100 nm in order to exclude contributions from the upper surface of lamellipodia. (**a**) The variance of z_SMLM_ per cell was plotted for different conditions. Arrows indicate the difference in variance to the control I-E^k^/MCC data, numbers the corresponding square root. (**b**) Mean z_SMLM_ per cell was plotted for different conditions. Arrows indicate the difference to the control I-E^k^/MCC data. To calculate the mean distance of TCR microclusters from the glass surface we considered only localizations within microclusters (termed “only MCs”) in experiments performed both at high and low densities of ICAM-1. (n = 15 cells per condition).

According to the kinetic segregation model, the axial dimension of the intercellular cleft determines the accessibility of the large phosphatase CD45. We hence quantified the absolute distance of the TCR from the SLB, when compared to fluorescently labeled SLB-anchored I-E^k^/MCC. Generally, we observed similar trends as for the standard deviations: with increasing densities of I-E^k^/MCC and ICAM-1 the TCR was observed to be closer to the SLB surface (**Fig. 5b**). The separation varied between 18 nm at high densities of I-E^k^/MCC and ICAM-1 up to 31 nm for scanning T-cells recorded at low densities of ICAM-1. When selecting only signals corresponding to TCR microclusters for our distance analysis, we observed the expected close contact between TCR and I-E^k^/MCC, with a calculated separation of 5 nm. The residual separation reflected in all likelihood the distance separating the dye site-specifically conjugated to the single chain antibody fragment and the dye coupled to the MCC peptide’s C-terminus as presented by I-E^k 6^.

## DISCUSSION

We applied here a novel 3D-SMLM method to map and analyze the position of the TCR within the immunological synapse at isotropic localization precision below 15 nm. We studied the formation of hybrid synapses between T-cells and glass-supported functionalized lipid bilayers. Our study addressed three questions as is outlined below:

### i) To what extent does IRM contrast report on the position of the T-cell membrane?

IRM contrast arises from interferences due to optical path-length differences between the beam reflected at the glass-water interface and beams reflected from surfaces within the sample. While in principle IRM can be calibrated such that it allows for quantification of absolute distances, a few shortcomings hamper such quantification in biological samples. For example, tilted membranes or the presence of second or third order interferences are difficult to account for ^9^. In addition, multiple reflections from different layers of varying refractive index induce phase shifts in the IRM intensity profiles, thereby impeding absolute distance measurements. In fact, protein ectodomains and the glycocalyx contribute to the change in refractive index between the aqueous environment and the cell, rendering the plane of reflection rather poorly-defined. Finally, IRM images are of diffraction-limited spatial resolution, and thereby yield averages of the interference contrast over a few 100 nm. Fluctuations at smaller scales would hence be averaged out.

Recording 3D-SMLM localization maps is fundamentally different from IRM. Using supercritical angle detection combined with defocused imaging, our method essentially determines the three-dimensional position of all visible dye molecules. We employed here an AlexaF647-conjugated single chain antibody fragment that specifically recognizes the TCR β subunit ^6^. In previous studies we have shown that labelling neither activates T-cells nor impedes specific antigen recognition ^6^. Of note, we used TIR excitation in order to confine imaging to the synapse region. While the evanescent field was narrow enough to prevent contributions from TCRs at the top of the T-cell, we could observe both the bottom and the top membranes of lamellipodia.

Indeed, there was in general a good qualitative agreement between the two imaging modalities. Substantial differences, however, arose when distal membrane regions contributed to the IRM contrast values: Then, contrast values did not unequivocally correspond to the axial membrane separation. This was especially observed in lamellipodia, where both the bottom and the top membrane contributed to the signal.

### ii) What is the physical reason for the observation of TCR microclusters?

In TIR excitation, fluorescent molecules close to the glass coverslip naturally contribute with higher brightness than distal molecules. This effect is further amplified by the collection of supercritical angle fluorescence when using high NA objectives ^25^. Taken together, what appears as a bright spot in diffraction-limited fluorescence microscopy could be the consequence of increased density or of increased brightness of the fluorophores. Given the close contact between TCR microclusters and the surface ^12^ researchers became concerned, whether microclusters actually reflect TCR enrichment.

3D-SMLM provides ground truth information on the origin of apparent TCR microclusters in diffraction-limited TIR microscopy, as it allows for disentangling molecular enrichment from brightness changes. We observed only marginal contributions from single molecule brightness increase; as brightness was largely attributable to molecular enrichment. This finding is in accord with our observation of a rather smooth interface between the SLB and the T-cell membrane under activating conditions, so that there are globally only minor variations in the single molecule’s z-distance in comparison to the TIR penetration depth.

### iii) Does the observed cleft size support CD45 segregation?

A prominent model for TCR triggering involves the balance in the activities of the kinase lck and the phosphatase CD45 for ITAM phosphorylation. In this kinetic segregation model, proteins with bulky extracellular domains – such as the large phosphatase CD45 – are proposed to be segregated locally from the comparably short pMHC-TCR complexes ^4^. While there are ample of reports that would be consistent with this model ^34, 35, 36, 37, 38^ the community has not reached a consensus yet ^39, 40, 41^.

One difficulty has been the precise measurement of the cleft size between the T-cell surface and the opposing membrane. Given the dimension of the TCR-pMHC-CD4 ternary complex of 10 nm ^42^ and our apparent distance measurements between TCR and I-E^k^/MCC within TCR microclusters of 5 nm, the obtained distance undervalues the cleft size by 5 nm. Upon correcting for this effect, an average cleft size within the whole synapse ranging between 23 nm and 36 nm for the conditions shown in **Fig. 5b** can be estimated. In addition to the average cleft size, however, distance fluctuations within the synapse were of the same order (**Fig. 5a**), indicating the presence of multiple contact sites between the two membranes. Assuming an axial length of CD45R0, the smallest CD45 isoform, of approximately 22 nm ^34^, our measurements hence indicate the existence of numerous membrane contact sites that would be too narrow to host CD45, both for resting and activating conditions. In particular, an average cleft size of 30 nm together with an average surface roughness of 29 nm, as observed for scanning T-cells at high ICAM-1 densities, render the presence of multiple CD45 exclusion zones likely. Similar data were recently reported for the tips of microvilli which showed segregation of the TCR and CD45 prior to T-cell activation ^43^. Given that none of these scenarios promoted T-cell activation it is possible that the ITAMs of these segregated TCRs are not accessible to kinases ^44^. In addition, TCR-CD45 segregation may be too transient to trigger stable phosphorylation of TCR ITAMs. Kinetic data about protein mobility in conjunction with the superresolution images would be needed in order to obtain a quantitative understanding of these key aspects of the TCR triggering process.

## MATERIALS AND METHODS

### Reagents and proteins

Phosphate buffer saline (PBS), Hanks’ balanced salt solution (HBSS), biotin labeled bovine albumin (BSA-biotin), fetal bovine serum (FCS), Histopaque-1119, glucose, glucose oxidase, catalase and cysteamine were purchased from Sigma Aldrich (Merck KGaA, Germany). 1,2-dioleoyl-sn-glycero-3-phosphocholine (DOPC), 1,2-dipalmitoyl-sn-glycero-3-phosphocholine (DPPC) and 1,2-dioleoyl-sn-glycero-3-[(N-(5-amino-1-arboxypentyl)iminodiacetic acid)succinyl] (nickel salt) (Ni-NTA-DGS) were from Avanti Polar Lipids, Inc. (USA). Streptavidin-AF647, IL-2 and formaldehyde were from Thermo Fischer scientific (USA). Cell medium RPMI 1640, penicillin, streptomycin, glutamate, sodium pyruvate, 1x non-essential amino acids and mercaptoenthanol were from Gibco, Life Technologies (USA). B7-1 and ICAM-1 were from Sino Biological (China). I-E^k^ and MCC peptide were prepared and labeled as described previously ^6^. H57-scFv was expressed in inclusion bodies, refolded, site-specifically labeled with maleimide functionalized AF647 (Thermo Fisher Scientific, USA), and purified as described in ^6^.

### Coating of glass spheres

Glass spheres (d=2mm, Schott, Germany) were plasma-cleaned (PDC-002, Plasma Cleaner Harrick Plasma, USA) for 10 minutes and incubated with 5 % BSA-biotin for 60 minutes. The spheres were then washed with PBS and incubated with 1 μg/ml Streptavidin-AF647 for 30 minutes. Afterwards, spheres were rinsed with PBS and placed into 8-well chambers (Nunc Lab-Tek, Thermo Scientific, USA) with plasma cleaned glass coverslip (MENZEL-Gläser Deckgläser 24 × 60 mm #1.5, Thermo Fisher Scientific, USA) for imaging.

### Supported lipid bilayers

DOPC and Ni-NTA-DGS were dissolved in chloroform and mixed in a glass tube at a molar ratio of 98 (DOPC) : 2 (Ni-NTA-DGS), and subsequently dried under N_2_ flow for 20 minutes. Next, the lipid mixture was resuspended in 1 ml of PBS and sonicated for 10 minutes (ultrasound bath USC500TH, VWR, England) to form small unilamellar vesicles (SUVs). Glass slides (MENZEL-Gläser Deckgläser 24 × 60 mm #1.5, Thermo Fisher Scientific, USA) were plasma cleaned for 10 minutes and glued on 8-well chambers. To form supported lipid bilayers, the freshly cleaned glass slides were incubated with SUVs for 20 min and washed extensively with PBS. Next, SLBs were incubated with His-tag proteins (Activating conditions: 10 ng I-E^k^/MCC, 50 ng B7-1 and 30 ng or 0.3 ng ICAM-1; resting conditions: 30 or 0.3 ng ICAM-1) for 1 hour, and extensively rinsed with PBS. Before seeding the cells, PBS was exchanged for HBSS. Immobile SLBs were prepared identically, except for the following changes: we used a lipid mixture of 98 % DPPC and 2 % Ni-NTA-DGS. For sonication the ultrasound bath was preheated to 50 °C and coverslips were incubated with SUVs on a hot plate (65°C). Finally, SLBs were incubated with 10 ng of I-E^k^/MCC-AF647.

### T-cells

T-cells were obtained from transgenic 5c.c7 mice as described previously ^45^. Briefly, T-cells were isolated from lymph nodes and stimulated with 2 μM HPLC-purified moth cytochrome C (MCC) peptide (ANERADLIAYLKQATK, Intavis Bioanalytical Instruments) in a T-cell medium (RPMI 1640 containing 10% FCS, 100 U/mL penicillin/streptomycin, 2mM glutamate, 1 mM sodium pyruvate, 1x non-essential amino acids, and 50 μM-Mercaptoenthanol). Culture volume was doubled and 100 U/mL IL-2 was added on day 2. On days 3 and 5, T-cell cultures were expanded in a ratio of 1:1. On day 6, dead cells were removed by centrifugation trough a Histopaque-1119 cushion. T-cell experiments were performed on days 7–9 after initial stimulation.

### Animal model and ethical compliance statement

5c.c7 αβ TCR-transgenic mice bred onto the B10.A background were a kind gift from Michael Dustin (University of Oxford, UK). Both male and female mice at 8-12 weeks old were randomly selected and sacrificed for isolation of T-cells from lymph nodes and spleen, which was evaluated by the ethics committees of the Medical University of Vienna and approved by the Federal Ministry of Science, Research and Economy, BMWFW (BMWFW-66.009/0378-WF/V/3b/2016). Animal husbandry, breeding and sacrifice of mice was performed in accordance to Austrian law (Federal Ministry for Science and Research, Vienna, Austria), the guidelines of the ethics committees of the Medical University of Vienna and the guidelines of the Federation of Laboratory Animal Science Associations (FELASA), which match those of Animal Research: Reporting in vivo Experiments (ARRIVE). Further, animal husbandry, breeding and sacrifice for T-cell isolation was conducted under Project License (I4BD9B9A8L) which was evaluated by the Animal Welfare and Ethical Review Body of the University of Oxford and approved by the Secretary of State of the UK Home Department. They were performed in accordance to Animals (Scientific Procedures) Act 1986, the guidelines of the ethics committees of the Medical Science of University of Oxford and the guidelines of the Federation of Laboratory Animal Science Associations (FELASA), which match those of Animal Research: Reporting in vivo Experiments (ARRIVE).

### TCR staining for imaging

In order to label the T-cell receptor, appr. 10^6^ cells were washed twice by centrifugation (300 RCF for 3 min) with 2 ml of HBSS containing 2% FCS. Next, T-cells were incubated with H57-scFv conjugated with AF647 at saturating conditions for 20 min on ice. After incubation the cells were washed twice by centrifugation with 2 ml of HBSS + 2% FCS at 4°C. The cells were then immediately seeded on the prepared SLBs and fixed after 5-15 minutes of spreading with 4% formaldehyde + 0.2% glutaraldehyde (Serva, Germany) for 10 min. Afterwards, the fixing solution was rinsed with PBS. Finally, PBS was exchanged with blinking buffer for dSTORM (PBS (pH 7.4), 10% glucose, 500 μg/ml glucose oxidase, 40 μg/ml catalase and 50 mM cysteamine) and cells were imaged immediately.

### Microscopy

The home-built experimental setup was based on an Olympus IX73 (Japan) microscope body equipped with a high NA objective (Carl Zeiss, alpha-plan apochromat, 1.46 NA, 100x, Germany) (**Supplementary Fig. 5**). Samples were illuminated with 640 nm laser light (100 mW nominal laser power, OBIS Laser box, Coherent, USA) coupled by an optical fiber into a high-speed galvo back focal scanner (iLas^2^, Visitron Systems, Germany). The iLas^2^ system was used for rapid spinning of the laser in the back focal plane of the objective (spinning TIR configuration), yielding an excitation intensity of 1 kW/cm^2^ at the sample. Finally, lasers were reflected to the specimen and filtered from the fluorescence signal by a quad dichroic mirror (Di01-R405/488/532/635, Semrock, USA) and an emission filter (ZET405/488/532/642m, Chroma, USA) placed in the upper deck of the microscope body. The same dichroic mirror was used for coupling in light from a blue LED (455 nm, Thor Labs, USA) for IRM imaging. An additional shortpass beamsplitter (HC BS 750 SP, Semrock, USA) was placed in the lower deck of the microscope for coupling in a home-built focus hold system based on a 785 nm laser diode (LM9LP, Thor Labs, USA), a camera (ac640-750um, Basler, Germany) and an objective piezo stage (P721.SL2, Physik Instrumente, Germany). Finally, the specimen was imaged on a camera (EM-CCD Ixon Ultra, Andor, UK). Illumination and image acquisition was operated by VisiView (Visitron Systems, Germany).

We used defocused imaging for determining the three-dimensional single molecule position, as described previously (Zelger et al 2020). Briefly, we used dSTORM to record spatially separated single molecule signals of the fluorescently labeled T-cells. Before the experiment, single molecules adhered to the glass coverslip next to the cells were brought into focus, and the objective was displaced by 500 nm towards the sample by the piezo stage (P721.SL2, Physik Instrumente, Germany). For the dSTORM experiment, 20.000 frames at 20 ms illumination time and 9 ms delay time were recorded for each analyzed image. After 10.000 frames a 405 nm UV laser was added to the illumination sequence (1 mW nominal laser power, OBIS Laser box, Coherent, USA). The distance of the objective to the coverslip was maintained by the focus hold system for the entire recording of the cells.

### Data Analysis

#### Localization of single molecules

The single molecule coordinates x, y, and z, the single molecule brightness (B) and the background signal (bg) were determined by maximum likelihood fitting of the calculated psf model on a 21 by 21 pixel subregion around the signal, as previously described ^46^. The psf model includes optical aberrations of the experimental setup retrieved from z-stack measurements of 100 nm fluorescent beads (TetraSpeck Microspheres, Thermo Fisher Scientific, USA). The defocus value was adjusted in the software so that the lowest localizations in z appeared close to 0. Single molecule signals closer than 1500 nm were discarded in order to avoid psf overlaps.

#### SMLM corrections

It is necessary to correct for field-dependent tilt aberrations, as they would introduce bias into the determined z-positions of the localized signals. For this purpose, we imaged immobile supported lipid bilayers functionalized with I-E^k^/MCC AF647 as a ground truth sample, in which all dye molecule should be located in the same z-plane. The localizations were fitted with a planar surface, which was then subtracted from all localizations for correction. In order to correct for overcounting artifacts, localizations were merged by a tracking algorithm (Trackpy) ^47^, yielding the average three-dimensional position and the average single molecule brightness. Parameters of Trackpy were the three-dimensional search radius, and the maximally allowed size of gaps within a given trajectory which may occur due to molecular blinking or fitting problems. Both parameters should be chosen sufficiently large to include most blinks originating from the same molecule, and sufficiently small to avoid connecting blinks from different molecules. To identify an appropriate choice of the three-dimensional search radius and the maximally allowed gap size, we analyzed the standard deviation of all determined z-coordinates *σ*_*z*_. Beyond a search radius of 60 nm and a gap size of 5 we observed a plateau in *σ*_*z*_ (**Supplementary Fig. 4a**); in the following we hence analyzed all data with a search radius of 60 nm and a gap size of 5. To find out whether the obtained values of *σ*_*z*_ correspond to the underlying surface roughness we simulated 3D-SMLM experiments (**Supplementary Fig. 4b**), using experimentally derived single molecule blinking statistics and localization precision. For a simulated surface roughness of 26 nm we found high quantitative agreement between our simulations and data recorded on T-cells seeded on activating supported lipid bilayers. For the selected values of 60 nm search radius and 5 gaps we overestimated the surface roughness only by 1.5 nm. Finally, localizations were filtered for the single molecule brightness B (500 < B < 4000) in order to avoid fitting of camera noise or of strong signals arising from overlapping molecules.

#### Data recorded on a glass sphere

First, the contact point of the sphere to the glass coverslip was determined by fitting a 2D second-order polynomial to the IRM intensity profile in **Fig**.**1b** between the central intensity minimum and the first intensity maximum. The expected distance z_0_ of the glass sphere from the coverslip was calculated via

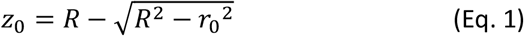

where *R* = 1 mm is the radius of the sphere and *r*_0_ is the radial distance to the contact point. The dependence of the normalized IRM intensity on the expected distance of the sphere to the coverslip z_0_ was fitted with

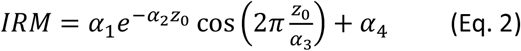

yielding *α*_1_ = −0.54, *α*_2_ = 0.0036 nm^-1^, *α*_3_ = 188,42 nm, *α*_4_ = 0.56.

For **Fig. 1d** we put the origin of the z-coordinates to the surface of the glass coverslip, to which we referenced all obtained z-positions. In **Fig**.**1e** the distance of the localizations to the coverslip z_SMLM_ was correlated with the IRM intensity of the corresponding pixels and overlaid with the fit from Eq. 2. The Cramér-Rao lower bounds shown as dashed lines were calculated considering the localizations’ brightness, background level, z position and the psf model including aberrations.

#### Data recorded on T-cells

All z-positions were referenced against z-positions determined from signals observed around the cells, which correspond to traces of fluorescent H57-scFv molecules adsorbed to the glass surface.

In **Fig**.**2c**, localizations were plotted versus the IRM intensity of the corresponding pixels and the localizations with z_SMLM_ < 100 nm were fitted with

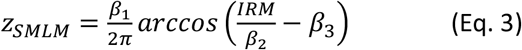

yielding *β*_1_ = −0.50, *β*_2_ = 225.69 nm, *β*_3_ = 0.39.

To determine the localization precision for x, y and z direction (**Supplementary Fig. 1**) we first calculated the standard deviation of localizations from signals occurring in subsequent frames within a three-dimensional distance of 60 nm and a maximum gap size of 5, yielding the single frame localization precision *σ*_1,*i*_ for molecule *i*. Merged localizations show an improved localization precision 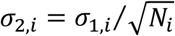, with *N*_*i*_ the number of observations per molecule. The overall localization precision is then given by 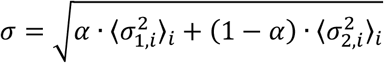, where *α* denotes the fraction of molecules observed only once. CRLBs were calculated from the estimates for each single molecule observation, *CRLB*_*i*_, accordingly. The histograms in **Supplementary Fig. 1** show the weighted sums of the normalized histograms of *σ*_1,*i*_ and *σ*_2,*i*_.

The z histogram of the lamellipodium shown in Fig. 2b (ii) was fitted with a sum of 2 gaussian functions with center positions at 68 and 228 nm (**Supplementary Fig. 2**).

To reconstruct diffraction-limited images from SMLM recordings (**Fig. 3ii**), localizations were convolved with the psf model taking into account the single molecule brightness B and the z position. The psf model was calculated for a focus position at z = 0 and included the aberrations of our optical setup.

For specific analysis of TCR microclusters, we ascribed pixels to TCR microclusters based on intensity thresholding of the diffraction limited images of activated T-cells (threshold: MC intensity per pixel > 1.5-times mean intensity per pixel of the whole cell) (see **Supplementary Fig. 6** for an example). Localizations were assigned either to microcluster pixels (termed “in”) or to the complementary pixels (termed “out”). The ratio in/out of the single molecule brightness values was calculated based on the single molecule brightness B. The density ratio was calculated from counting the number of localizations, N, per IN and OUT area.

### Simulations

To produce a continuous, rough surface we first generated a random 3D surface on a 10 nm grid with a size of 10 × 10 μm^2^ by placing normally distributed z-coordinates on every grid-point; the normal distribution was centered around zero with a standard deviation of 26 nm. Next, a Gaussian smoothing with a standard deviation of 0.5 μm (in x/y) was applied, and the z-coordinates were rescaled to preserve a standard deviation of 26 nm in z. Finally, the resulting surface was shifted along the z-axis so that the lowest point of the surface yielded z=0 nm. Onto this surface, we placed 1000 molecules randomly distributed in the x/y plane. For each molecule a blinking sequence of on-and off-states was generated (total of 20000 frames), using the single molecule blinking statistics determined by analyzing the blinking kinetics of AF647 in underlabeled T-cells. We observed a slight z-dependence of the mean on-time 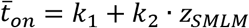, with *k*_1_ = 1.5 *frames* and *k*_2_ = 0.01 *frames /nm*, which was taken into account by including each molecule’s z-position; the off-time 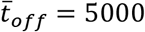*frames* was largely independent of the z-coordinate. Both on- and off-times were assigned using exponential distributions with the corresponding mean values 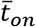 and 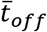. Localizations were assigned to each molecule’s position in the on-periods according to the determined localization errors of 12 nm in x and y direction. In case of localization errors along the z-direction we took into account the decreased excitation along the optical axis upon using TIR excitation, which was determined from underlabeled cells; we obtained *σ*_*z*_ = *k*_3_ *+ k*_4_ *· z*_*SMLM*_, with *k*_3_ = 12 *nm* and *k*_4_ = 0.05. The final data set was used for merging analysis via Trackpy with the different search radii and gap sizes (see **Supplementary Fig. 4**).

## Supporting information

Supplementary Image Gallery

## ACKNOWLEDGEMENTS

The study was supported by the Austrian Science Fund (FWF) (P 30214-N36, P 32105-N28), by the European Union’s Horizon 2020 research and innovation programme under the Marie Skłodowska-Curie grant agreement (721358), and by the Wellcome Trust 100262Z/12/Z and the Kennedy Trust for Rheumatology Research.

## SUPPLEMENTARY FIGURES

**Supplementary Figure 1:**
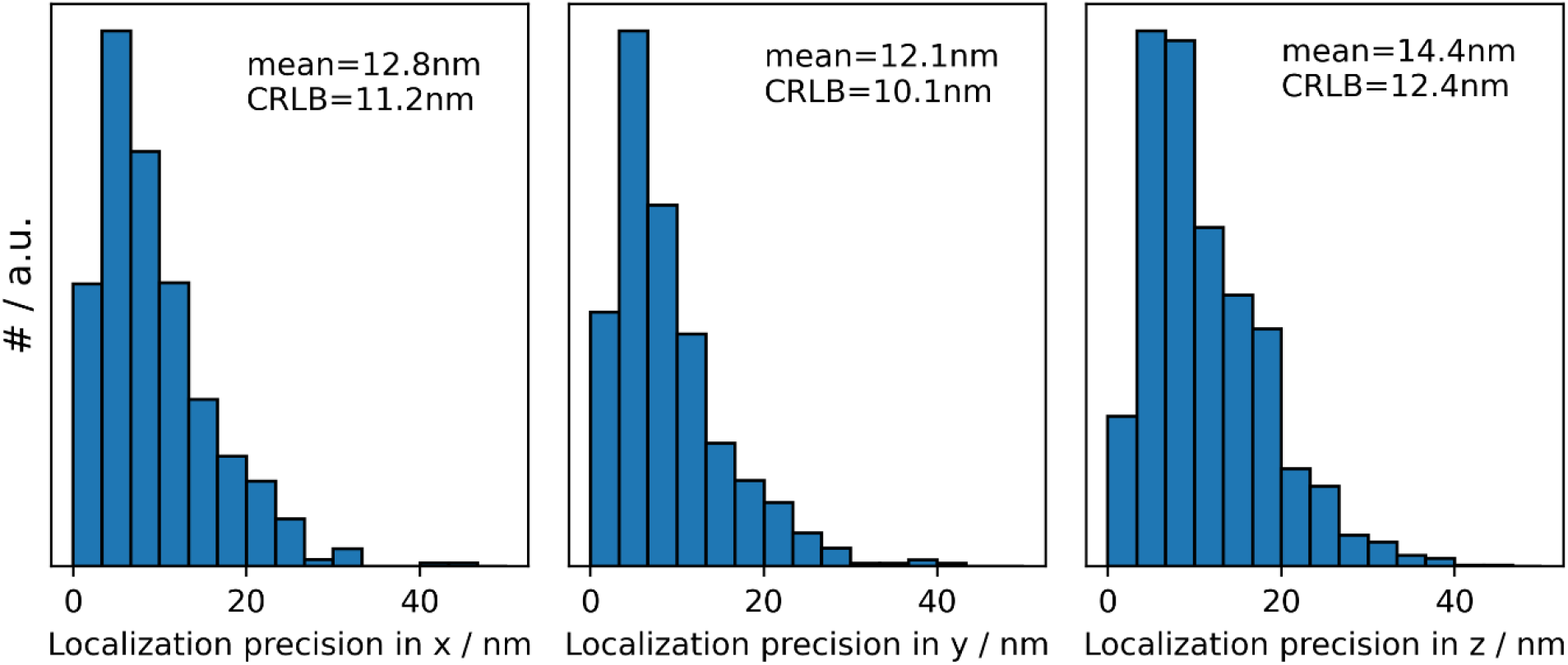
Histogram of localization precision in x, y and z direction. Localization precision was calculated as the standard deviation of localizations of the same molecule appearing in consecutive frames. The mean localization precision and the CRLB are indicated for the different dimensions.

**Supplementary Figure 2:**
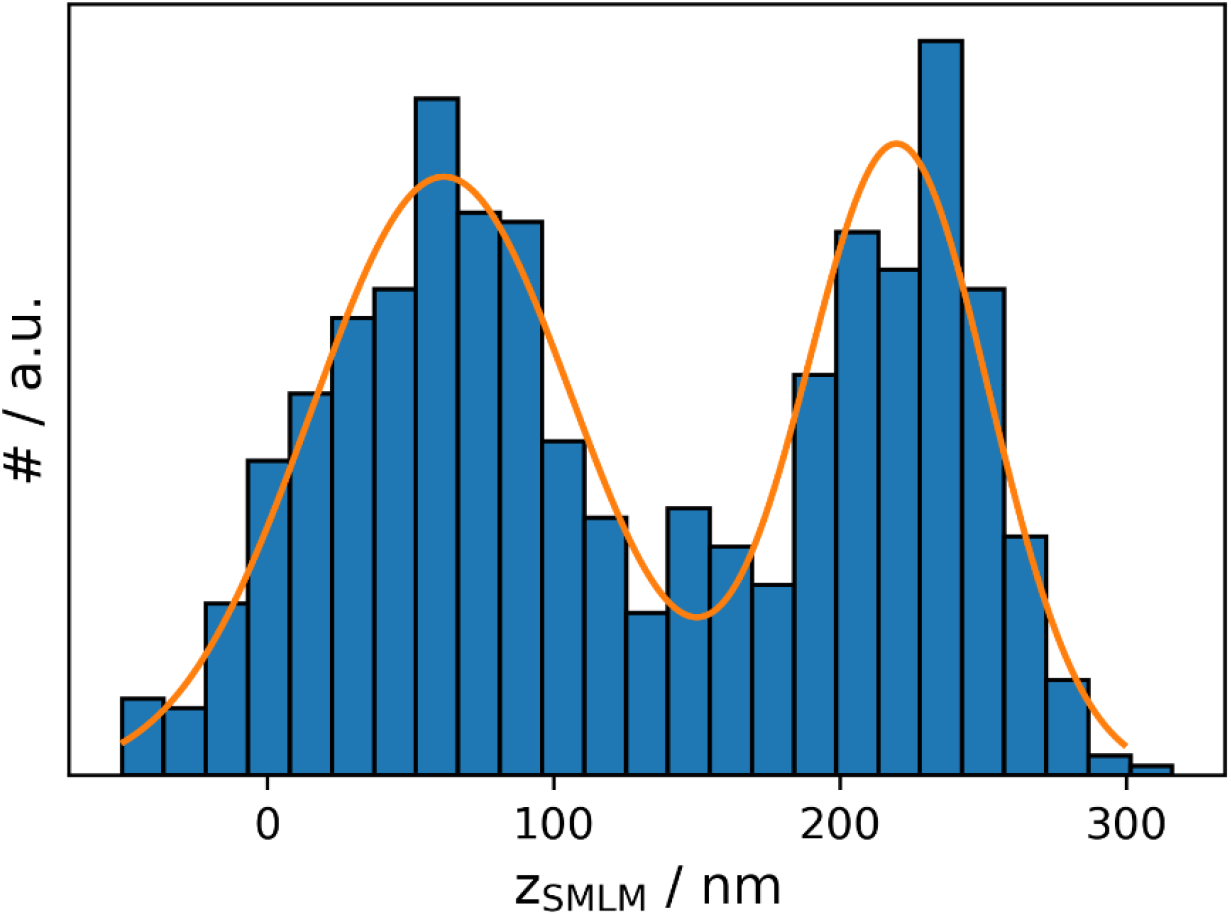
Histogram of z_SMLM_ positions in the area of the lamellipodium shown in **Fig. 2bii**, yielding 2 peaks at a mutual distance of 160 nm.

**Supplementary Figure 3:**
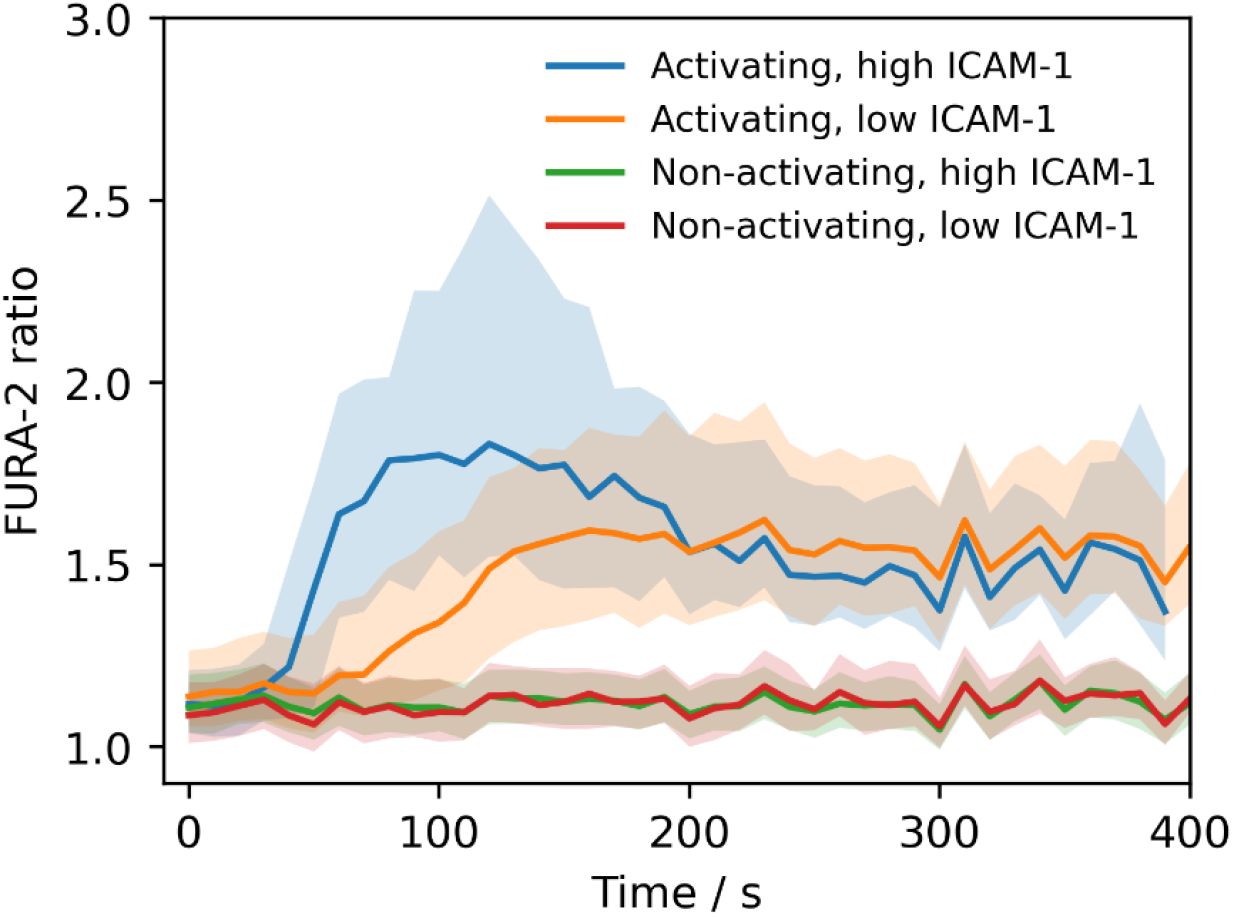
Release of intracellular calcium assessed via the median FURA-2 ratio. Data are shown as a function of time post seeding for T-cells subjected to activating and non-activating conditions while employing the ICAM-1 at indicated densities.

**Supplementary Figure 4:**
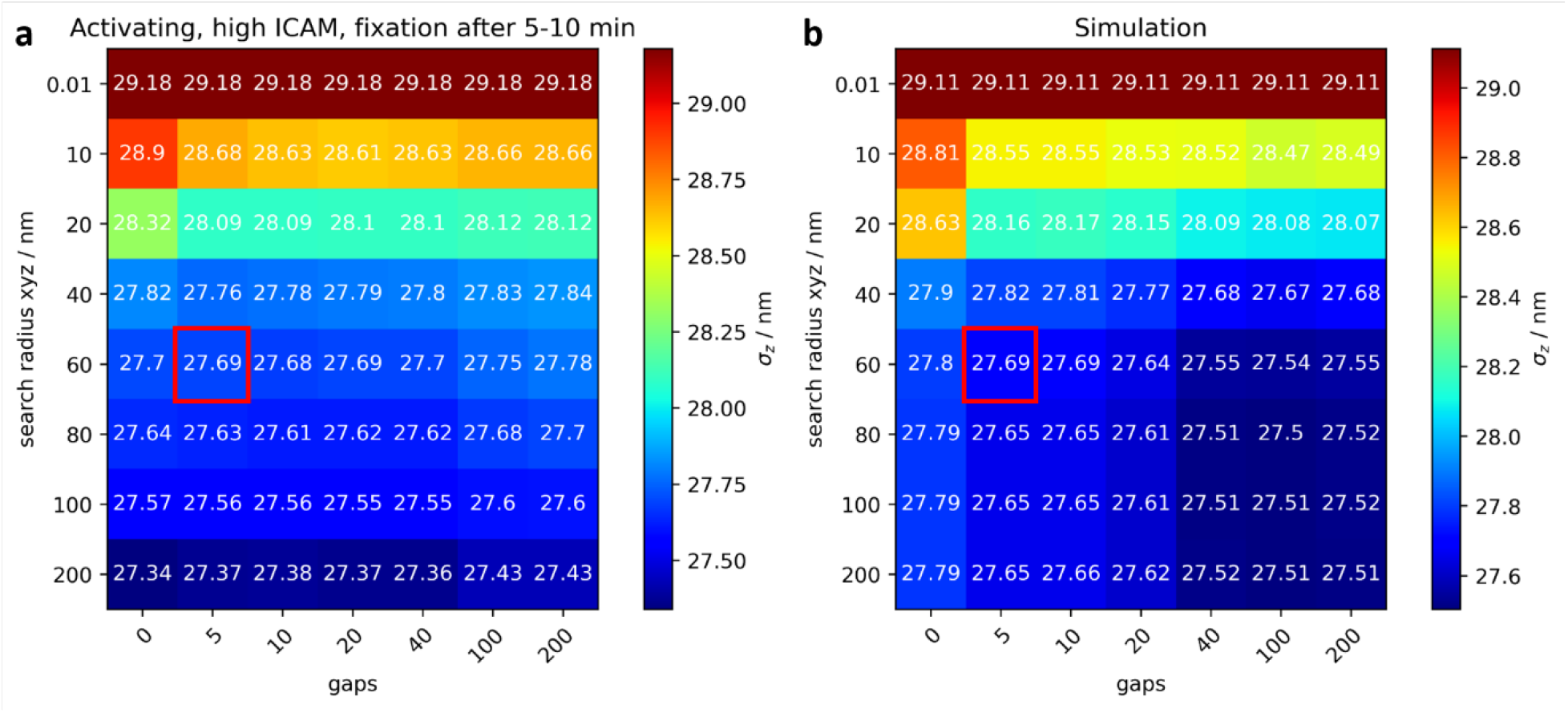
Selecting the appropriate parameters for correction of overcounts. **a)** 3D SMLM experiments were performed on T-cells recorded under activating conditions as described for experiments shown in **Fig. 3**. We determined the standard deviation of z_SMLM_, *σ*_*z*_, of all localization obtained after merging using the indicated three-dimensional search radius and maximally allowed gaps. The chosen value is highlighted by the red square. **b)** For comparison, we analyzed simulated 3D-SMLM data via the same approach. The simulation was based on experimentally derived single molecule blinking statistics and localization precision (see Methods for details). For the simulations we assumed a surface roughness of 26nm.

**Supplementary Figure 5:**
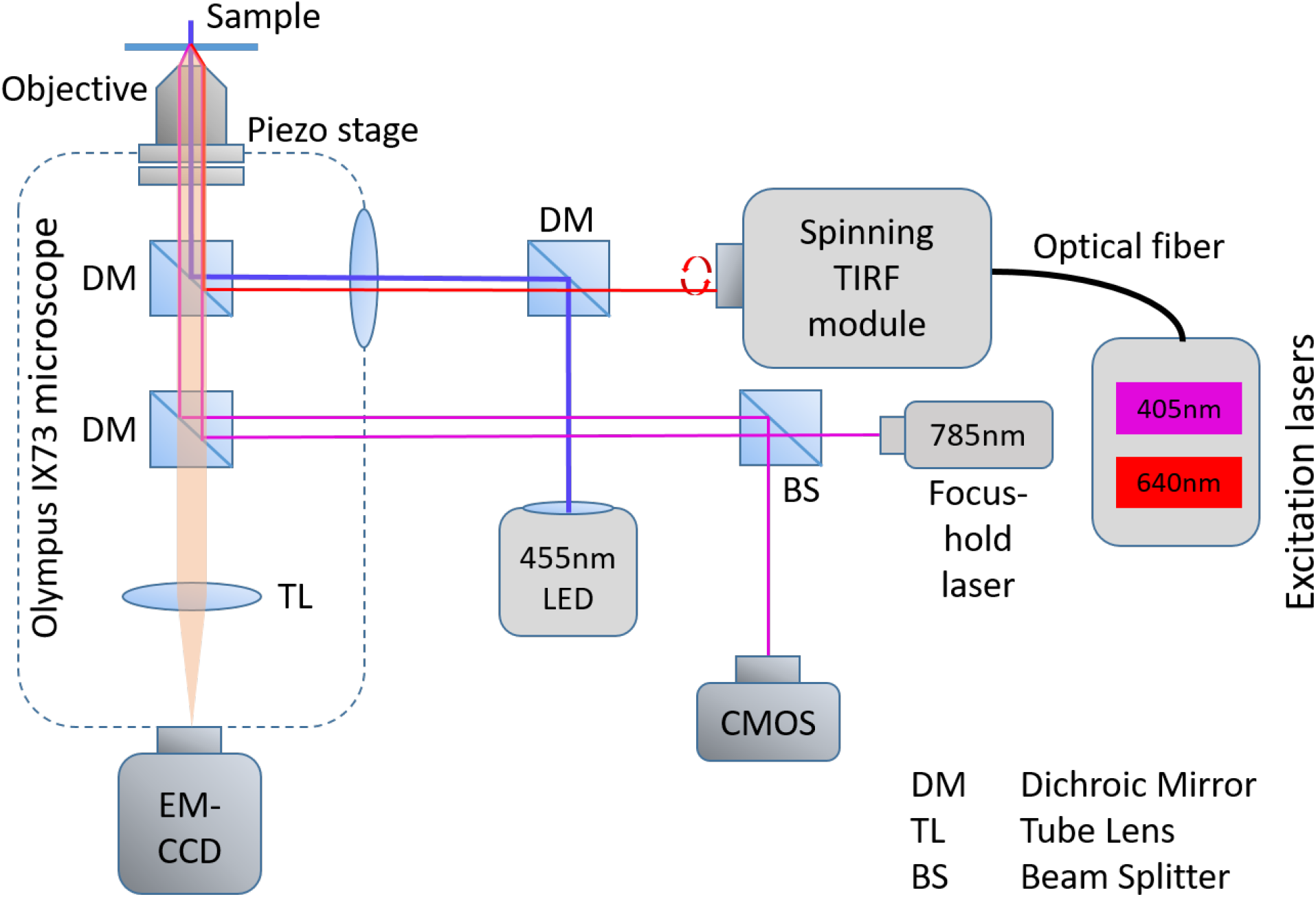
Scheme of the experimental setup.

**Supplementary figure 6:**
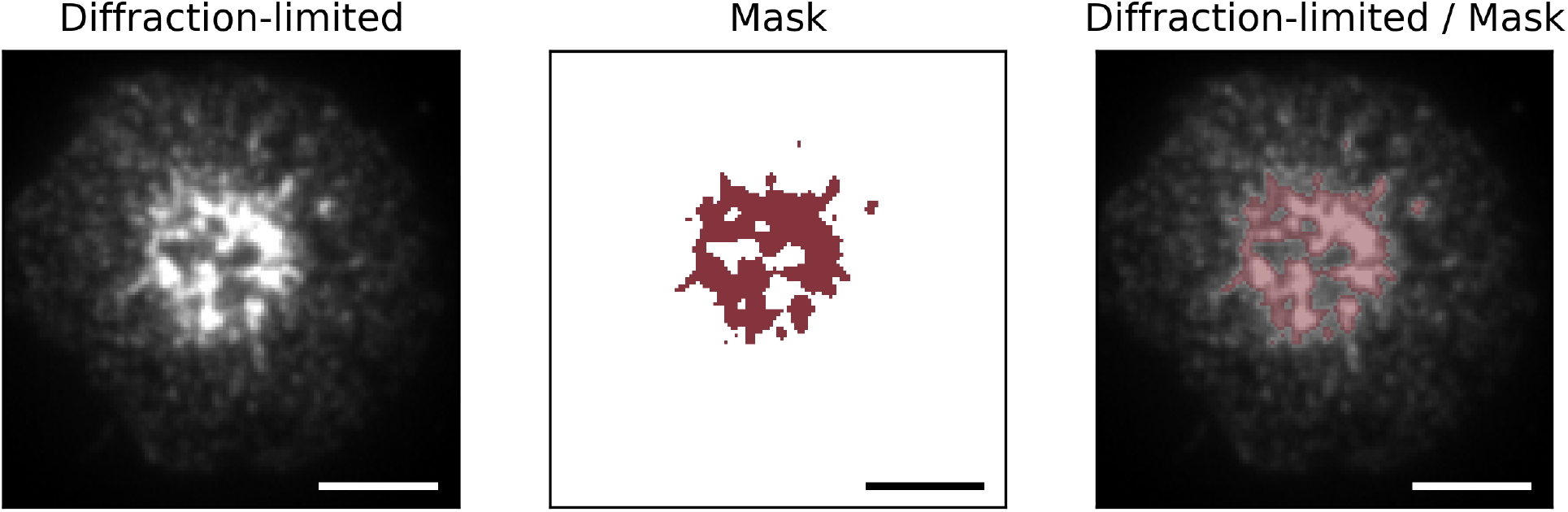
Intensity thresholding was applied to the diffraction-limited images of T-cells to identify TCR microclusters. Pixels with an intensity exceeding the mean intensity of the cell by a factor of 1.5 were considered as part of microclusters (“in”), all other pixels as outside of the microclusters (“out”). Scale bar 5 μm.

## Notes

### Competing Interest Statement

The authors have declared no competing interest.

## REFERENCES

1. Hu J, Lipowsky R, Weikl TR. Binding constants of membrane-anchored receptors and ligands depend strongly on the nanoscale roughness of membranes. Proc Natl Acad Sci U S A 110, 15283–15288 (2013).

2. Dustin ML, et al. Low affinity interaction of human or rat T cell adhesion molecule CD2 with its ligand aligns adhering membranes to achieve high physiological affinity. J Biol Chem 272, 30889–30898 (1997).

3. Cartwright ANR, Griggs J, Davis DM. The immune synapse clears and excludes molecules above a size threshold. Nature Communications 5, 5479 (2014).

4. Davis SJ, van der Merwe PA. The kinetic-segregation model: TCR triggering and beyond. Nat Immunol 7, 803–809 (2006).

5. Grakoui A, et al. The immunological synapse: a molecular machine controlling T cell activation. Science 285, 221–227 (1999).

6. Huppa JB, et al. TCR-peptide-MHC interactions in situ show accelerated kinetics and increased affinity. Nature 463, 963–967 (2010).

7. Groves JT, Dustin ML. Supported planar bilayers in studies on immune cell adhesion and communication. J Immunol Methods 278, 19–32 (2003).

8. Curtis ASG. The Mechanism of Adhesion of Cells to Glass: A Study by Interference Reflection Microscopy. Journal of Cell Biology 20, 199–215 (1964).

9. Limozin L, Sengupta K. Quantitative Reflection Interference Contrast Microscopy (RICM) in Soft Matter and Cell Adhesion. ChemPhysChem 10, 2752–2768 (2009).

10. Smith A-S, Sengupta K, Goennenwein S, Seifert U, Sackmann E. Force-induced growth of adhesion domains is controlled by receptor mobility. Proc Natl Acad Sci U S A 105, 6906–6911 (2008).

11. Balagopalan L, Sherman E, Barr VA, Samelson LE. Imaging techniques for assaying lymphocyte activation in action. Nat Rev Immunol 11, 21–33 (2011).

12. Cai E, et al. Visualizing dynamic microvillar search and stabilization during ligand detection by T cells. Science 356, eaal3118 (2017).

13. Axelrod D, Burghardt TP, Thompson NL. Total internal reflection fluorescence. Annu Rev Biophys Bioeng 13, 247–268 (1984).

14. Hashimoto-Tane A, Saito T. Dynamic Regulation of TCR–Microclusters and the Microsynapse for T Cell Activation. Frontiers in Immunology 7, 255 (2016).

15. Yokosuka T, et al. Newly generated T cell receptor microclusters initiate and sustain T cell activation by recruitment of Zap70 and SLP-76. Nat Immunol 6, 1253–1262 (2005).

16. Varma R, Campi G, Yokosuka T, Saito T, Dustin ML. T cell receptor-proximal signals are sustained in peripheral microclusters and terminated in the central supramolecular activation cluster. Immunity 25, 117–127 (2006).

17. Rossy J, Owen DM, Williamson DJ, Yang Z, Gaus K. Conformational states of the kinase Lck regulate clustering in early T cell signaling. Nat Immunol 14, 82–89 (2013).

18. Pageon SV, et al. Functional role of T-cell receptor nanoclusters in signal initiation and antigen discrimination. Proceedings of the National Academy of Sciences 113, E5454–5463 (2016).

19. Rossboth B, et al. TCRs are randomly distributed on the plasma membrane of resting antigen-experienced T cells. Nature Immunology 19, 821–827 (2018).

20. Simoncelli S, et al. Multi-color Molecular Visualization of Signaling Proteins Reveals How C-Terminal Src Kinase Nanoclusters Regulate T Cell Receptor Activation. Cell Reports 33, 108523 (2020).

21. Jung Y, et al. Three-dimensional localization of T-cell receptors in relation to microvilli using a combination of superresolution microscopies. Proceedings of the National Academy of Sciences 113, E5916–E5924 (2016).

22. Jung Y, Wen L, Altman A, Ley K. CD45 pre-exclusion from the tips of microvilli establishes a phosphatase-free zone for early TCR triggering. bioRxiv, 2020.2005.2021.109074 (2020).

23. Sauer M, Heilemann M. Single-Molecule Localization Microscopy in Eukaryotes. Chemical Reviews, (2017).

24. Hajj B, El Beheiry M, Izeddin I, Darzacq X, Dahan M. Accessing the third dimension in localization-based super-resolution microscopy. Physical Chemistry Chemical Physics 16, 16340–16348 (2014).

25. Oheim M, Salomon A, Brunstein M. Supercritical Angle Fluorescence Microscopy and Spectroscopy. Biophysical Journal 118, 2339–2348 (2020).

26. Bourg N, et al. Direct optical nanoscopy with axially localized detection. Nat Photon 9, 587–593 (2015).

27. Zelger P, Bodner L, Velas L, Schütz GJ, Jesacher A. Defocused imaging exploits supercritical-angle fluorescence emission for precise axial single molecule localization microscopy. Biomed Opt Express 11, 775–790 (2020).

28. Ober RJ, Ram S, Ward ES. Localization accuracy in single-molecule microscopy. Biophys J 86, 1185–1200 (2004).

29. Hellmeier J, et al. DNA origami demonstrate the unique stimulatory power of single pMHCs as T cell antigens. Proceedings of the National Academy of Sciences 118, e2016857118 (2021).

30. Abraham VC, Krishnamurthi V, Taylor DL, Lanni F. The Actin-Based Nanomachine at the Leading Edge of Migrating Cells. Biophysical Journal 77, 1721–1732 (1999).

31. Hashimoto-Tane A, et al. Dynein-Driven Transport of T Cell Receptor Microclusters Regulates Immune Synapse Formation and T Cell Activation. Immunity 34, 919–931 (2011).

32. Onnis A, Baldari CT. Orchestration of Immunological Synapse Assembly by Vesicular Trafficking. Frontiers in Cell and Developmental Biology 7, (2019).

33. Choudhuri K, et al. Polarized release of T-cell-receptor-enriched microvesicles at the immunological synapse. Nature 507, 118–123 (2014).

34. Chang VT, et al. Initiation of T cell signaling by CD45 segregation at ‘close contacts’. Nat Immunol 17, 574–582 (2016).

35. Santos AM, et al. Capturing resting T cells: the perils of PLL. Nature Immunology, (2018).

36. Choudhuri K, Wiseman D, Brown MH, Gould K, van der Merwe PA. T-cell receptor triggering is critically dependent on the dimensions of its peptide-MHC ligand. Nature 436, 578–582 (2005).

37. Taylor MJ, Husain K, Gartner ZJ, Mayor S, Vale RD. A DNA-Based T Cell Receptor Reveals a Role for Receptor Clustering in Ligand Discrimination. Cell 169, 108-119.e120 (2017).

38. Fernandes RA, et al. A cell topography-based mechanism for ligand discrimination by the T cell receptor. Proceedings of the National Academy of Sciences 116, 14002–14010 (2019).

39. Al-Aghbar MA, Chu Y-S, Chen B-M, Roffler SR. High-Affinity Ligands Can Trigger T Cell Receptor Signaling Without CD45 Segregation. Frontiers in Immunology 9, (2018).

40. Courtney AH, et al. CD45 functions as a signaling gatekeeper in T cells. Science Signaling 12, eaaw8151 (2019).

41. Cai H, et al. Full control of ligand positioning reveals spatial thresholds for T cell receptor triggering. Nature Nanotechnology 13, 610–617 (2018).

42. Yin Y, Wang XX, Mariuzza RA. Crystal structure of a complete ternary complex of T-cell receptor, peptide-MHC, and CD4. Proc Natl Acad Sci U S A 109, 5405–5410 (2012).

43. Jung Y, Wen L, Altman A, Ley K. CD45 pre-exclusion from the tips of T cell microvilli prior to antigen recognition. Nature Communications 12, 3872 (2021).

44. Lanz A-L, et al. Allosteric activation of T cell antigen receptor signaling by quaternary structure relaxation. Cell Reports 36, (2021).

45. Huppa JB, Gleimer M, Sumen C, Davis MM. Continuous T cell receptor signaling required for synapse maintenance and full effector potential. Nat Immunol 4, 749–755 (2003).

46. Zelger P, Kaser K, Rossboth B, Velas L, Schütz GJ, Jesacher A. Three-dimensional localization microscopy using deep learning. Optics Express 26, 33166–33179 (2018).

47. Allan DB, Caswell T, Keim NC, van der Wel CM, Verweij RW. trackpy: Trackpy v0.5.0. Zenodo, http://doi.org/10.5281/zenodo.4682814 (2021).

