## Supplementary Image Gallery for "3D single molecule localization microscopy reveals the topography of the immunological synapse at isotropic precision below 15 nm"

**Supplementary gallery**

**Figure 1a: Activating conditions, high ICAM-1 density, fixation: 5-10 min post seeding**

**Correlative 3D-SMLM, IRM, and diffraction-limited TIR microscopy of the immunological synapse.** T cells were activated on an SLB functionalized with I-$E^k$/MCC, B7-1 and high density of ICAM-1, and fixed 5-10 minutes post seeding. The T cell was imaged with IRM and fluorescence microscopy: (i) Diffraction-limited TIR image of the T cell. (ii) Reconstruction of the diffraction-limited image by convolving the 3D-SMLM image with the corresponding psf. (iii) Overlay of the diffraction-limited TIR image with the 3D-SMLM image. Color-code indicates distance to the coverslip $z_{SMLM}$. (iv) IRM image. (v) Overlay of the IRM image with the diffraction-limited image. (vi) Overlay of the IRM and the 3D-SMLM image. Bottom row images were generated by calculating the pixel-wise average of the 3D-SMLM images (pixel size of 146nm is consistent with diffraction-limited image) according to pixelated mean single molecule (SM) intensity (vii), pixelated number of localizations (viii) and pixelated mean $z_{SMLM}$ (ix). Scale bars 5 µm.

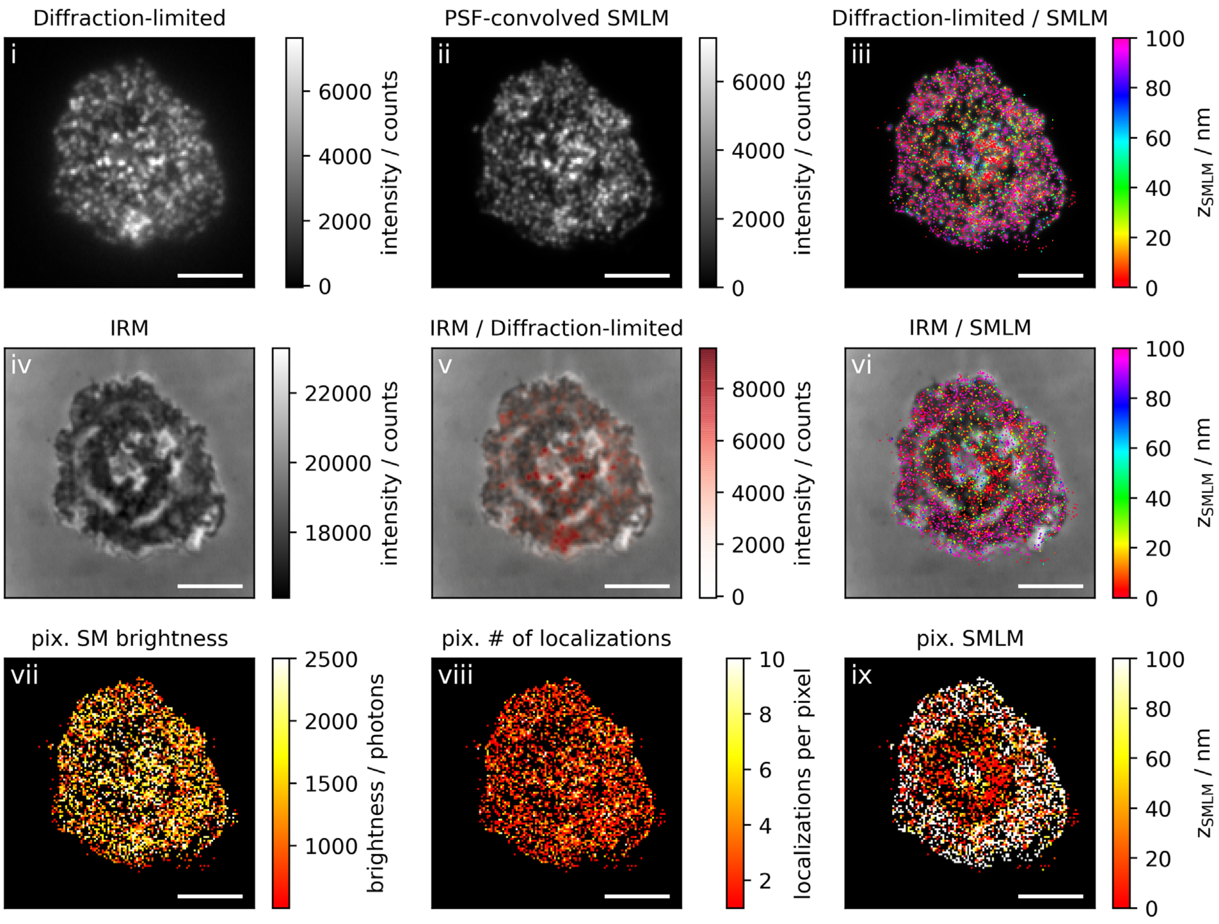

Diffraction-limited
i
intensity / counts
0
2000
4000
6000
PSF-convolved SMLM
ii
Diffraction-limited / SMLM
iii
z_SMLM / nm
0
20
40
60
80
100
IRM
iv
18000
20000
22000
IRM / Diffraction-limited
v
8000
IRM / SMLM
vi
pix. SM brightness
vii
brightness / photons
1000
1500
2000
2500
pix. # of localizations
viii
localizations per pixel
2
4
6
8
10
pix. SMLM
ix

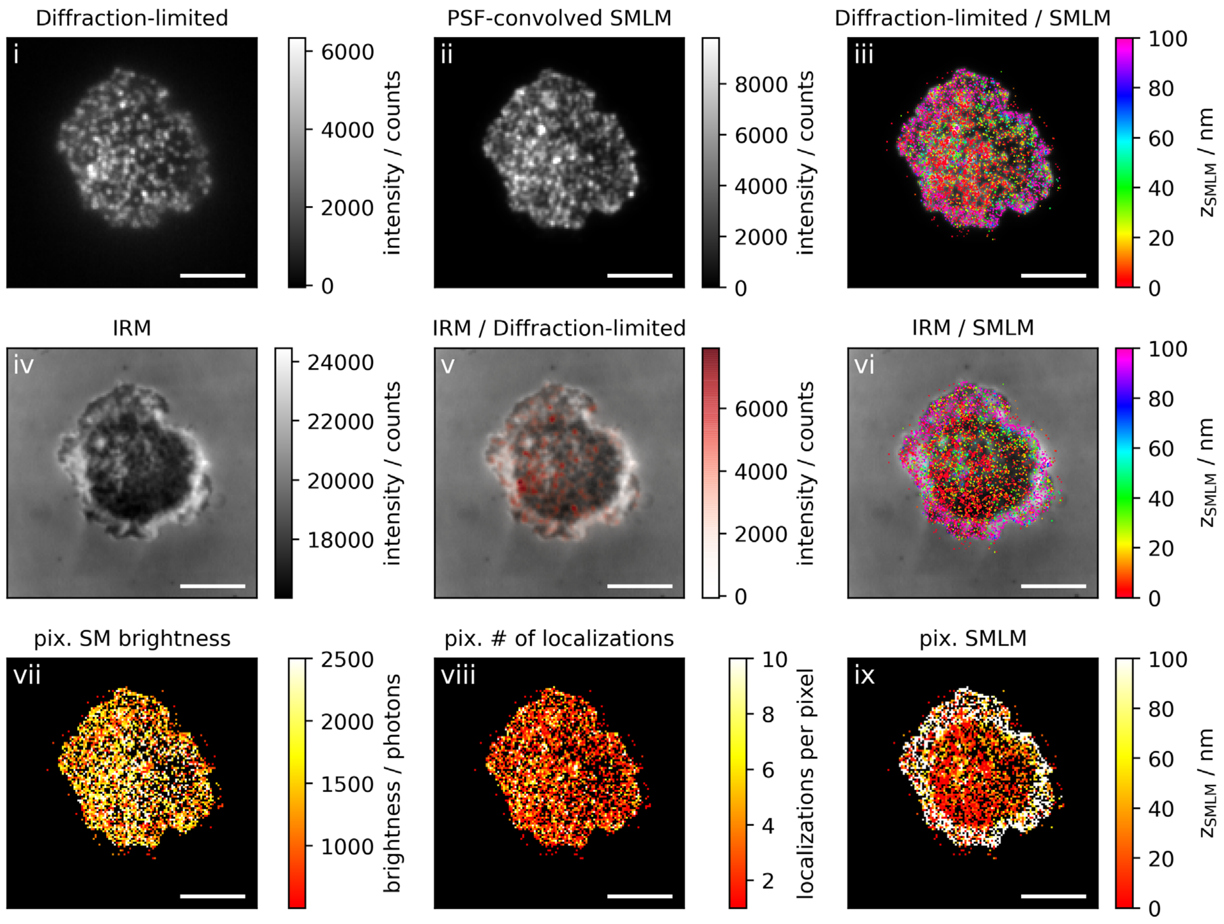

Diffraction-limited
i
6000
4000
2000
0
intensity / counts
PSF-convolved SMLM
ii
8000
6000
4000
2000
0
intensity / counts
Diffraction-limited / SMLM
iii
100
80
60
40
20
0
$z_{SMLM}$ / nm
IRM
iv
24000
22000
20000
18000
intensity / counts
IRM / Diffraction-limited
v
6000
4000
2000
0
intensity / counts
IRM / SMLM
vi
100
80
60
40
20
0
$z_{SMLM}$ / nm
pix. SM brightness
vii
2500
2000
1500
1000
brightness / photons
pix. # of localizations
viii
10
8
6
4
2
localizations per pixel
pix. SMLM
ix
100
80
60
40
20
0
$z_{SMLM}$ / nm

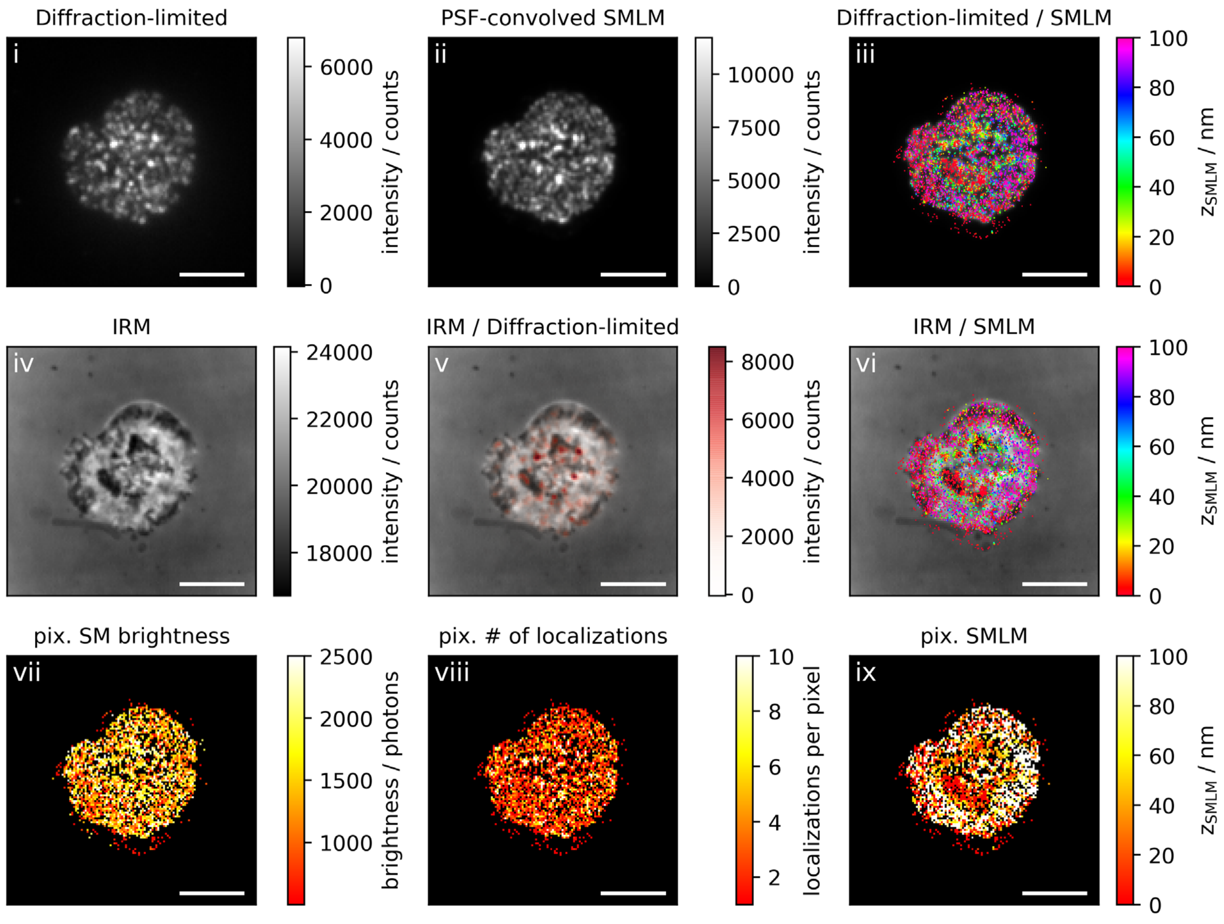

Diffraction-limited
i
6000
4000
2000
0
intensity / counts
PSF-convolved SMLM
ii
10000
7500
5000
2500
0
intensity / counts
Diffraction-limited / SMLM
iii
100
80
60
40
20
0
z_SMLM / nm
IRM
iv
24000
22000
20000
18000
intensity / counts
IRM / Diffraction-limited
v
8000
6000
4000
2000
0
intensity / counts
IRM / SMLM
vi
100
80
60
40
20
0
z_SMLM / nm
pix. SM brightness
vii
2500
2000
1500
1000
brightness / photons
pix. # of localizations
viii
10
8
6
4
2
localizations per pixel
pix. SMLM
ix
100
80
60
40
20
0
z_SMLM / nm

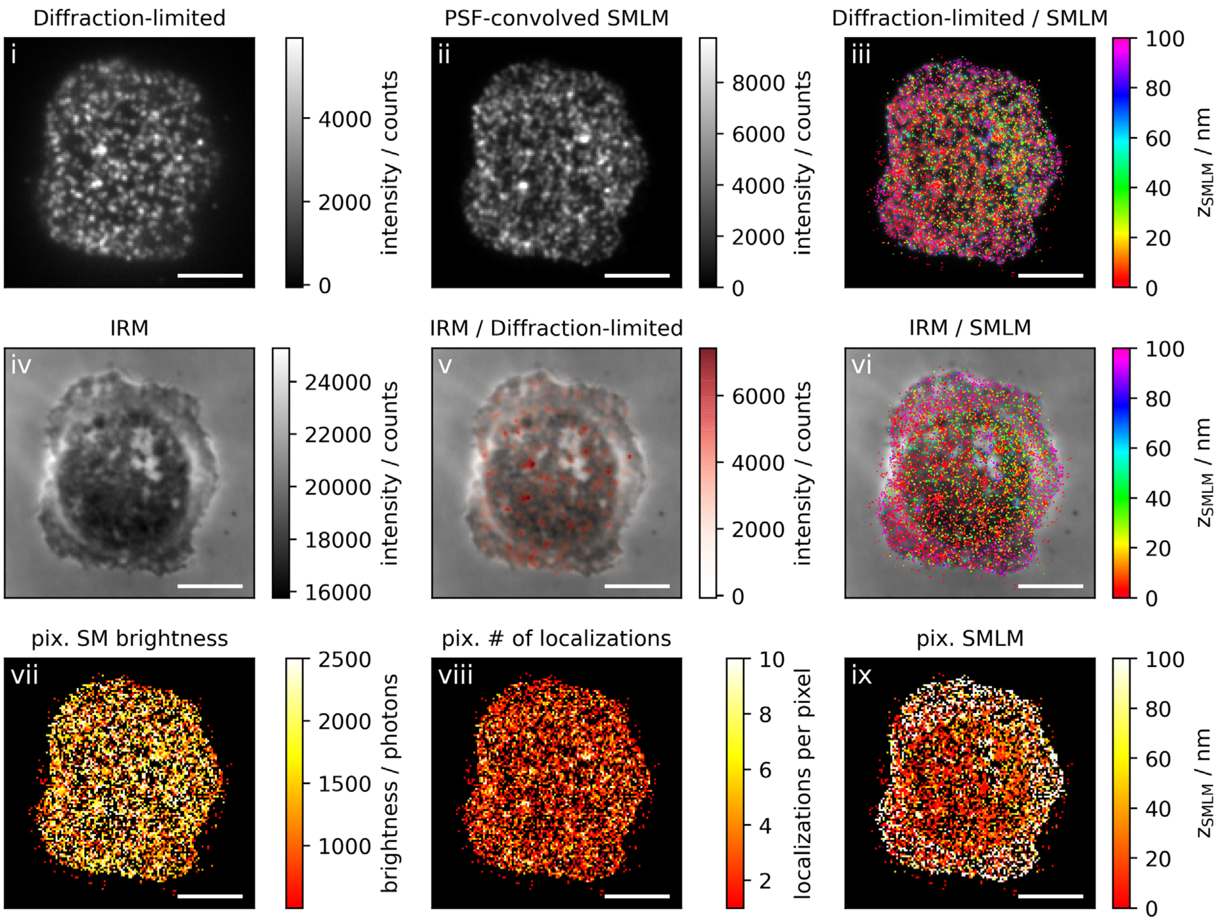
Diffraction-limited
PSF-convolved SMLM
Diffraction-limited / SMLM
IRM
IRM / Diffraction-limited
IRM / SMLM
pix. SM brightness
pix. # of localizations
pix. SMLM
i
ii
iii
iv
v
vi
vii
viii
ix
intensity / counts
$z_{SMLM}$ / nm
brightness / photons
localizations per pixel

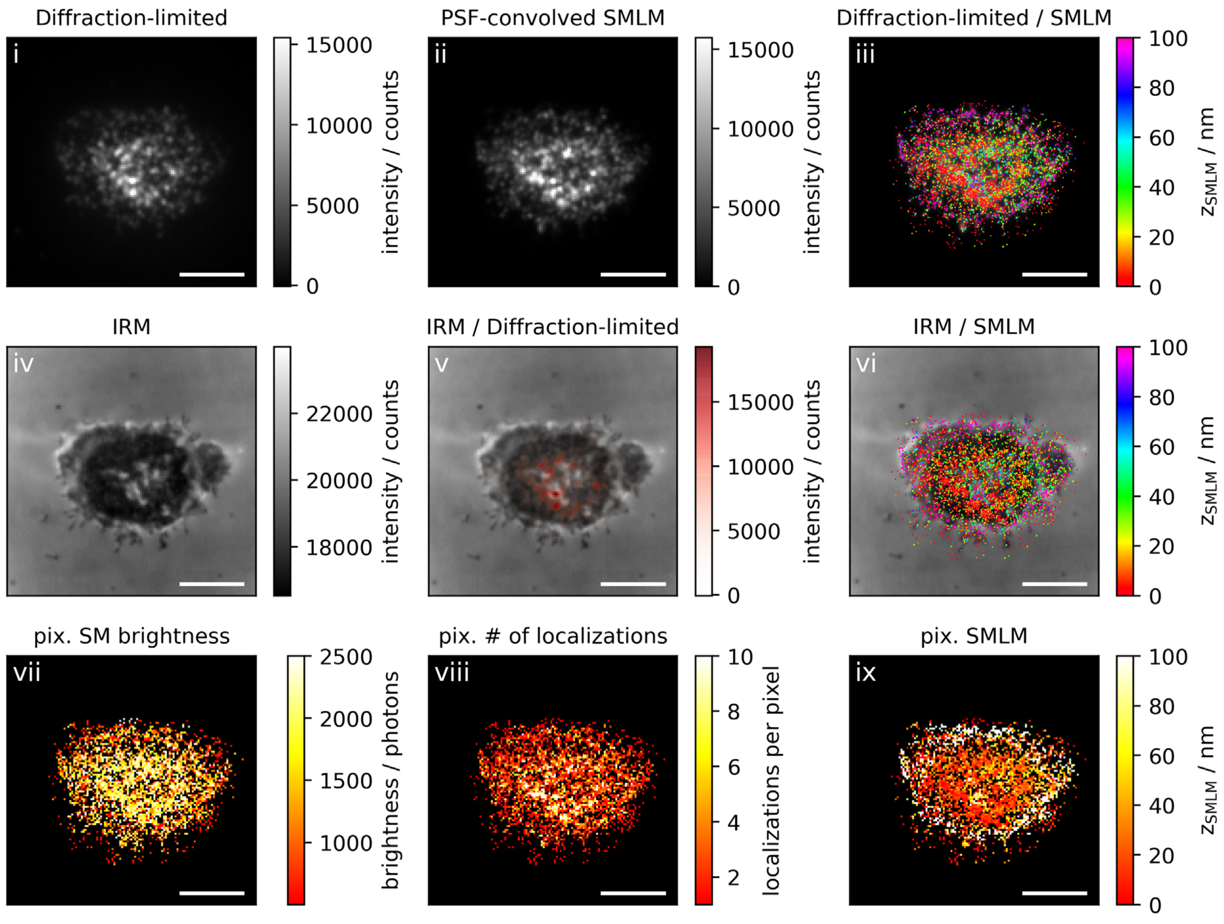
Diffraction-limited
i
15000
10000
5000
0
intensity / counts
PSF-convolved SMLM
ii
15000
10000
5000
0
intensity / counts
Diffraction-limited / SMLM
iii
100
80
60
40
20
0
$z_{SMLM}$ / nm
IRM
iv
22000
20000
18000
intensity / counts
IRM / Diffraction-limited
v
15000
10000
5000
0
intensity / counts
IRM / SMLM
vi
100
80
60
40
20
0
$z_{SMLM}$ / nm
pix. SM brightness
vii
2500
2000
1500
1000
brightness / photons
pix. # of localizations
viii
10
8
6
4
2
localizations per pixel
pix. SMLM
ix
100
80
60
40
20
0
$z_{SMLM}$ / nm

**Figure 1b: Activating conditions, high ICAM-1 density, fixation: 10 min post seeding**

**Correlative 3D-SMLM, IRM, and diffraction-limited TIR microscopy of the immunological synapse.** T cells were activated on an SLB functionalized with I-$E^k$/MCC, B7-1 and high density of ICAM-1, and fixed 10 minutes post seeding. The T cell was imaged with IRM and fluorescence microscopy: (i) Diffraction-limited TIR image of the T cell. (ii) Reconstruction of the diffraction-limited image by convolving the 3D-SMLM image with the corresponding psf. (iii) Overlay of the diffraction-limited TIR image with the 3D-SMLM image. Color-code indicates distance to the coverslip $z_{SMLM}$. (iv) IRM image. (v) Overlay of the IRM image with the diffraction-limited image. (vi) Overlay of the IRM and the 3D-SMLM image. Bottom row images were generated by calculating the pixel-wise average of the 3D-SMLM images (pixel size of 146nm is consistent with diffraction-limited image) according to pixelated mean single molecule (SM) intensity (vii), pixelated number of localizations (viii) and pixelated mean $z_{SMLM}$ (ix). Scale bars 5 µm.

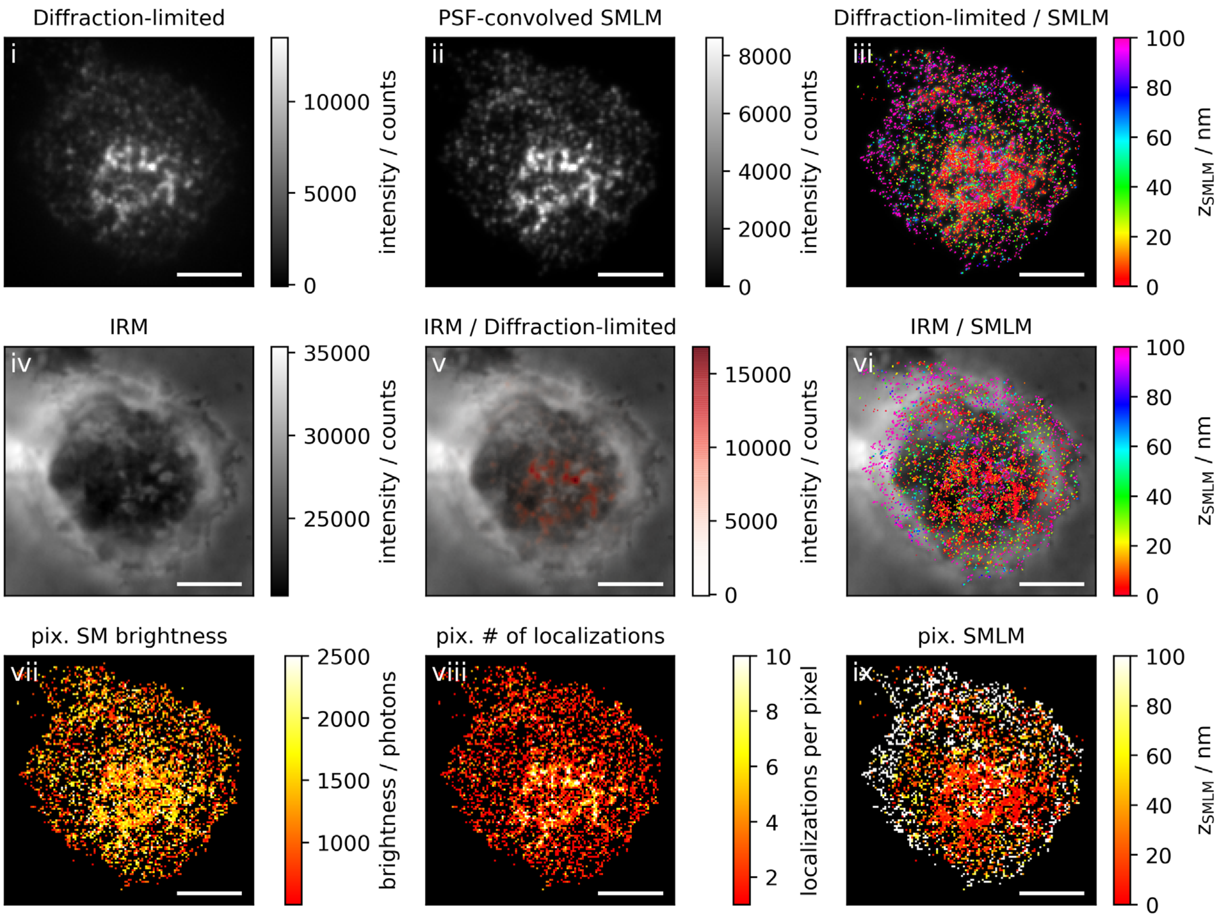

Diffraction-limited
i
intensity / counts
PSF-convolved SMLM
ii
intensity / counts
Diffraction-limited / SMLM
iii
$z_{SMLM}$ / nm
IRM
iv
intensity / counts
IRM / Diffraction-limited
v
intensity / counts
IRM / SMLM
vi
$z_{SMLM}$ / nm
pix. SM brightness
vii
brightness / photons
pix. # of localizations
viii
localizations per pixel
pix. SMLM
ix
$z_{SMLM}$ / nm

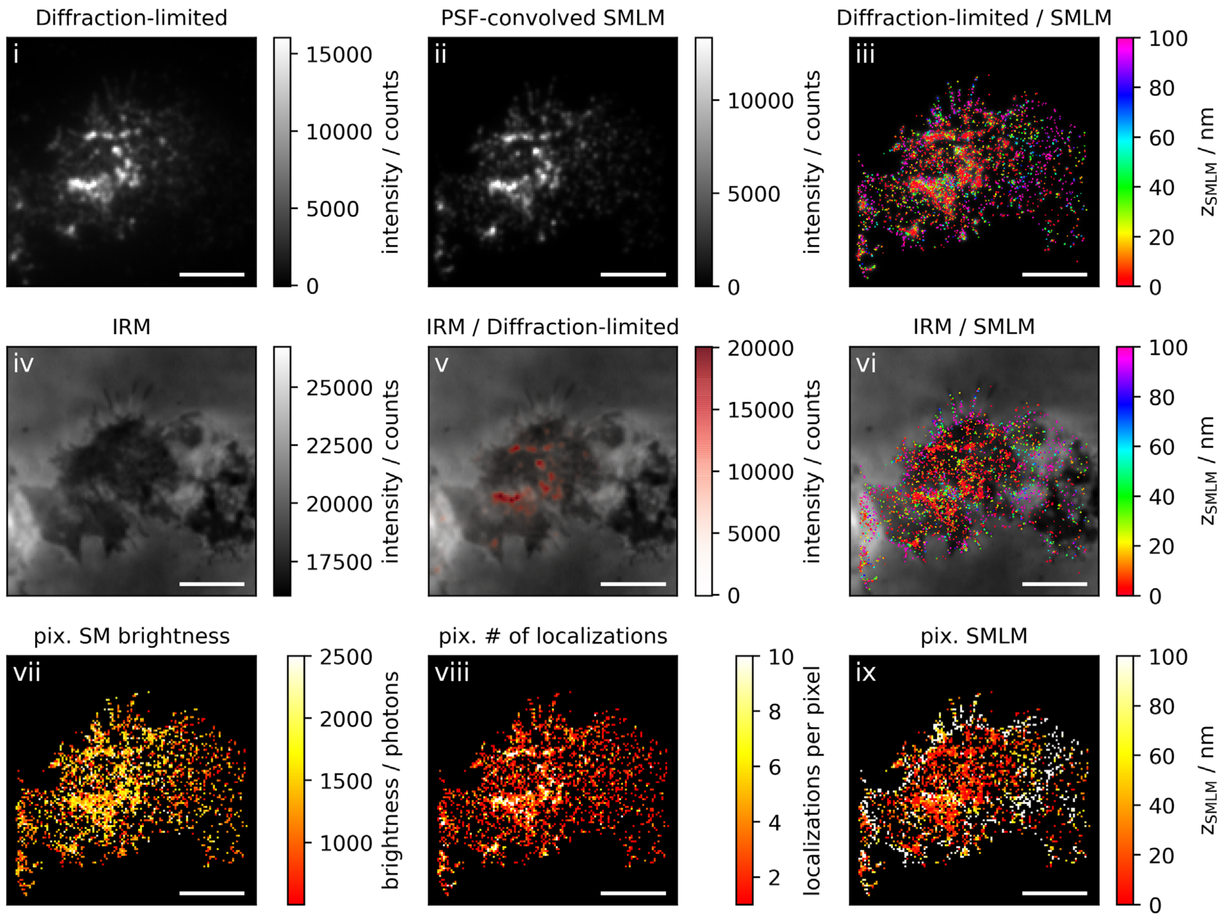
Diffraction-limited
i
15000
10000
5000
0
intensity / counts
PSF-convolved SMLM
ii
10000
5000
0
intensity / counts
Diffraction-limited / SMLM
iii
100
80
60
40
20
0
z_SMLM / nm
IRM
iv
25000
22500
20000
17500
intensity / counts
IRM / Diffraction-limited
v
20000
15000
10000
5000
0
intensity / counts
IRM / SMLM
vi
100
80
60
40
20
0
z_SMLM / nm
pix. SM brightness
vii
2500
2000
1500
1000
brightness / photons
pix. # of localizations
viii
10
8
6
4
2
localizations per pixel
pix. SMLM
ix
100
80
60
40
20
0
z_SMLM / nm

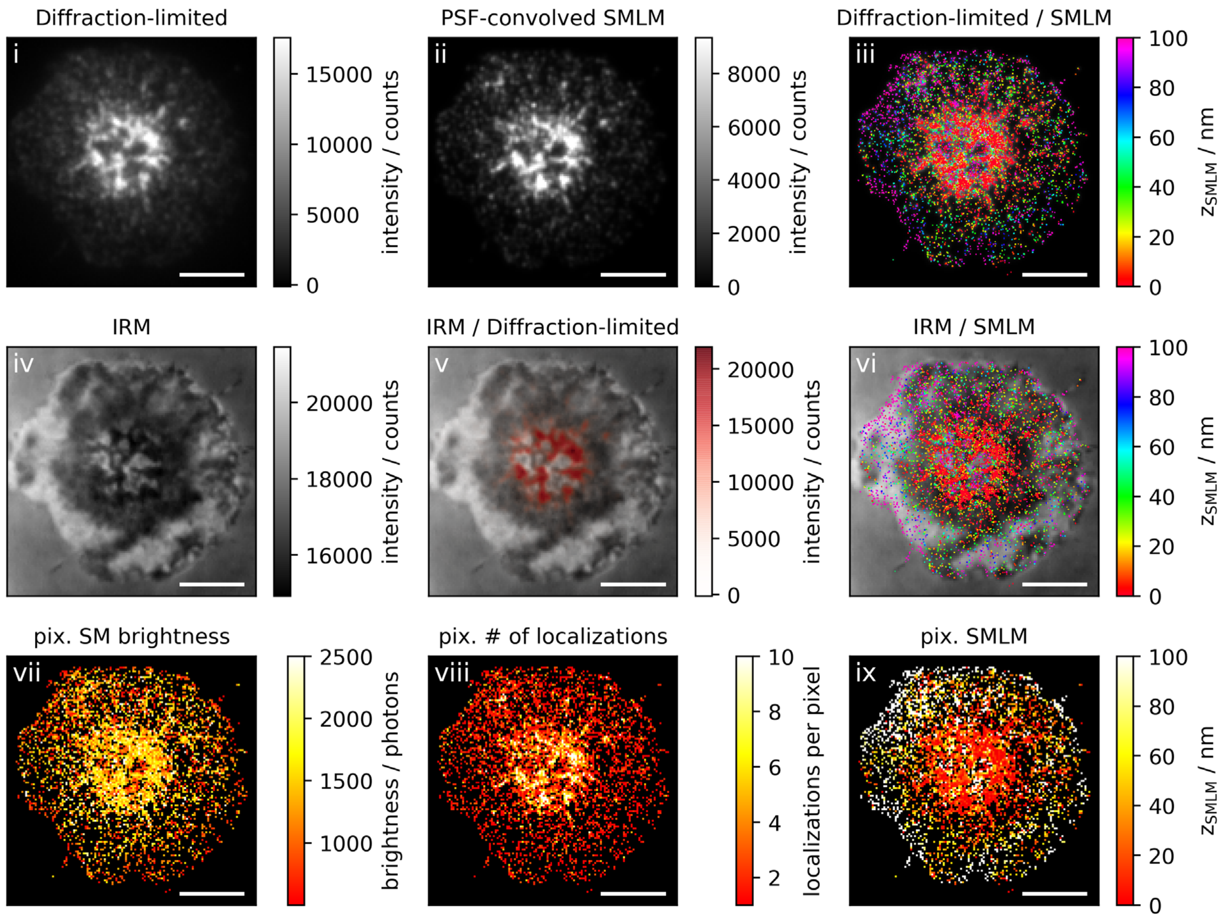

Diffraction-limited
PSF-convolved SMLM
Diffraction-limited / SMLM
IRM
IRM / Diffraction-limited
IRM / SMLM
pix. SM brightness
pix. # of localizations
pix. SMLM
i
ii
iii
iv
v
vi
vii
viii
ix
intensity / counts
$z_{SMLM}$ / nm
brightness / photons
localizations per pixel
0
5000
10000
15000
2000
4000
6000
8000
20
40
60
80
100
16000
18000
20000
1000
1500
2000
2500
2
4
6
8
10

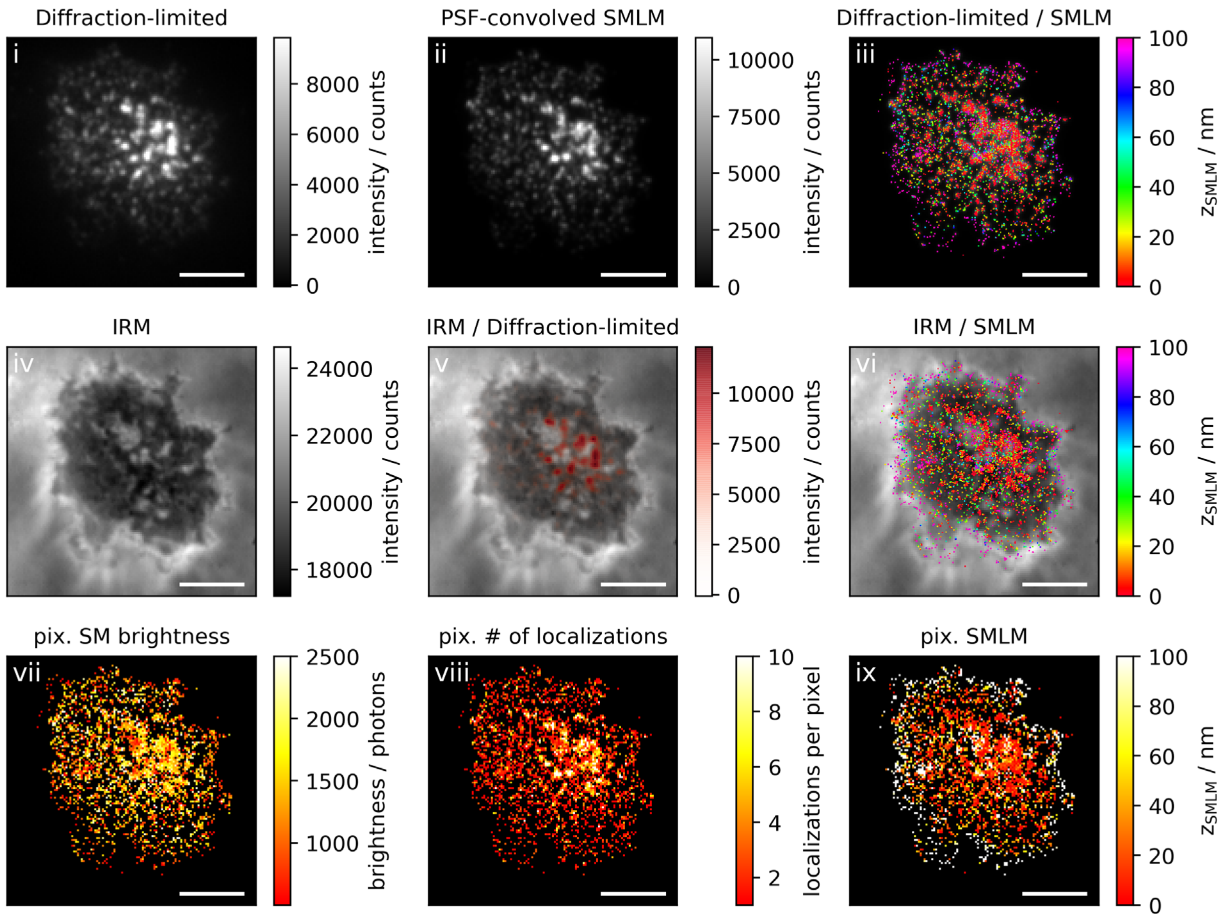
Diffraction-limited
i
intensity / counts
PSF-convolved SMLM
ii
intensity / counts
Diffraction-limited / SMLM
iii
$z_{SMLM}$ / nm
IRM
iv
intensity / counts
IRM / Diffraction-limited
v
intensity / counts
IRM / SMLM
vi
$z_{SMLM}$ / nm
pix. SM brightness
vii
brightness / photons
pix. # of localizations
viii
localizations per pixel
pix. SMLM
ix
$z_{SMLM}$ / nm

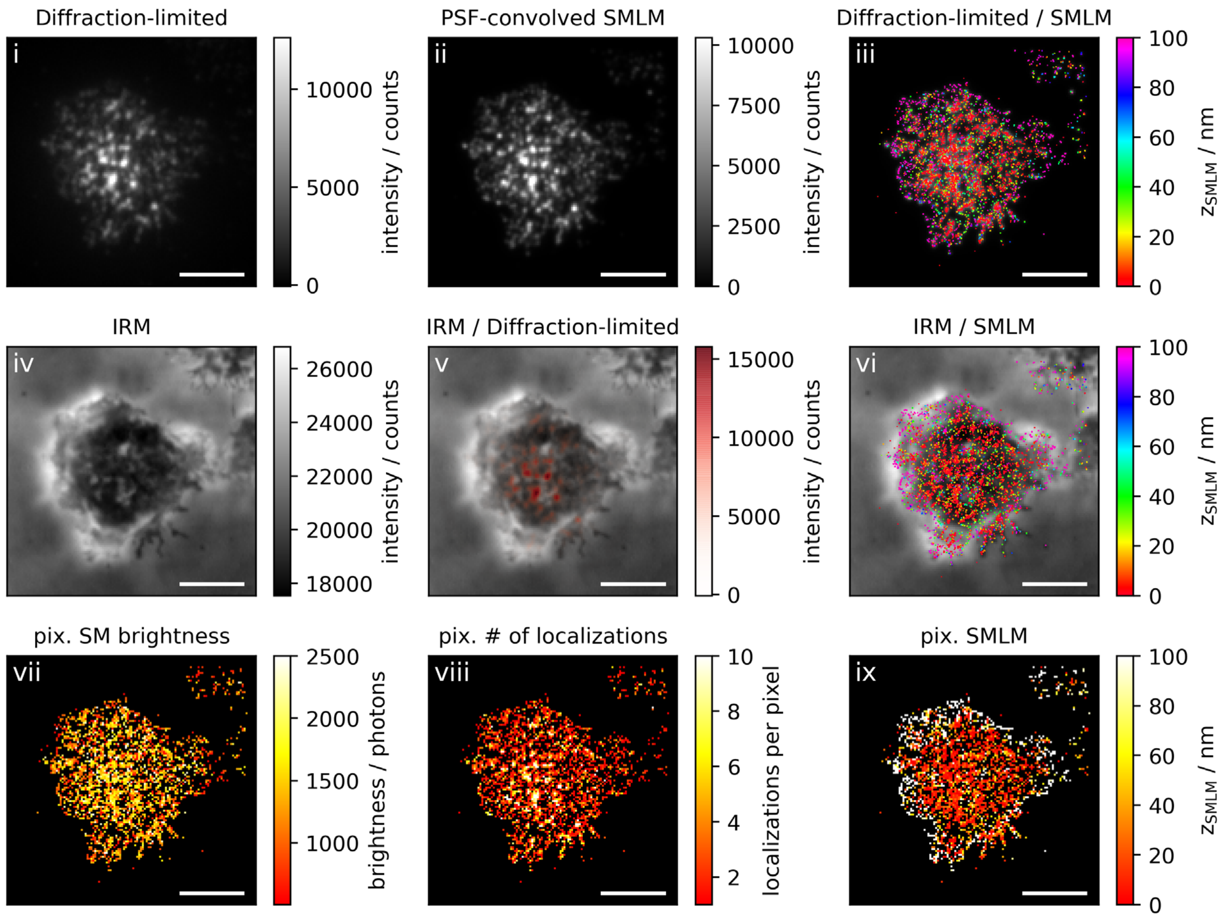

Diffraction-limited
i
intensity / counts
PSF-convolved SMLM
ii
intensity / counts
Diffraction-limited / SMLM
iii
$z_{SMLM}$ / nm
IRM
iv
intensity / counts
IRM / Diffraction-limited
v
intensity / counts
IRM / SMLM
vi
$z_{SMLM}$ / nm
pix. SM brightness
vii
brightness / photons
pix. # of localizations
viii
localizations per pixel
pix. SMLM
ix
$z_{SMLM}$ / nm

**Figure 1c: Activating conditions, high ICAM-1 density, fixation: 10-15 min post seeding**

**Correlative 3D-SMLM, IRM, and diffraction-limited TIR microscopy of the immunological synapse.** T cells were activated on an SLB functionalized with I-$E^k$/MCC, B7-1 and high density of ICAM-1, and fixed 10-15 minutes post seeding. The T cell was imaged with IRM and fluorescence microscopy: (i) Diffraction-limited TIR image of the T cell. (ii) Reconstruction of the diffraction-limited image by convolving the 3D-SMLM image with the corresponding psf. (iii) Overlay of the diffraction-limited TIR image with the 3D-SMLM image. Color-code indicates distance to the coverslip $z_{SMLM}$. (iv) IRM image. (v) Overlay of the IRM image with the diffraction-limited image. (vi) Overlay of the IRM and the 3D-SMLM image. Bottom row images were generated by calculating the pixel-wise average of the 3D-SMLM images (pixel size of 146nm is consistent with diffraction-limited image) according to pixelated mean single molecule (SM) intensity (vii), pixelated number of localizations (viii) and pixelated mean $z_{SMLM}$ (ix). Scale bars 5 µm.

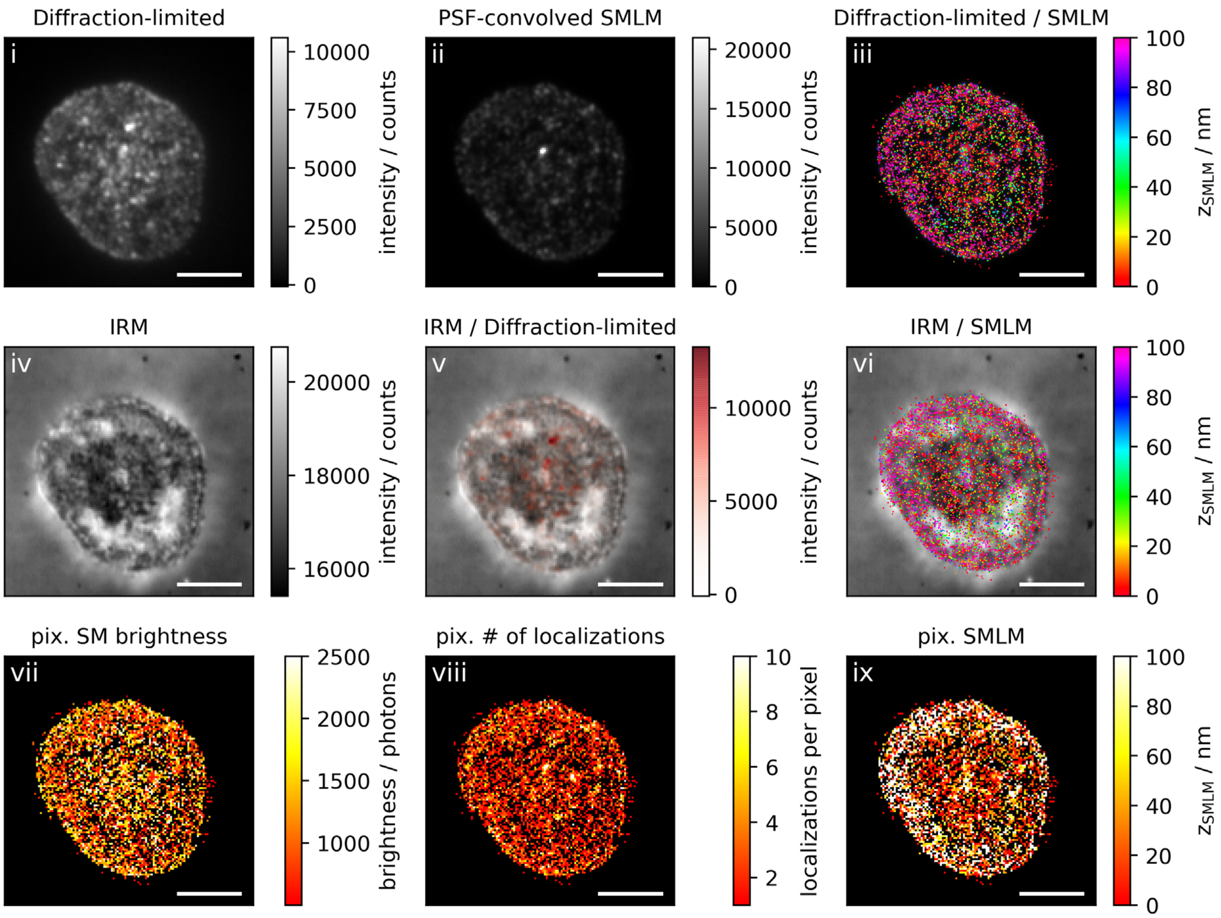
Diffraction-limited
i
10000
7500
5000
2500
0
intensity / counts
PSF-convolved SMLM
ii
20000
15000
10000
5000
0
intensity / counts
Diffraction-limited / SMLM
iii
100
80
60
40
20
0
$z_{SMLM}$ / nm
IRM
iv
20000
18000
16000
intensity / counts
IRM / Diffraction-limited
v
10000
5000
0
intensity / counts
IRM / SMLM
vi
100
80
60
40
20
0
$z_{SMLM}$ / nm
pix. SM brightness
vii
2500
2000
1500
1000
brightness / photons
pix. # of localizations
viii
10
8
6
4
2
localizations per pixel
pix. SMLM
ix
100
80
60
40
20
0
$z_{SMLM}$ / nm

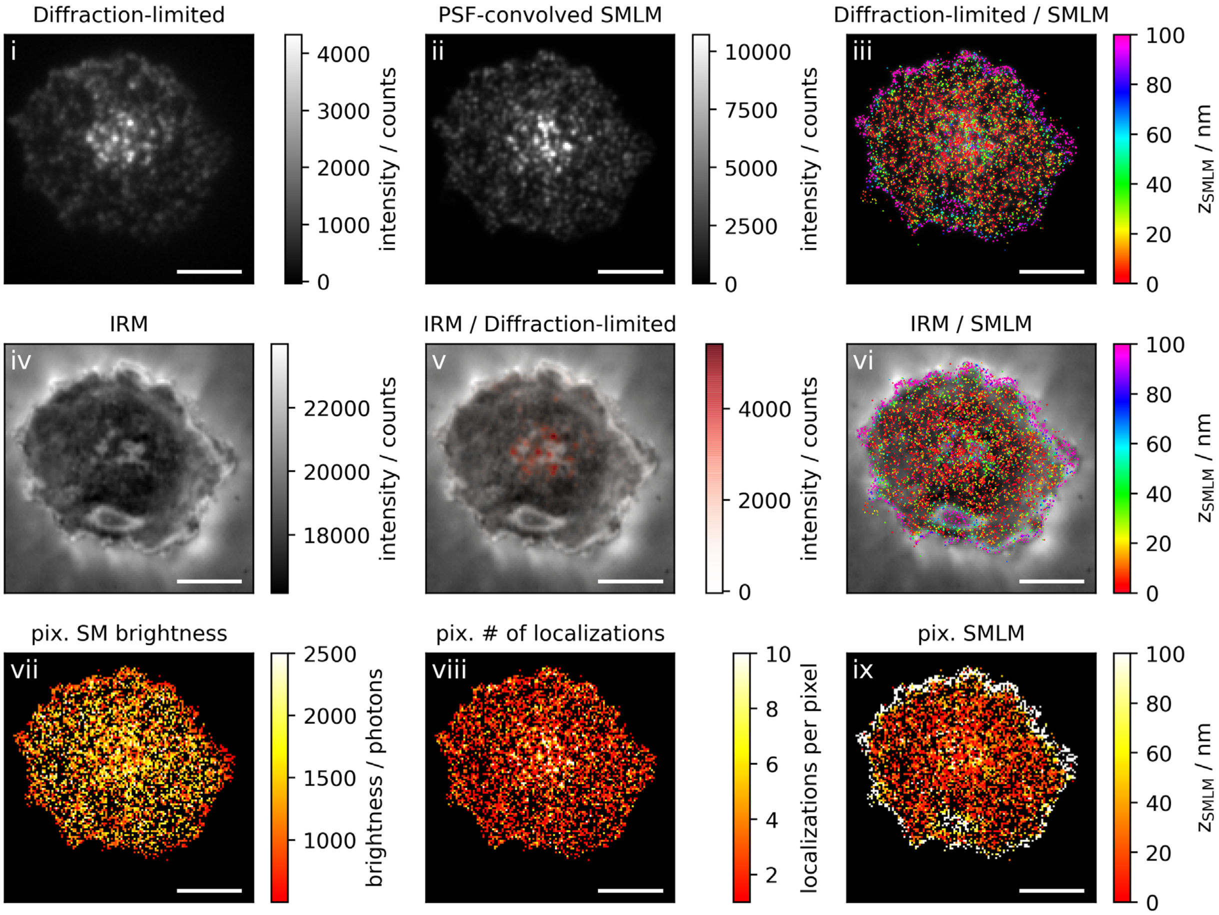

Diffraction-limited
i
intensity / counts
PSF-convolved SMLM
ii
Diffraction-limited / SMLM
iii
$z_{SMLM}$ / nm
IRM
iv
IRM / Diffraction-limited
v
IRM / SMLM
vi
pix. SM brightness
vii
brightness / photons
pix. # of localizations
viii
localizations per pixel
pix. SMLM
ix

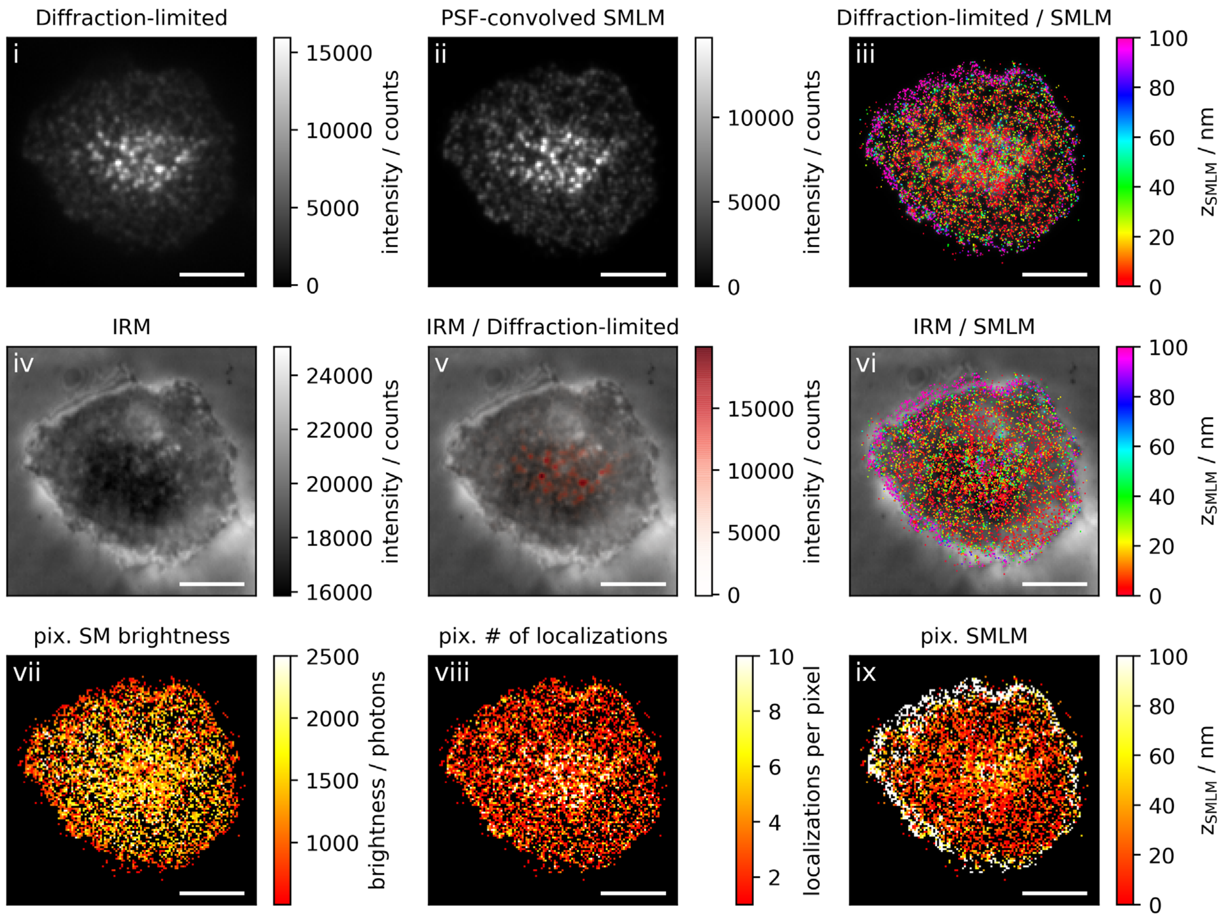

Diffraction-limited
i
15000
10000
5000
0
intensity / counts
PSF-convolved SMLM
ii
10000
5000
0
intensity / counts
Diffraction-limited / SMLM
iii
100
80
60
40
20
0
zSMLM / nm
IRM
iv
24000
22000
20000
18000
16000
intensity / counts
IRM / Diffraction-limited
v
15000
10000
5000
0
intensity / counts
IRM / SMLM
vi
100
80
60
40
20
0
zSMLM / nm
pix. SM brightness
vii
2500
2000
1500
1000
brightness / photons
pix. # of localizations
viii
10
8
6
4
2
localizations per pixel
pix. SMLM
ix
100
80
60
40
20
0
zSMLM / nm

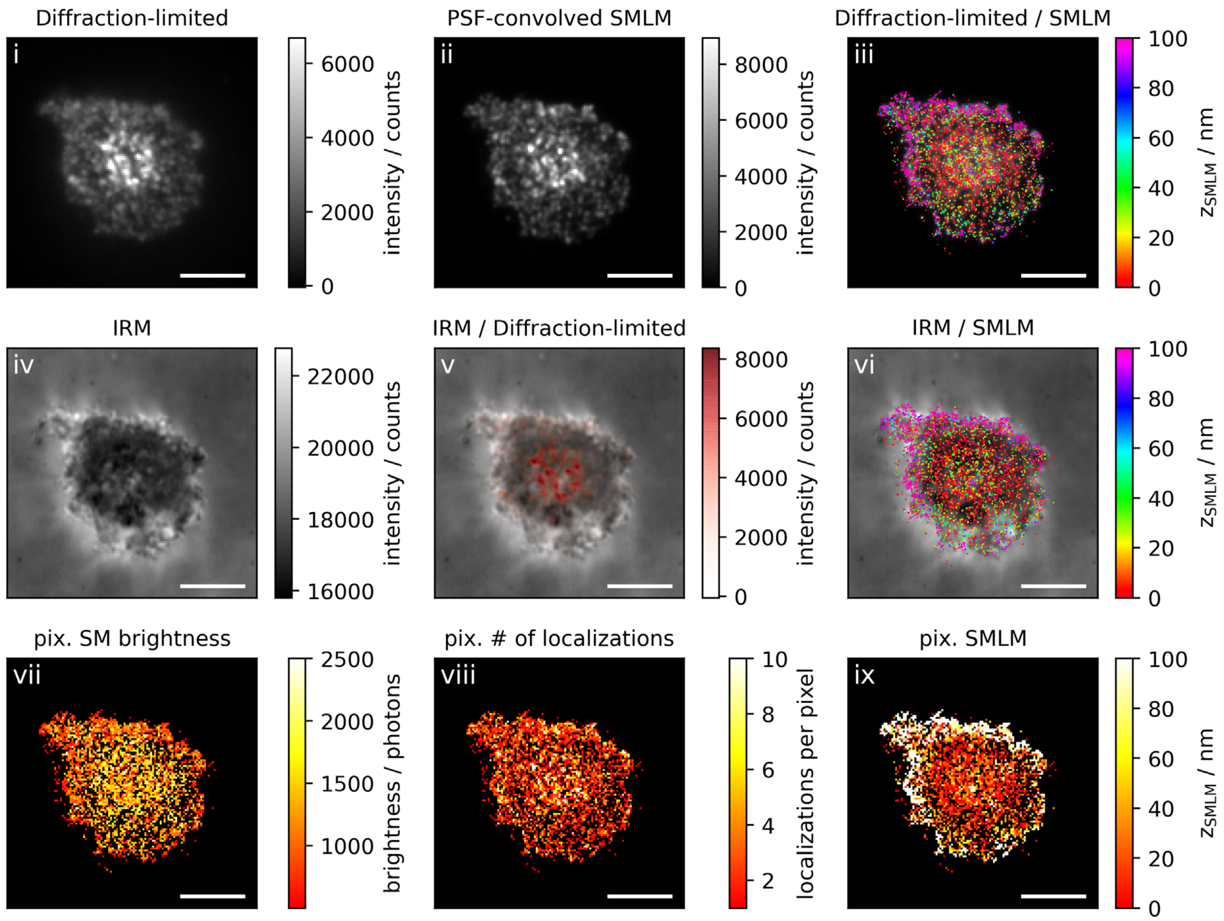

Diffraction-limited
i
6000
4000
2000
0
intensity / counts
PSF-convolved SMLM
ii
8000
6000
4000
2000
0
intensity / counts
Diffraction-limited / SMLM
iii
100
80
60
40
20
0
z_SMLM / nm
IRM
iv
22000
20000
18000
16000
intensity / counts
IRM / Diffraction-limited
v
8000
6000
4000
2000
0
intensity / counts
IRM / SMLM
vi
100
80
60
40
20
0
z_SMLM / nm
pix. SM brightness
vii
2500
2000
1500
1000
brightness / photons
pix. # of localizations
viii
10
8
6
4
2
localizations per pixel
pix. SMLM
ix
100
80
60
40
20
0
z_SMLM / nm

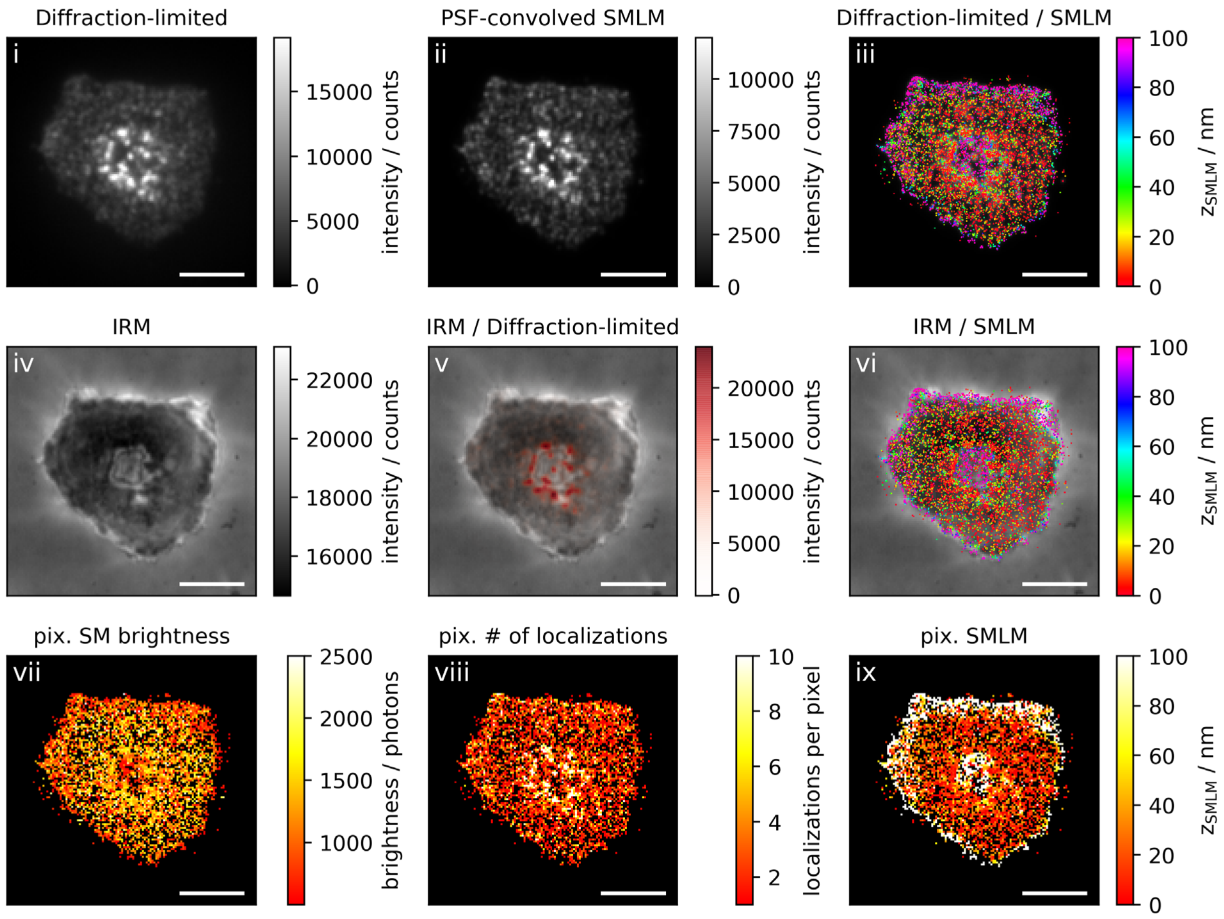
Diffraction-limited
i
15000
10000
5000
0
intensity / counts
PSF-convolved SMLM
ii
10000
7500
5000
2500
0
intensity / counts
Diffraction-limited / SMLM
iii
100
80
60
40
20
0
$z_{SMLM}$ / nm
IRM
iv
22000
20000
18000
16000
intensity / counts
IRM / Diffraction-limited
v
20000
15000
10000
5000
0
intensity / counts
IRM / SMLM
vi
100
80
60
40
20
0
$z_{SMLM}$ / nm
pix. SM brightness
vii
2500
2000
1500
1000
brightness / photons
pix. # of localizations
viii
10
8
6
4
2
localizations per pixel
pix. SMLM
ix
100
80
60
40
20
0
$z_{SMLM}$ / nm

**Figure 2a: Activating conditions, low ICAM-1 density, fixation: 5-10 min post seeding**

**Correlative 3D-SMLM, IRM, and diffraction-limited TIR microscopy of the immunological synapse.** T cells were activated on an SLB functionalized with I-$E^k$/MCC, B7-1 and low density of ICAM-1, and fixed 5-10 minutes post seeding. The T cell was imaged with IRM and fluorescence microscopy: (i) Diffraction-limited TIR image of the T cell. (ii) Reconstruction of the diffraction-limited image by convolving the 3D-SMLM image with the corresponding psf. (iii) Overlay of the diffraction-limited TIR image with the 3D-SMLM image. Color-code indicates distance to the coverslip $z_{SMLM}$. (iv) IRM image. (v) Overlay of the IRM image with the diffraction-limited image. (vi) Overlay of the IRM and the 3D-SMLM image. Bottom row images were generated by calculating the pixel-wise average of the 3D-SMLM images (pixel size of 146nm is consistent with diffraction-limited image) according to pixelated mean single molecule (SM) intensity (vii), pixelated number of localizations (viii) and pixelated mean $z_{SMLM}$ (ix). Scale bars 5 µm.

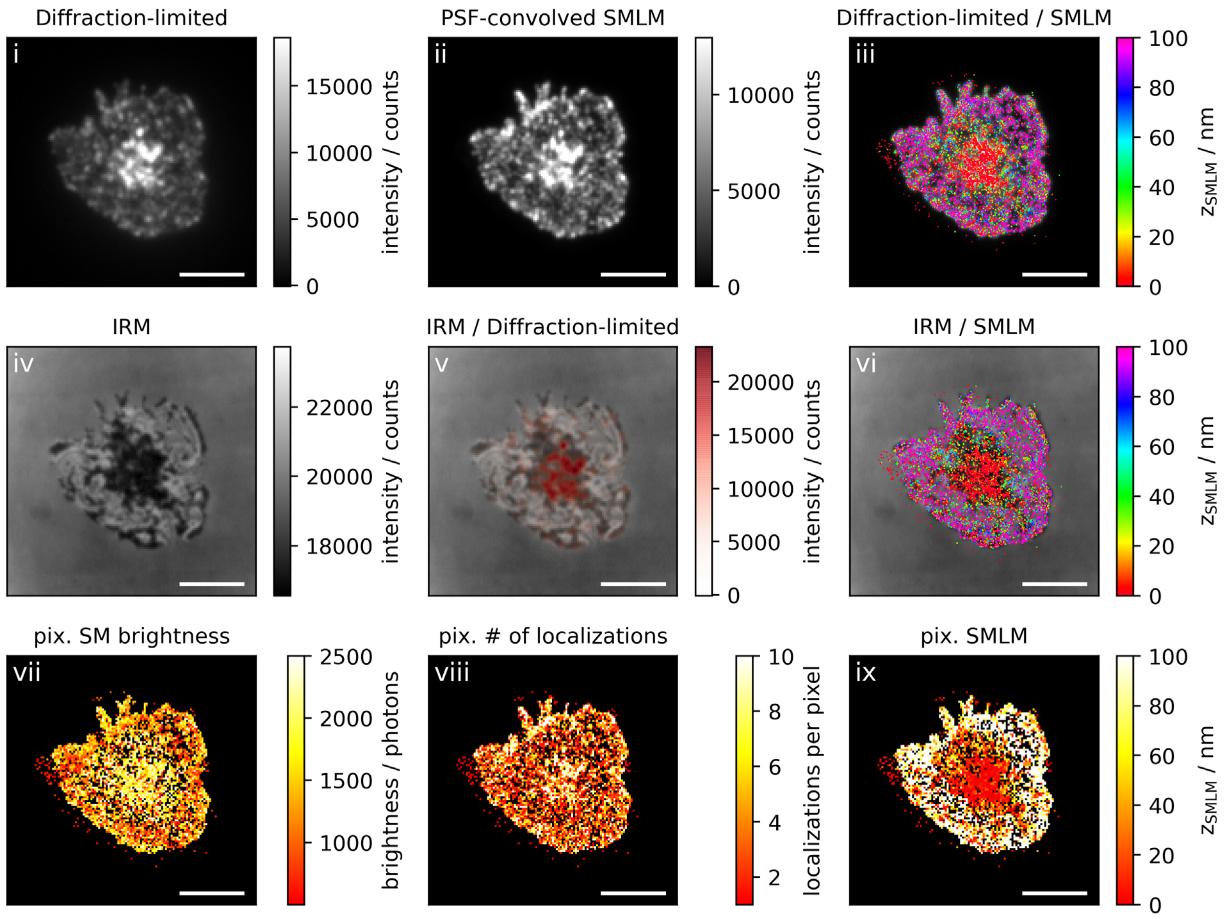
Diffraction-limited
i
15000
10000
5000
0
intensity / counts
PSF-convolved SMLM
ii
10000
5000
0
intensity / counts
Diffraction-limited / SMLM
iii
100
80
60
40
20
0
zSMLM / nm
IRM
iv
22000
20000
18000
intensity / counts
IRM / Diffraction-limited
v
20000
15000
10000
5000
0
intensity / counts
IRM / SMLM
vi
100
80
60
40
20
0
zSMLM / nm
pix. SM brightness
vii
2500
2000
1500
1000
brightness / photons
pix. # of localizations
viii
10
8
6
4
2
localizations per pixel
pix. SMLM
ix
100
80
60
40
20
0
zSMLM / nm

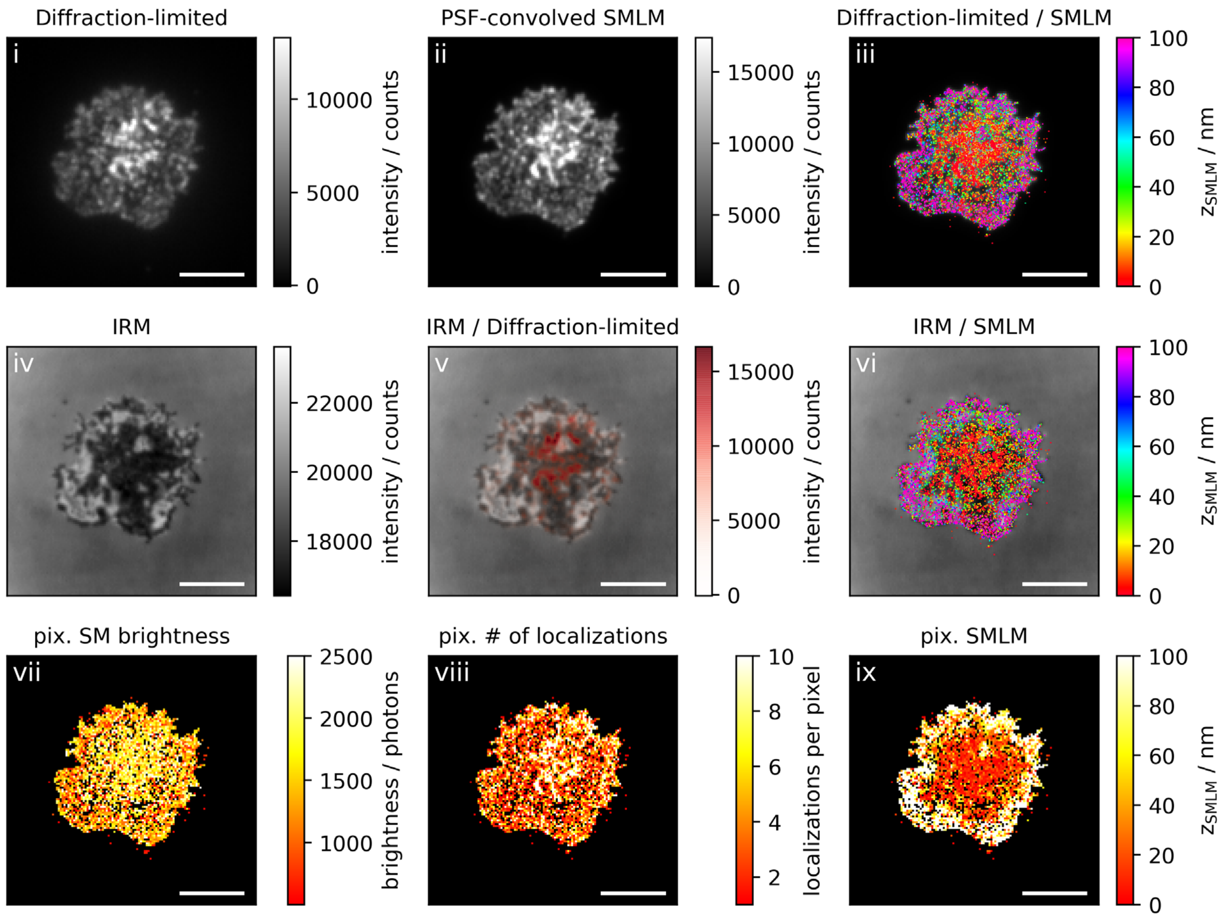
Diffraction-limited
i
0
5000
10000
intensity / counts
PSF-convolved SMLM
ii
0
5000
10000
15000
intensity / counts
Diffraction-limited / SMLM
iii
0
20
40
60
80
100
zSMLM / nm
IRM
iv
18000
20000
22000
intensity / counts
IRM / Diffraction-limited
v
0
5000
10000
15000
intensity / counts
IRM / SMLM
vi
0
20
40
60
80
100
zSMLM / nm
pix. SM brightness
vii
1000
1500
2000
2500
brightness / photons
pix. # of localizations
viii
2
4
6
8
10
localizations per pixel
pix. SMLM
ix
0
20
40
60
80
100
zSMLM / nm

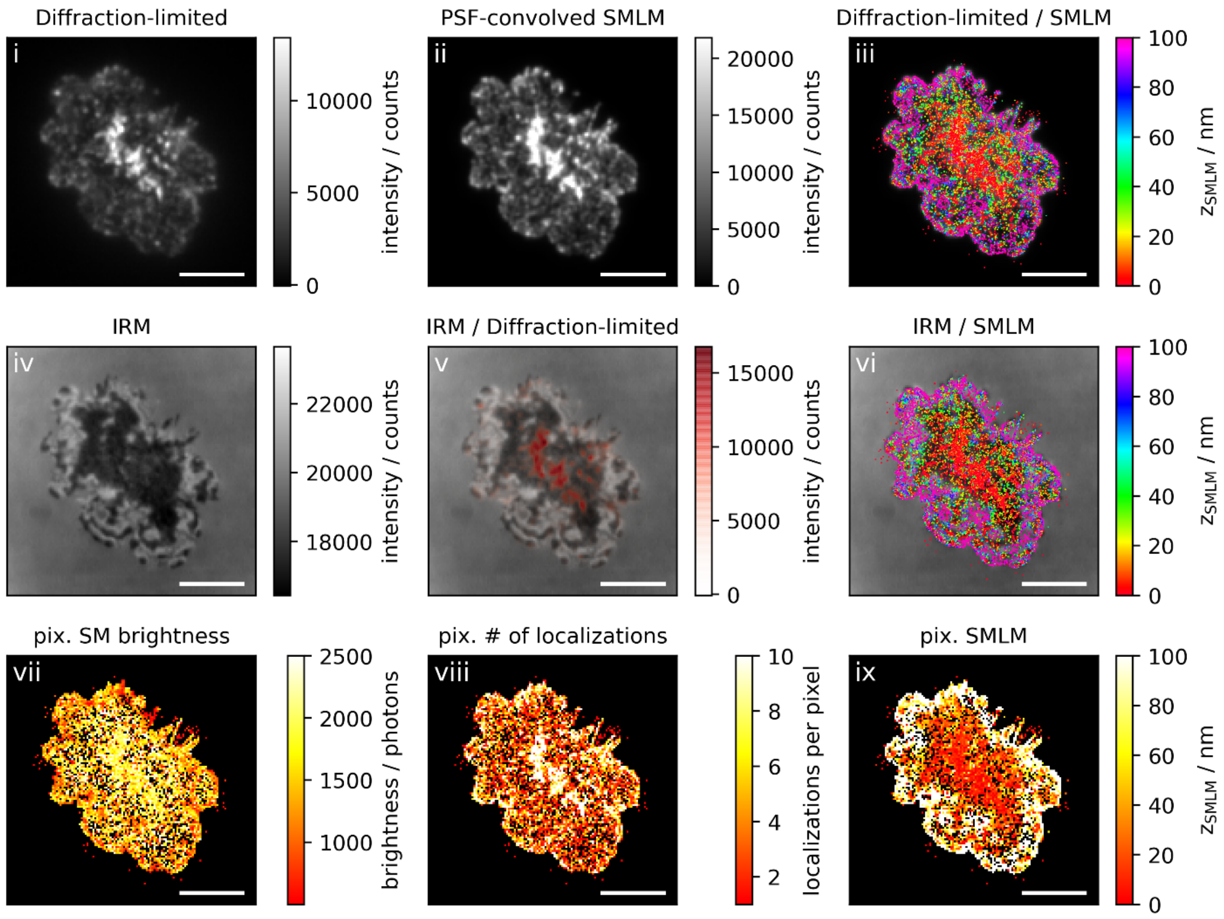

Diffraction-limited
i
intensity / counts
PSF-convolved SMLM
ii
Diffraction-limited / SMLM
iii
$z_{SMLM}$ / nm
IRM
iv
IRM / Diffraction-limited
v
IRM / SMLM
vi
pix. SM brightness
vii
brightness / photons
pix. # of localizations
viii
localizations per pixel
pix. SMLM
ix

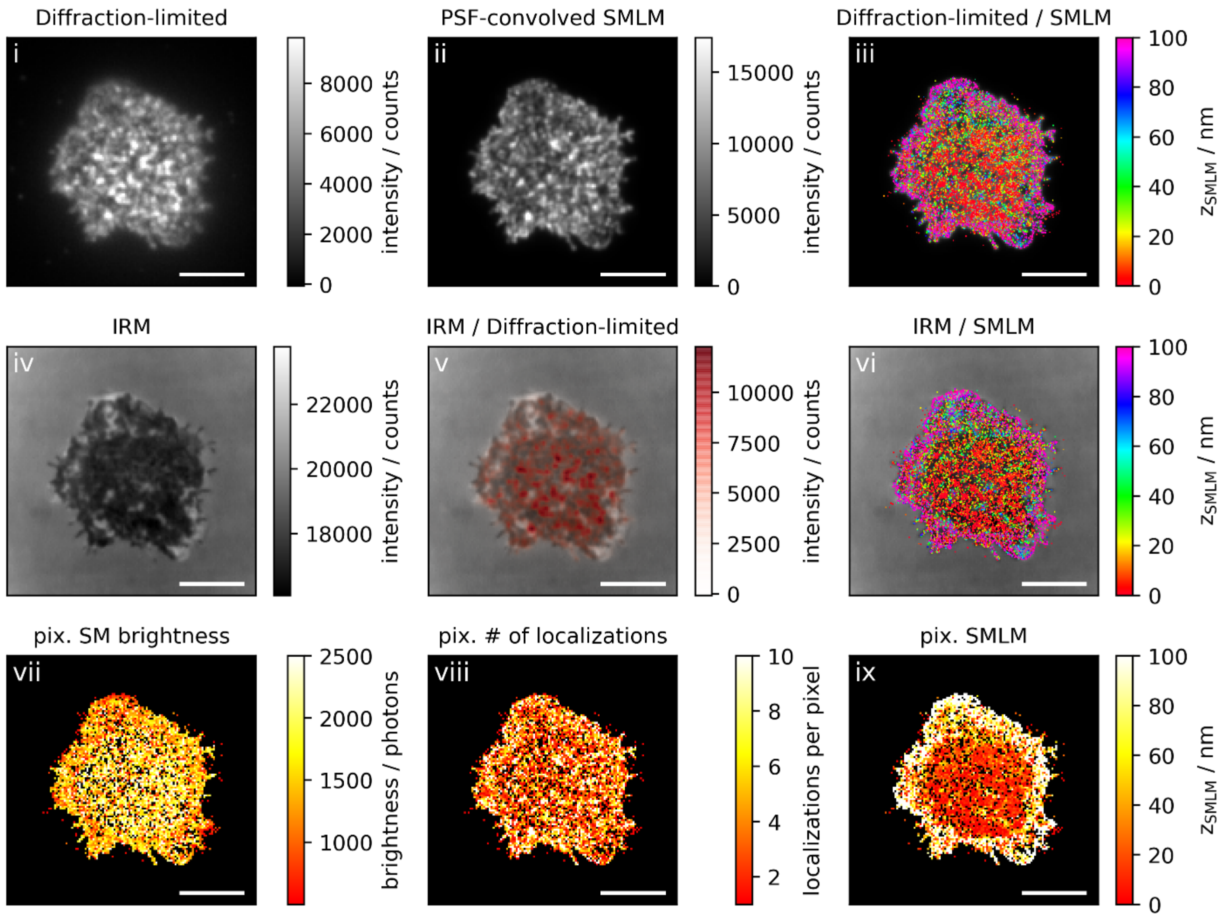
Diffraction-limited
i
intensity / counts
0
2000
4000
6000
8000
PSF-convolved SMLM
ii
intensity / counts
0
5000
10000
15000
Diffraction-limited / SMLM
iii
$z_{SMLM}$ / nm
0
20
40
60
80
100
IRM
iv
intensity / counts
18000
20000
22000
IRM / Diffraction-limited
v
intensity / counts
0
2500
5000
7500
10000
IRM / SMLM
vi
$z_{SMLM}$ / nm
0
20
40
60
80
100
pix. SM brightness
vii
brightness / photons
1000
1500
2000
2500
pix. # of localizations
viii
localizations per pixel
2
4
6
8
10
pix. SMLM
ix
$z_{SMLM}$ / nm
0
20
40
60
80
100

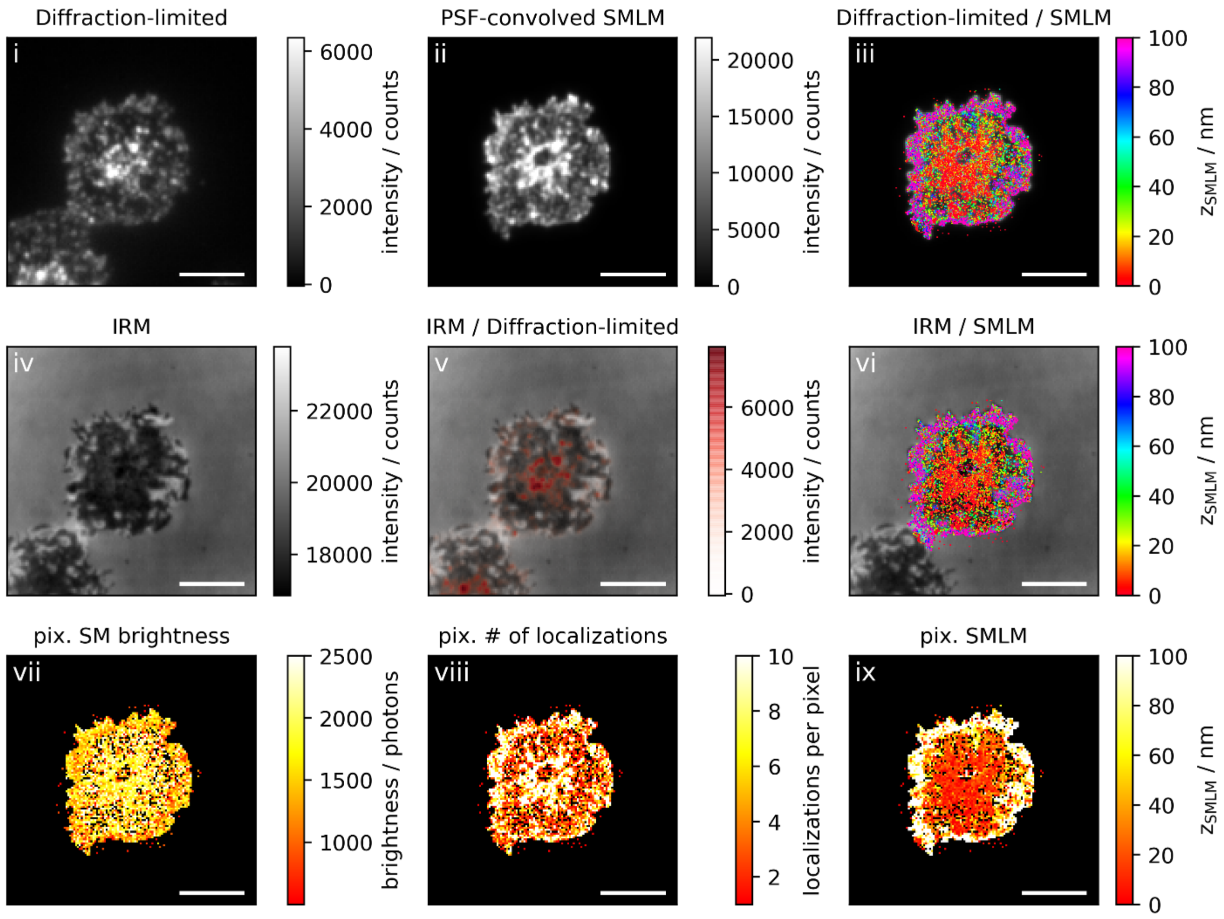
Diffraction-limited
i
intensity / counts
PSF-convolved SMLM
ii
intensity / counts
Diffraction-limited / SMLM
iii
$z_{SMLM}$ / nm
IRM
iv
intensity / counts
IRM / Diffraction-limited
v
intensity / counts
IRM / SMLM
vi
$z_{SMLM}$ / nm
pix. SM brightness
vii
brightness / photons
pix. # of localizations
viii
localizations per pixel
pix. SMLM
ix
$z_{SMLM}$ / nm

**Figure 2b: Activating conditions, low ICAM-1 density, fixation: 10 min post seeding**

**Correlative 3D-SMLM, IRM, and diffraction-limited TIR microscopy of the immunological synapse.** T cells were activated on an SLB functionalized with I-$E^k$/MCC, B7-1 and low density of ICAM-1, and fixed 10 minutes post seeding. The T cell was imaged with IRM and fluorescence microscopy: (i) Diffraction-limited TIR image of the T cell. (ii) Reconstruction of the diffraction-limited image by convolving the 3D-SMLM image with the corresponding psf. (iii) Overlay of the diffraction-limited TIR image with the 3D-SMLM image. Color-code indicates distance to the coverslip $z_{SMLM}$. (iv) IRM image. (v) Overlay of the IRM image with the diffraction-limited image. (vi) Overlay of the IRM and the 3D-SMLM image. Bottom row images were generated by calculating the pixel-wise average of the 3D-SMLM images (pixel size of 146nm is consistent with diffraction-limited image) according to pixelated mean single molecule (SM) intensity (vii), pixelated number of localizations (viii) and pixelated mean $z_{SMLM}$ (ix). Scale bars 5 µm.

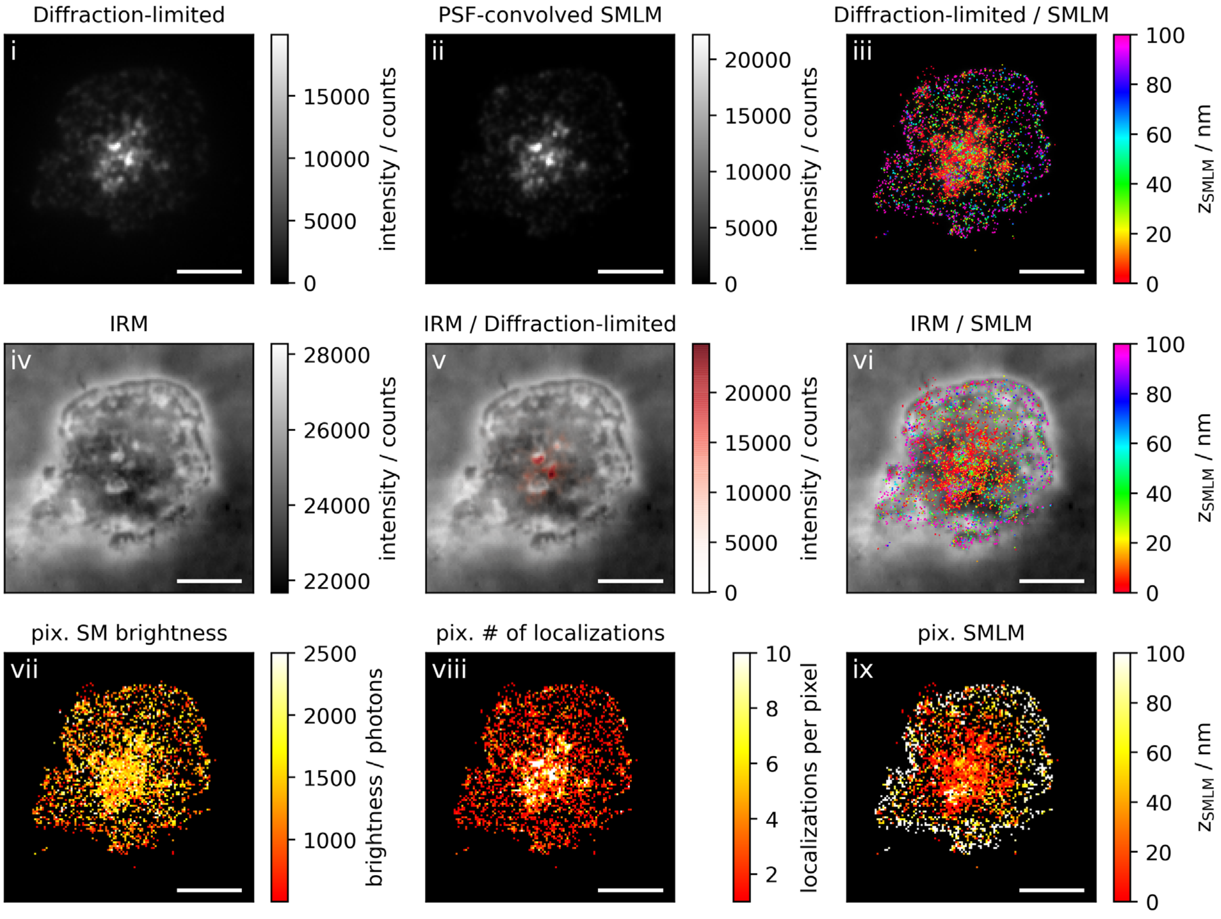
Diffraction-limited
i
intensity / counts
PSF-convolved SMLM
ii
Diffraction-limited / SMLM
iii
$z_{SMLM}$ / nm
IRM
iv
IRM / Diffraction-limited
v
IRM / SMLM
vi
pix. SM brightness
vii
brightness / photons
pix. # of localizations
viii
localizations per pixel
pix. SMLM
ix

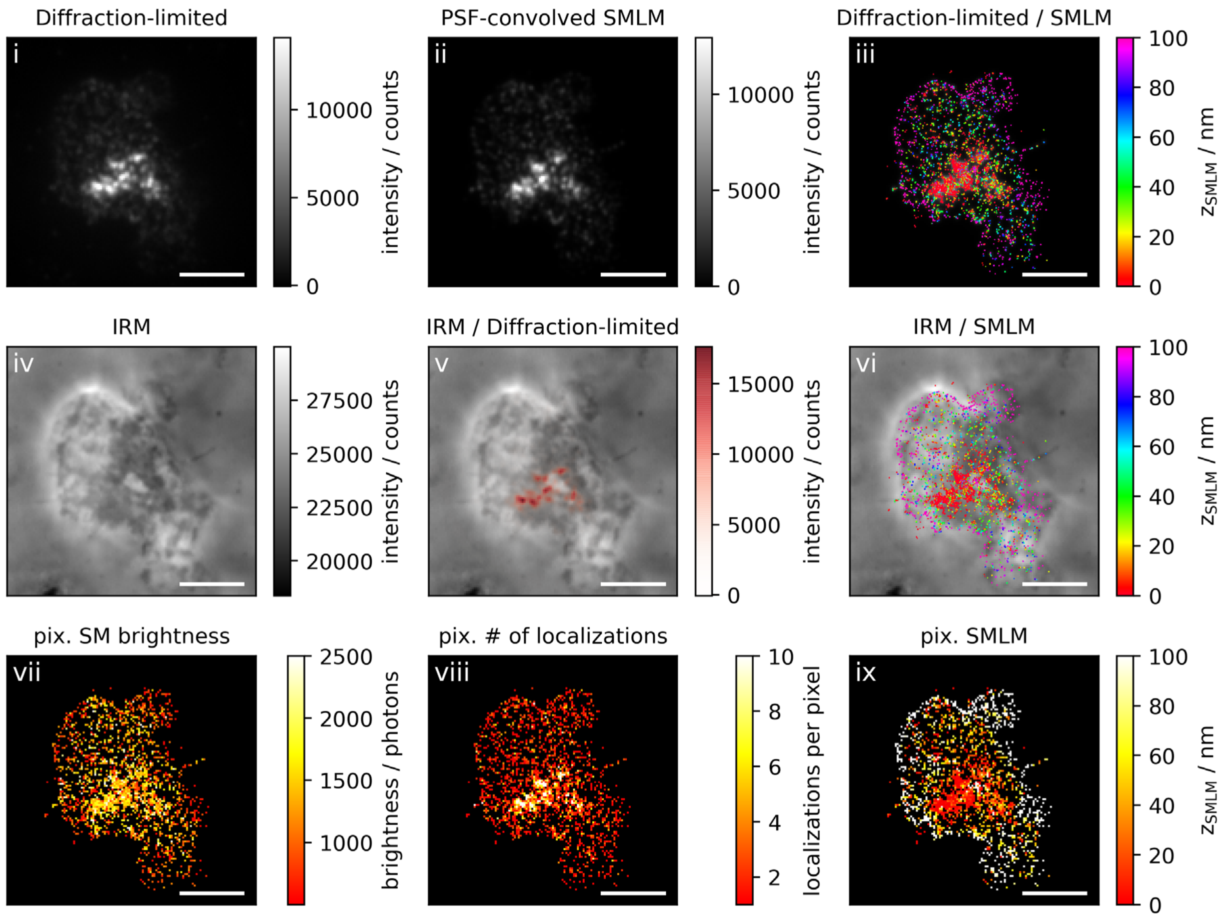

Diffraction-limited
i
intensity / counts
0
5000
10000
PSF-convolved SMLM
ii
intensity / counts
0
5000
10000
Diffraction-limited / SMLM
iii
$z_{SMLM}$ / nm
0
20
40
60
80
100
IRM
iv
intensity / counts
20000
22500
25000
27500
IRM / Diffraction-limited
v
intensity / counts
0
5000
10000
15000
IRM / SMLM
vi
$z_{SMLM}$ / nm
0
20
40
60
80
100
pix. SM brightness
vii
brightness / photons
1000
1500
2000
2500
pix. # of localizations
viii
localizations per pixel
2
4
6
8
10
pix. SMLM
ix
$z_{SMLM}$ / nm
0
20
40
60
80
100

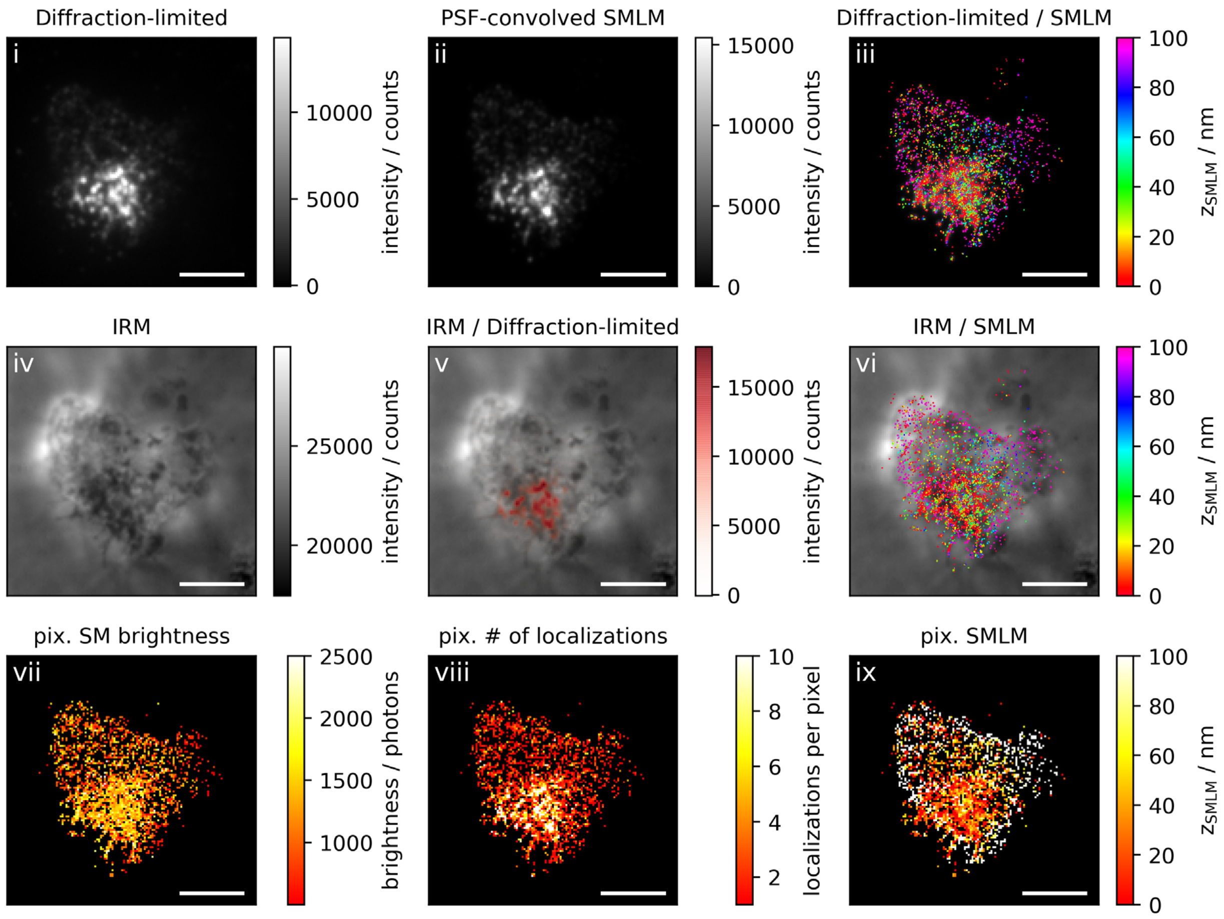

Diffraction-limited
i
10000
5000
0
intensity / counts
PSF-convolved SMLM
ii
15000
10000
5000
0
intensity / counts
Diffraction-limited / SMLM
iii
100
80
60
40
20
0
$z_{SMLM}$ / nm
IRM
iv
25000
20000
intensity / counts
IRM / Diffraction-limited
v
15000
10000
5000
0
intensity / counts
IRM / SMLM
vi
100
80
60
40
20
0
$z_{SMLM}$ / nm
pix. SM brightness
vii
2500
2000
1500
1000
brightness / photons
pix. # of localizations
viii
10
8
6
4
2
localizations per pixel
pix. SMLM
ix
100
80
60
40
20
0
$z_{SMLM}$ / nm

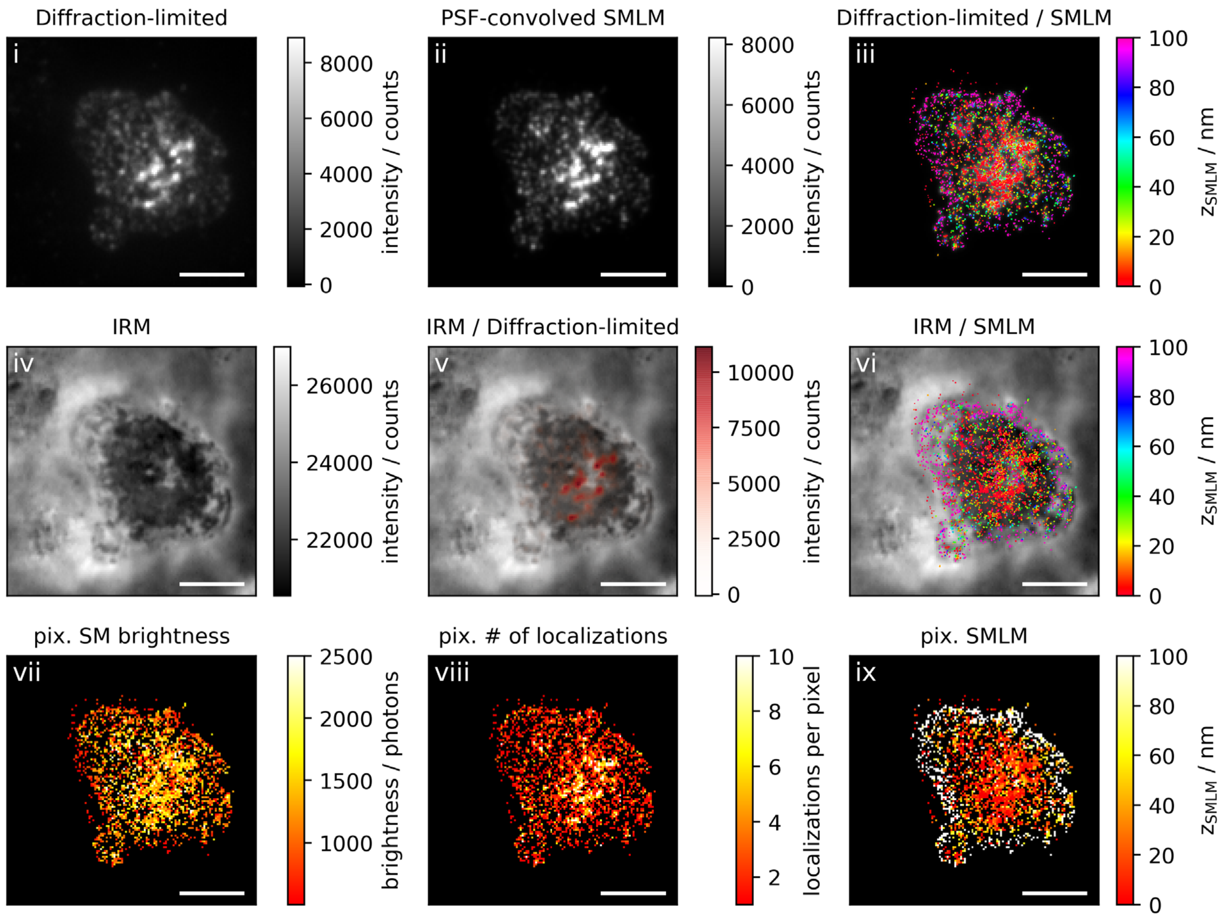

Diffraction-limited
i
intensity / counts
PSF-convolved SMLM
ii
intensity / counts
Diffraction-limited / SMLM
iii
$z_{SMLM}$ / nm
IRM
iv
intensity / counts
IRM / Diffraction-limited
v
intensity / counts
IRM / SMLM
vi
$z_{SMLM}$ / nm
pix. SM brightness
vii
brightness / photons
pix. # of localizations
viii
localizations per pixel
pix. SMLM
ix
$z_{SMLM}$ / nm

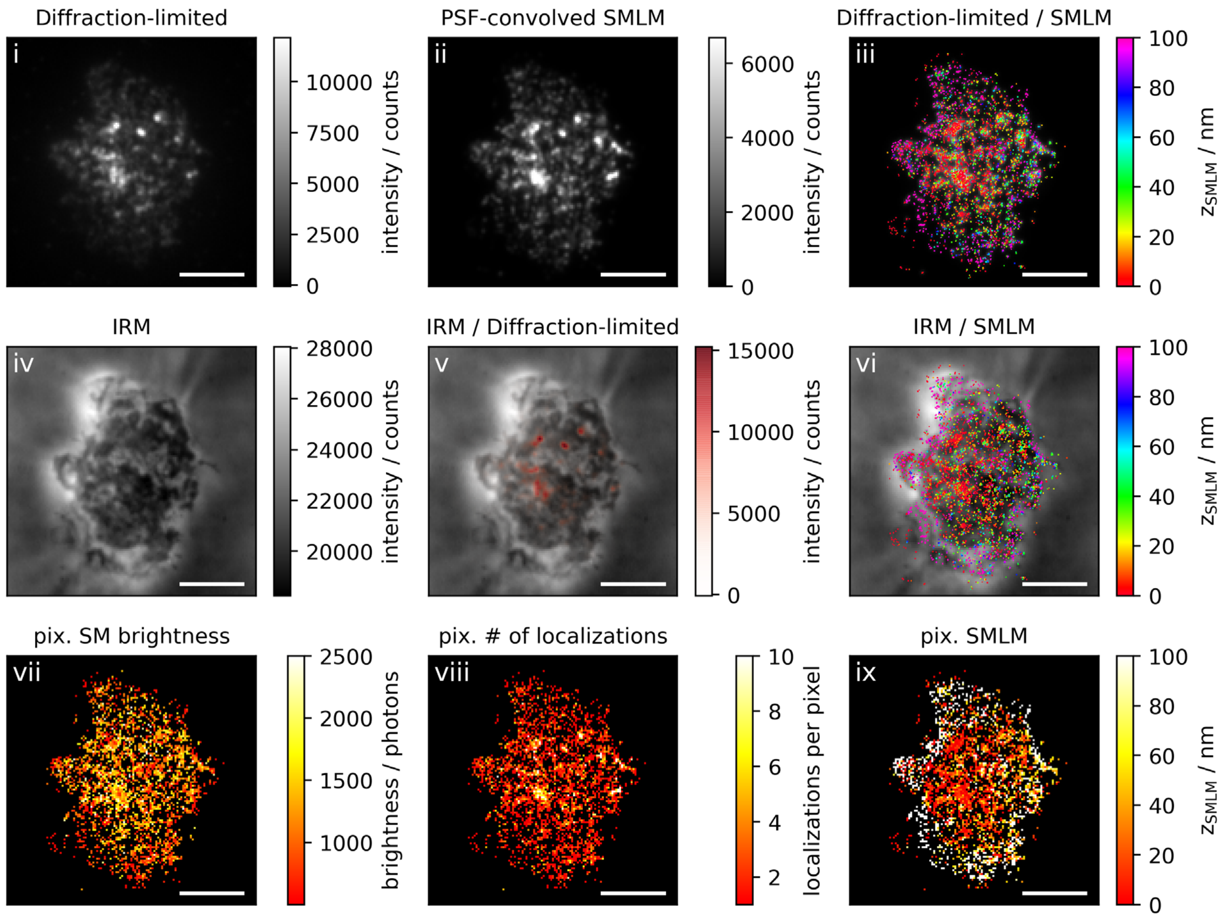
Diffraction-limited
i
intensity / counts
0
2500
5000
7500
10000
PSF-convolved SMLM
ii
intensity / counts
0
2000
4000
6000
Diffraction-limited / SMLM
iii
$z_{SMLM}$ / nm
0
20
40
60
80
100
IRM
iv
intensity / counts
20000
22000
24000
26000
28000
IRM / Diffraction-limited
v
intensity / counts
0
5000
10000
15000
IRM / SMLM
vi
$z_{SMLM}$ / nm
0
20
40
60
80
100
pix. SM brightness
vii
brightness / photons
1000
1500
2000
2500
pix. # of localizations
viii
localizations per pixel
2
4
6
8
10
pix. SMLM
ix
$z_{SMLM}$ / nm
0
20
40
60
80
100

**Figure 2c: Activating conditions, low ICAM-1 density, fixation: 10-15 min post seeding**

**Correlative 3D-SMLM, IRM, and diffraction-limited TIR microscopy of the immunological synapse.** T cells were activated on an SLB functionalized with I-$E^k$/MCC, B7-1 and low density of ICAM-1, and fixed 10-15 minutes post seeding. The T cell was imaged with IRM and fluorescence microscopy: (i) Diffraction-limited TIR image of the T cell. (ii) Reconstruction of the diffraction-limited image by convolving the 3D-SMLM image with the corresponding psf. (iii) Overlay of the diffraction-limited TIR image with the 3D-SMLM image. Color-code indicates distance to the coverslip $z_{SMLM}$. (iv) IRM image. (v) Overlay of the IRM image with the diffraction-limited image. (vi) Overlay of the IRM and the 3D-SMLM image. Bottom row images were generated by calculating the pixel-wise average of the 3D-SMLM images (pixel size of 146nm is consistent with diffraction-limited image) according to pixelated mean single molecule (SM) intensity (vii), pixelated number of localizations (viii) and pixelated mean $z_{SMLM}$ (ix). Scale bars 5 µm.

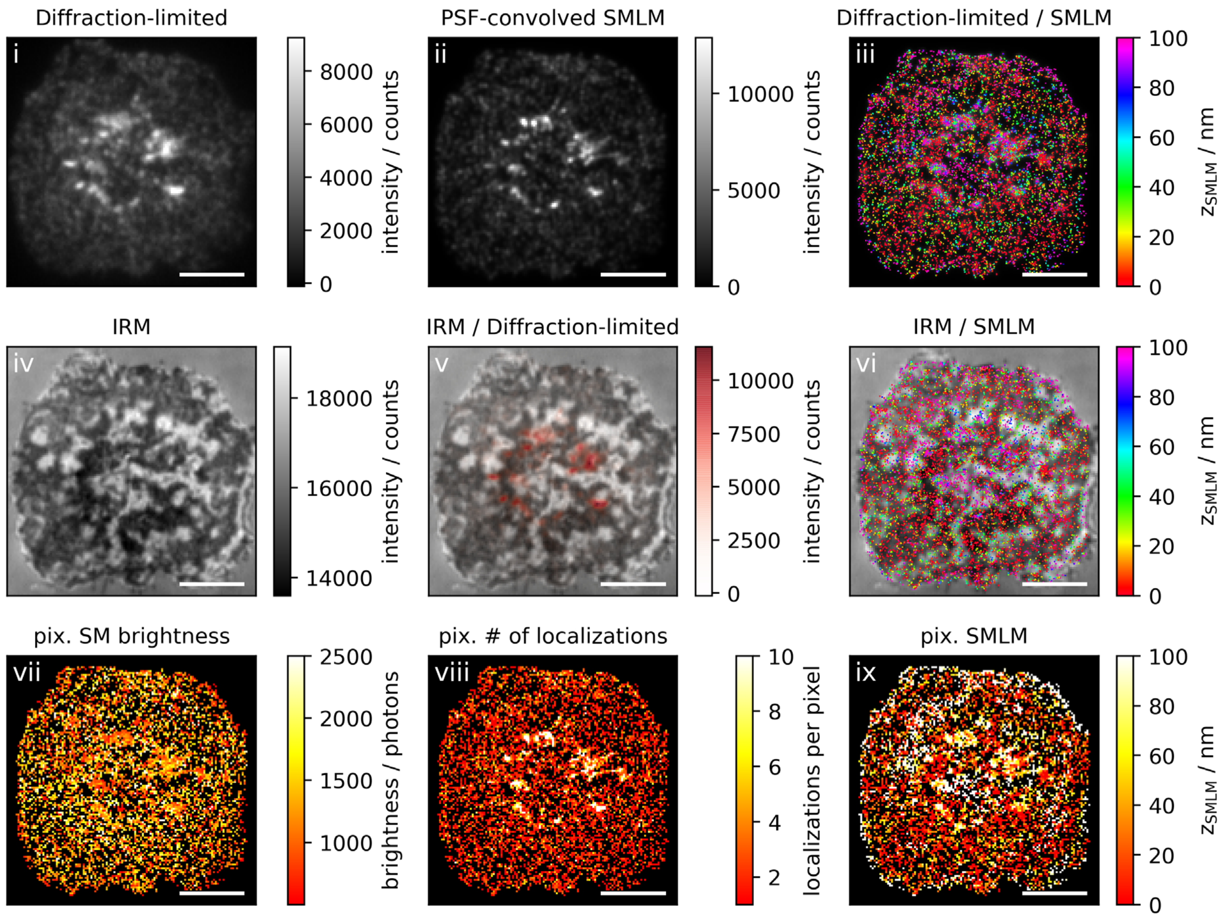
Diffraction-limited
i
8000
6000
4000
2000
0
intensity / counts
PSF-convolved SMLM
ii
10000
5000
0
intensity / counts
Diffraction-limited / SMLM
iii
100
80
60
40
20
0
$z_{SMLM}$ / nm
IRM
iv
18000
16000
14000
intensity / counts
IRM / Diffraction-limited
v
10000
7500
5000
2500
0
intensity / counts
IRM / SMLM
vi
100
80
60
40
20
0
$z_{SMLM}$ / nm
pix. SM brightness
vii
2500
2000
1500
1000
brightness / photons
pix. # of localizations
viii
10
8
6
4
2
localizations per pixel
pix. SMLM
ix
100
80
60
40
20
0
$z_{SMLM}$ / nm

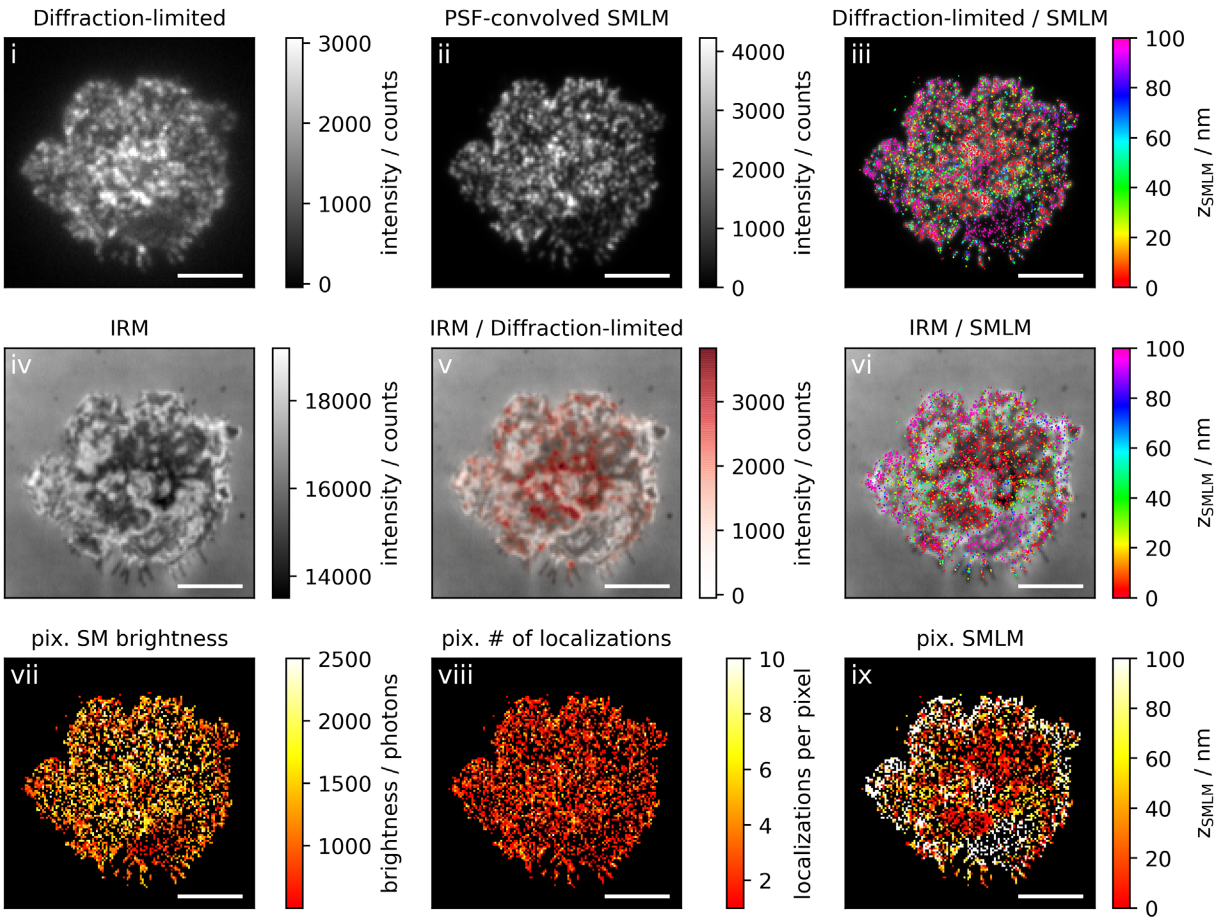
Diffraction-limited
i
3000
2000
1000
0
intensity / counts
PSF-convolved SMLM
ii
4000
3000
2000
1000
0
intensity / counts
Diffraction-limited / SMLM
iii
100
80
60
40
20
0
z_SMLM / nm
IRM
iv
18000
16000
14000
intensity / counts
IRM / Diffraction-limited
v
3000
2000
1000
0
intensity / counts
IRM / SMLM
vi
100
80
60
40
20
0
z_SMLM / nm
pix. SM brightness
vii
2500
2000
1500
1000
brightness / photons
pix. # of localizations
viii
10
8
6
4
2
localizations per pixel
pix. SMLM
ix
100
80
60
40
20
0
z_SMLM / nm

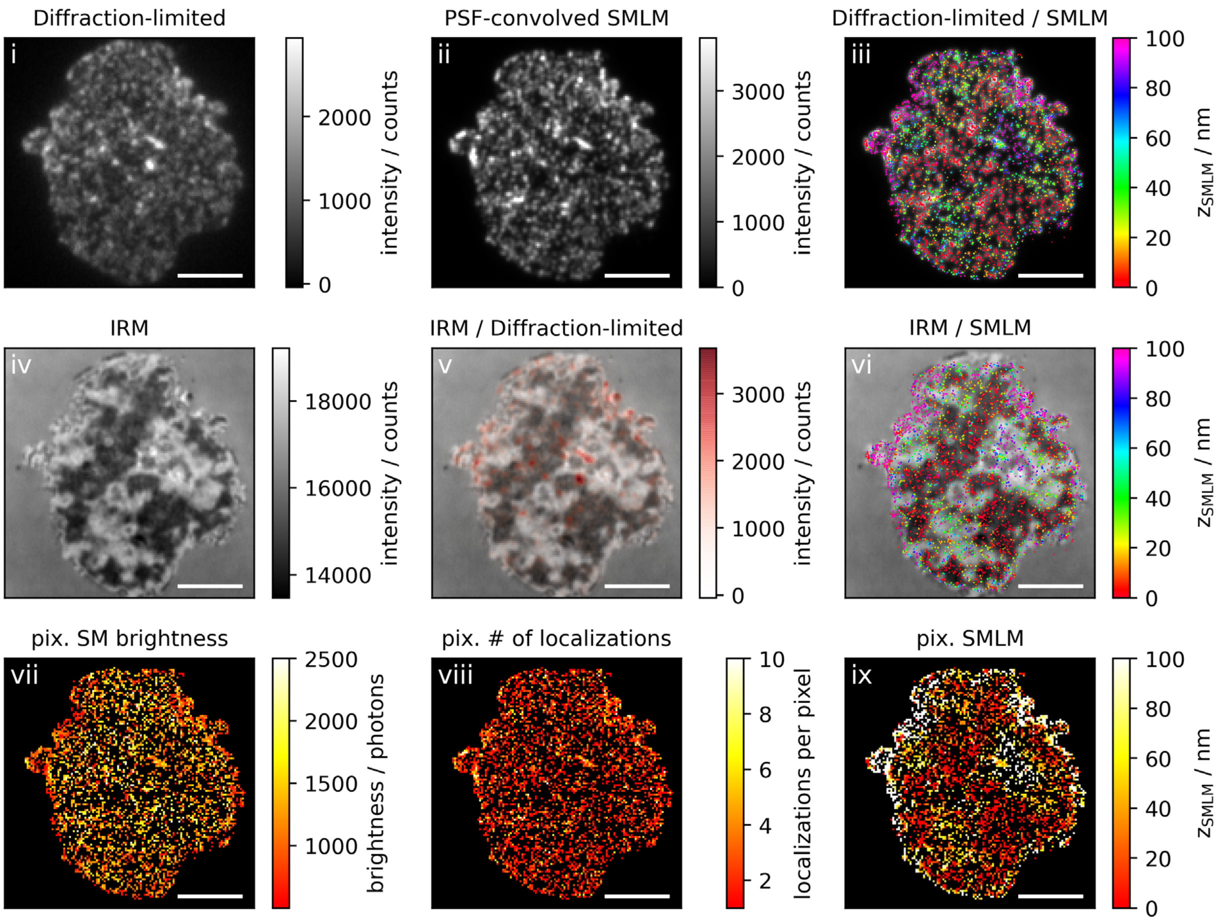
Diffraction-limited
i
intensity / counts
PSF-convolved SMLM
ii
Diffraction-limited / SMLM
iii
$z_{SMLM}$ / nm
IRM
iv
IRM / Diffraction-limited
v
IRM / SMLM
vi
pix. SM brightness
vii
brightness / photons
pix. # of localizations
viii
localizations per pixel
pix. SMLM
ix

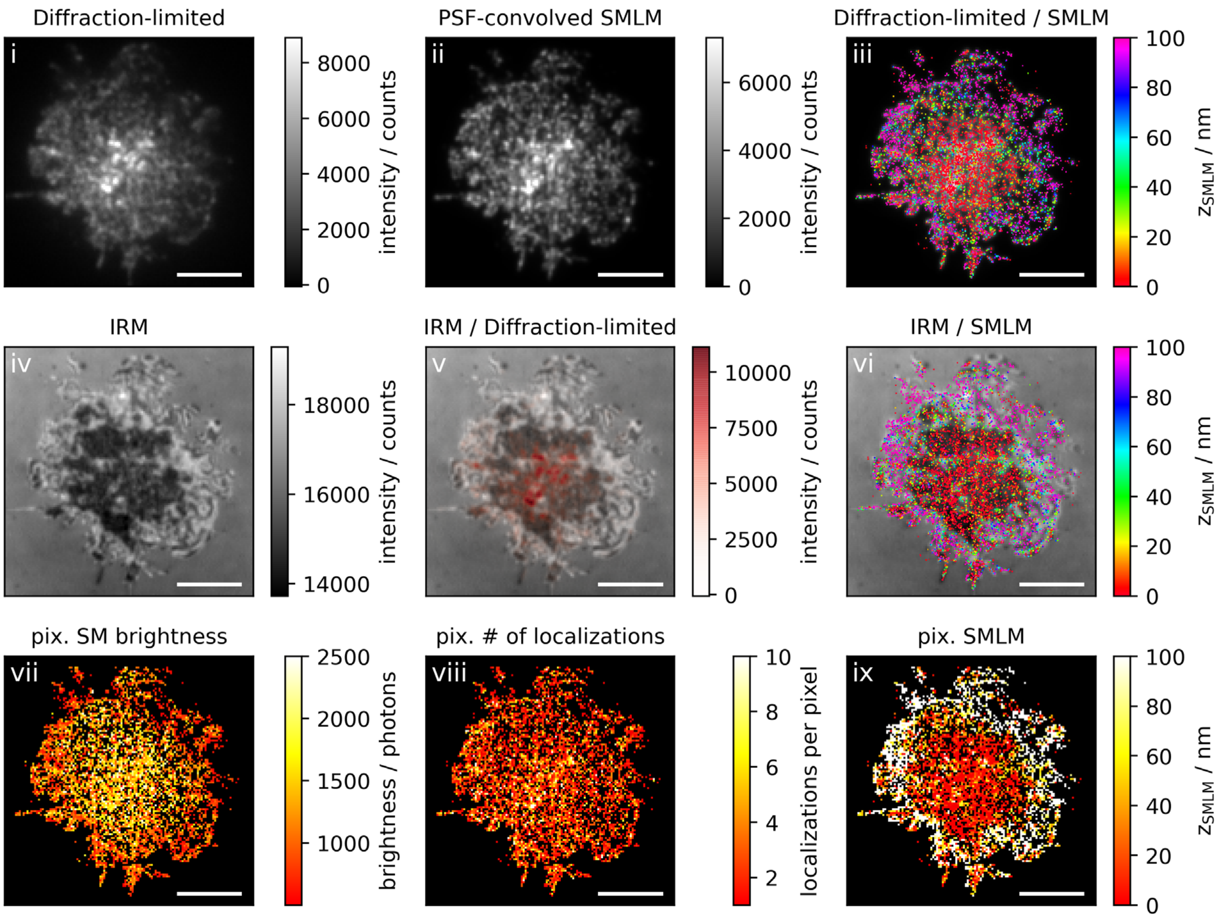
Diffraction-limited
i
intensity / counts
PSF-convolved SMLM
ii
intensity / counts
Diffraction-limited / SMLM
iii
$z_{SMLM}$ / nm
IRM
iv
intensity / counts
IRM / Diffraction-limited
v
intensity / counts
IRM / SMLM
vi
$z_{SMLM}$ / nm
pix. SM brightness
vii
brightness / photons
pix. # of localizations
viii
localizations per pixel
pix. SMLM
ix
$z_{SMLM}$ / nm

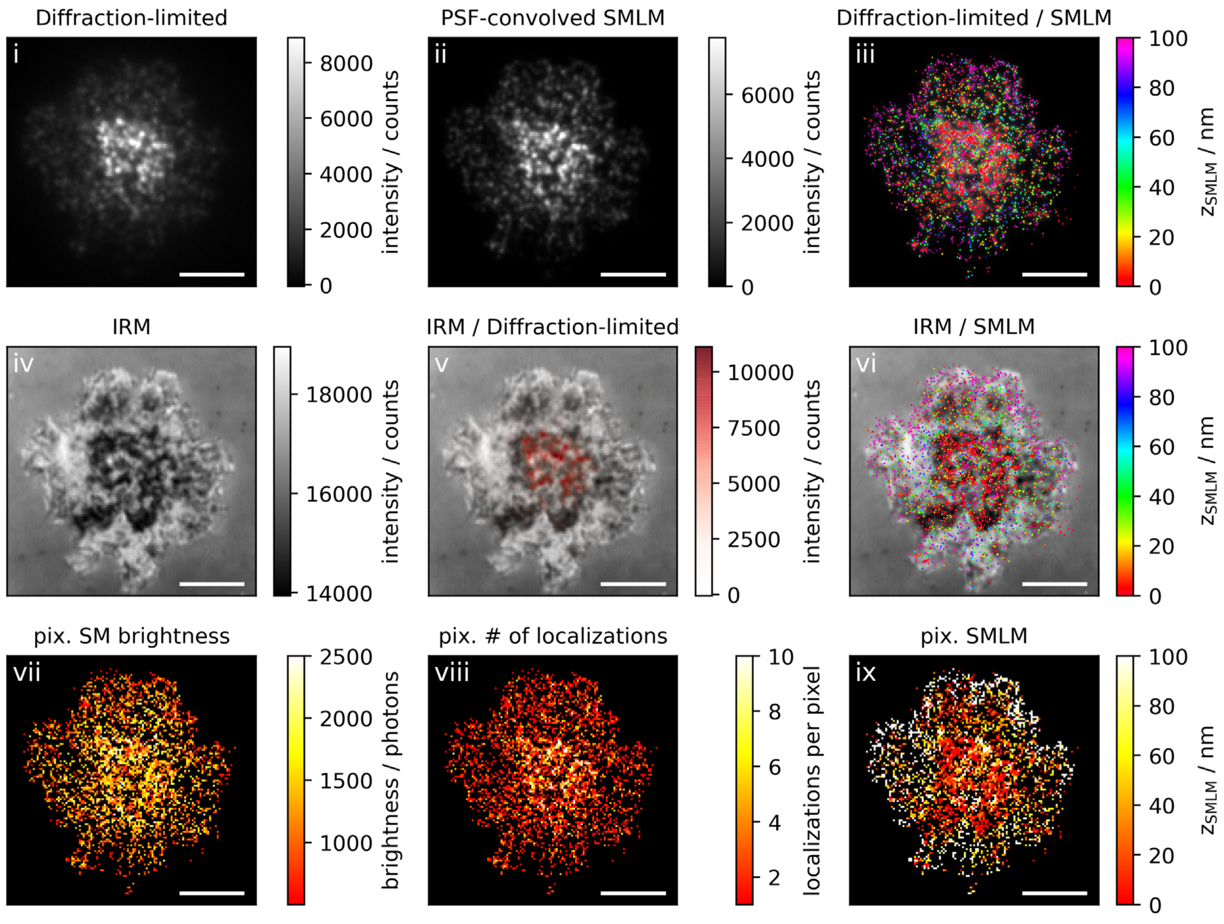
Diffraction-limited
i
8000
6000
4000
2000
0
intensity / counts
PSF-convolved SMLM
ii
6000
4000
2000
0
intensity / counts
Diffraction-limited / SMLM
iii
100
80
60
40
20
0
zSMLM / nm
IRM
iv
18000
16000
14000
intensity / counts
IRM / Diffraction-limited
v
10000
7500
5000
2500
0
intensity / counts
IRM / SMLM
vi
100
80
60
40
20
0
zSMLM / nm
pix. SM brightness
vii
2500
2000
1500
1000
brightness / photons
pix. # of localizations
viii
10
8
6
4
2
localizations per pixel
pix. SMLM
ix
100
80
60
40
20
0
zSMLM / nm

**Figure 3a: Non-activating conditions, high ICAM-1 density, fixation: 5-10 min post seeding**

**Correlative 3D-SMLM, IRM, and diffraction-limited TIR microscopy of the immunological synapse.** T cells were seeded on an SLB functionalized with high density of ICAM-1, and fixed 5-10 minutes post seeding. The T cell was imaged with IRM and fluorescence microscopy: (i) Diffraction-limited TIR image of the T cell. (ii) Reconstruction of the diffraction-limited image by convolving the 3D-SMLM image with the corresponding psf. (iii) Overlay of the diffraction-limited TIR image with the 3D-SMLM image. Color-code indicates distance to the coverslip $z_{SMLM}$. (iv) IRM image. (v) Overlay of the IRM image with the diffraction-limited image. (vi) Overlay of the IRM and the 3D-SMLM image. Bottom row images were generated by calculating the pixel-wise average of the 3D-SMLM images (pixel size of 146nm is consistent with diffraction-limited image) according to pixelated mean single molecule (SM) intensity (vii), pixelated number of localizations (viii) and pixelated mean $z_{SMLM}$ (ix). Scale bars 5 µm.

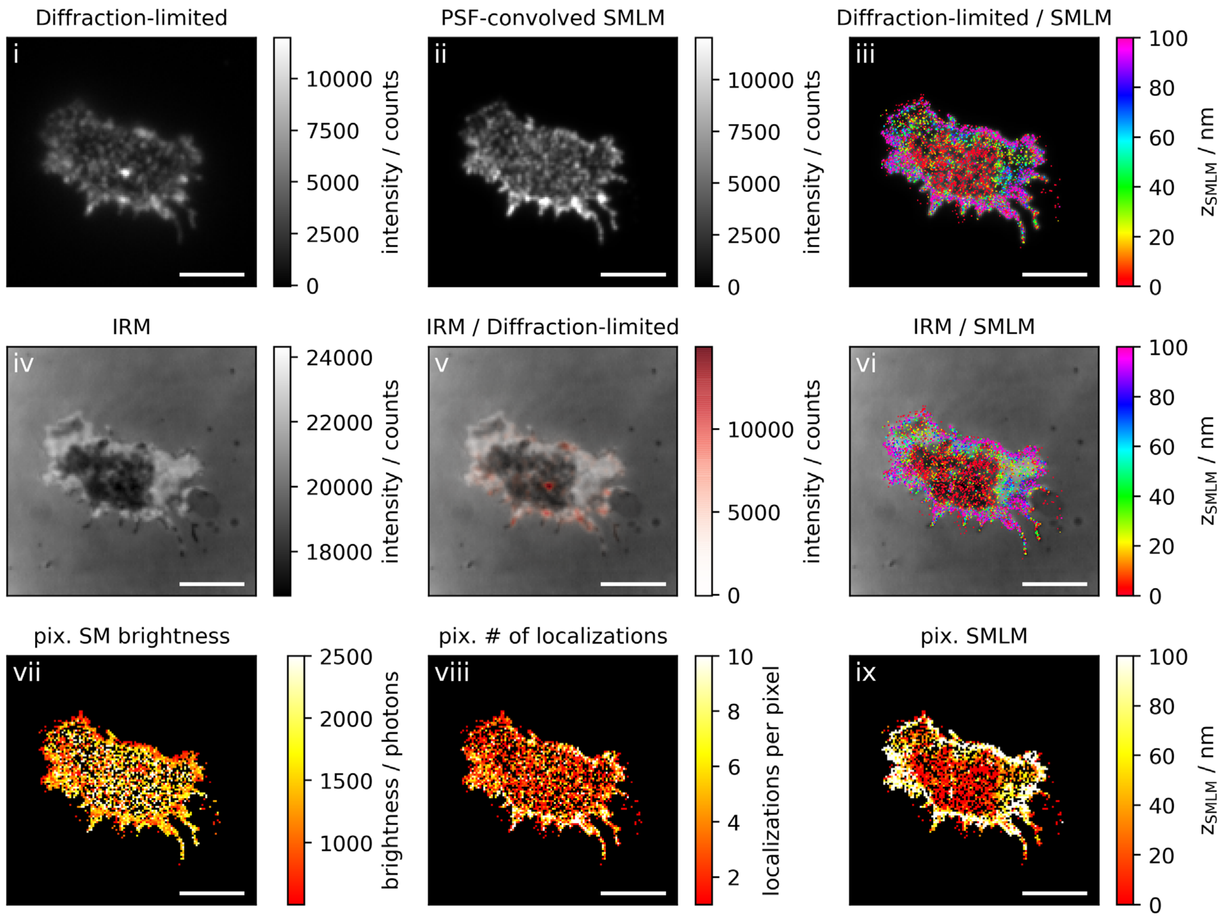
Diffraction-limited
i
intensity / counts
PSF-convolved SMLM
ii
Diffraction-limited / SMLM
iii
$z_{SMLM}$ / nm
IRM
iv
IRM / Diffraction-limited
v
IRM / SMLM
vi
pix. SM brightness
vii
brightness / photons
pix. # of localizations
viii
localizations per pixel
pix. SMLM
ix

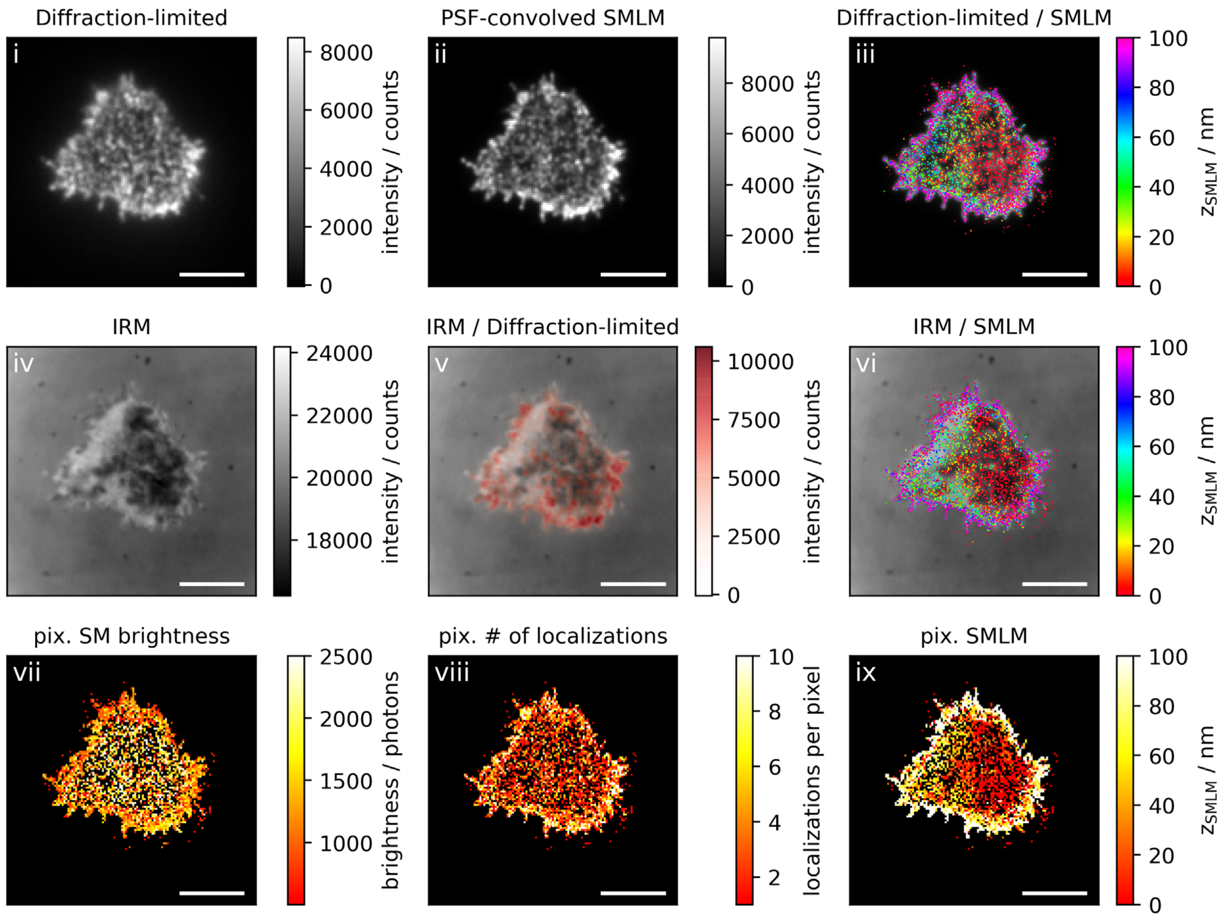
Diffraction-limited
i
8000
6000
4000
2000
0
intensity / counts
PSF-convolved SMLM
ii
8000
6000
4000
2000
0
intensity / counts
Diffraction-limited / SMLM
iii
100
80
60
40
20
0
$z_{SMLM}$ / nm
IRM
iv
24000
22000
20000
18000
intensity / counts
IRM / Diffraction-limited
v
10000
7500
5000
2500
0
intensity / counts
IRM / SMLM
vi
100
80
60
40
20
0
$z_{SMLM}$ / nm
pix. SM brightness
vii
2500
2000
1500
1000
brightness / photons
pix. # of localizations
viii
10
8
6
4
2
localizations per pixel
pix. SMLM
ix
100
80
60
40
20
0
$z_{SMLM}$ / nm

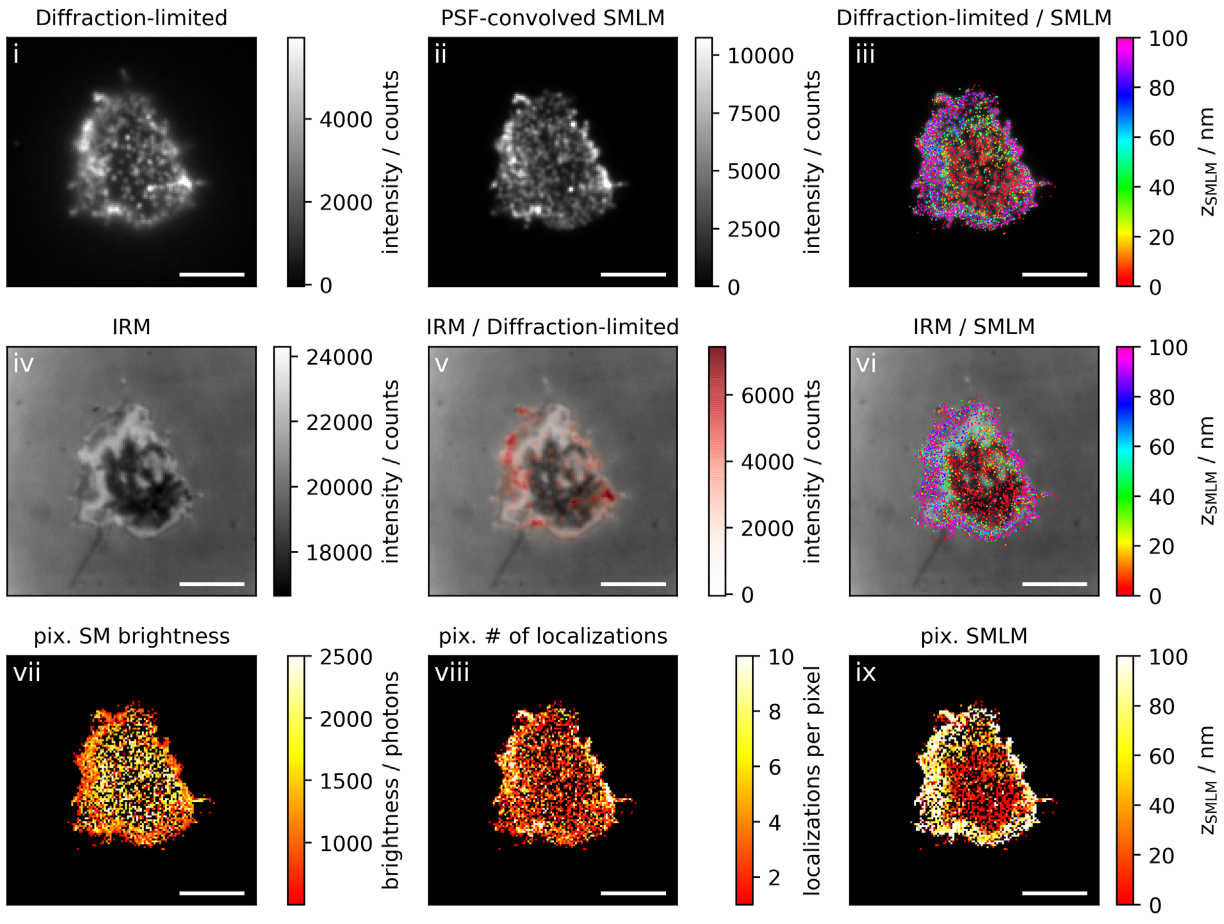

Diffraction-limited
i
4000
2000
0
intensity / counts
PSF-convolved SMLM
ii
10000
7500
5000
2500
0
intensity / counts
Diffraction-limited / SMLM
iii
100
80
60
40
20
0
$z_{SMLM}$ / nm
IRM
iv
24000
22000
20000
18000
intensity / counts
IRM / Diffraction-limited
v
6000
4000
2000
0
intensity / counts
IRM / SMLM
vi
100
80
60
40
20
0
$z_{SMLM}$ / nm
pix. SM brightness
vii
2500
2000
1500
1000
brightness / photons
pix. # of localizations
viii
10
8
6
4
2
localizations per pixel
pix. SMLM
ix
100
80
60
40
20
0
$z_{SMLM}$ / nm

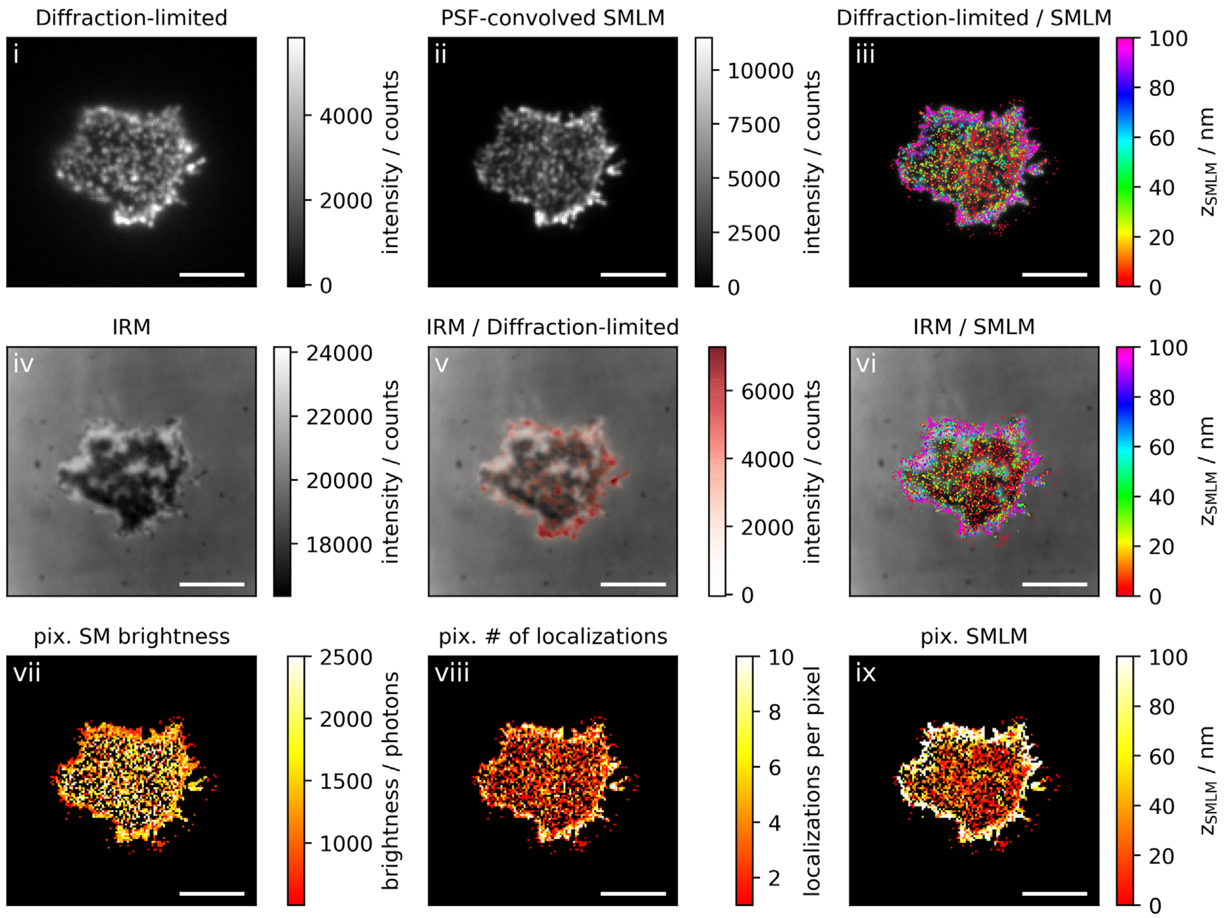
Diffraction-limited
i
4000
2000
0
intensity / counts
PSF-convolved SMLM
ii
10000
7500
5000
2500
0
intensity / counts
Diffraction-limited / SMLM
iii
100
80
60
40
20
0
$z_{SMLM}$ / nm
IRM
iv
24000
22000
20000
18000
intensity / counts
IRM / Diffraction-limited
v
6000
4000
2000
0
intensity / counts
IRM / SMLM
vi
100
80
60
40
20
0
$z_{SMLM}$ / nm
pix. SM brightness
vii
2500
2000
1500
1000
brightness / photons
pix. # of localizations
viii
10
8
6
4
2
localizations per pixel
pix. SMLM
ix
100
80
60
40
20
0
$z_{SMLM}$ / nm

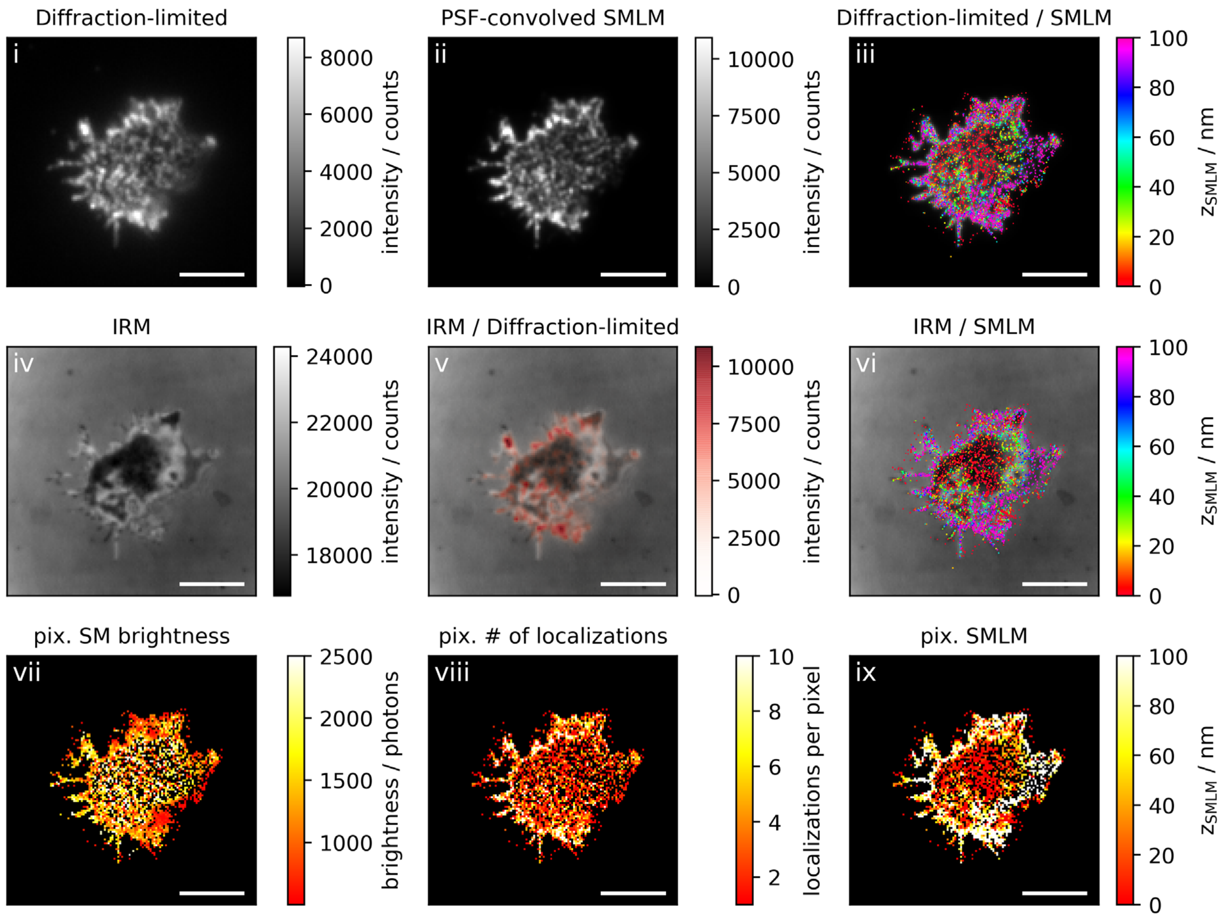

Diffraction-limited
i
intensity / counts
PSF-convolved SMLM
ii
Diffraction-limited / SMLM
iii
z_SMLM / nm
IRM
iv
IRM / Diffraction-limited
v
IRM / SMLM
vi
pix. SM brightness
vii
brightness / photons
pix. # of localizations
viii
localizations per pixel
pix. SMLM
ix

**Figure 3b: Non-activating conditions, high ICAM-1 density, fixation: 10 min post seeding**

**Correlative 3D-SMLM, IRM, and diffraction-limited TIR microscopy of the immunological synapse.** T cells were seeded on an SLB functionalized with high density of ICAM-1, and fixed 10 minutes post seeding. The T cell was imaged with IRM and fluorescence microscopy: (i) Diffraction-limited TIR image of the T cell. (ii) Reconstruction of the diffraction-limited image by convolving the 3D-SMLM image with the corresponding psf. (iii) Overlay of the diffraction-limited TIR image with the 3D-SMLM image. Color-code indicates distance to the coverslip $z_{SMLM}$. (iv) IRM image. (v) Overlay of the IRM image with the diffraction-limited image. (vi) Overlay of the IRM and the 3D-SMLM image. Bottom row images were generated by calculating the pixel-wise average of the 3D-SMLM images (pixel size of 146nm is consistent with diffraction-limited image) according to pixelated mean single molecule (SM) intensity (vii), pixelated number of localizations (viii) and pixelated mean $z_{SMLM}$ (ix). Scale bars 5 µm.

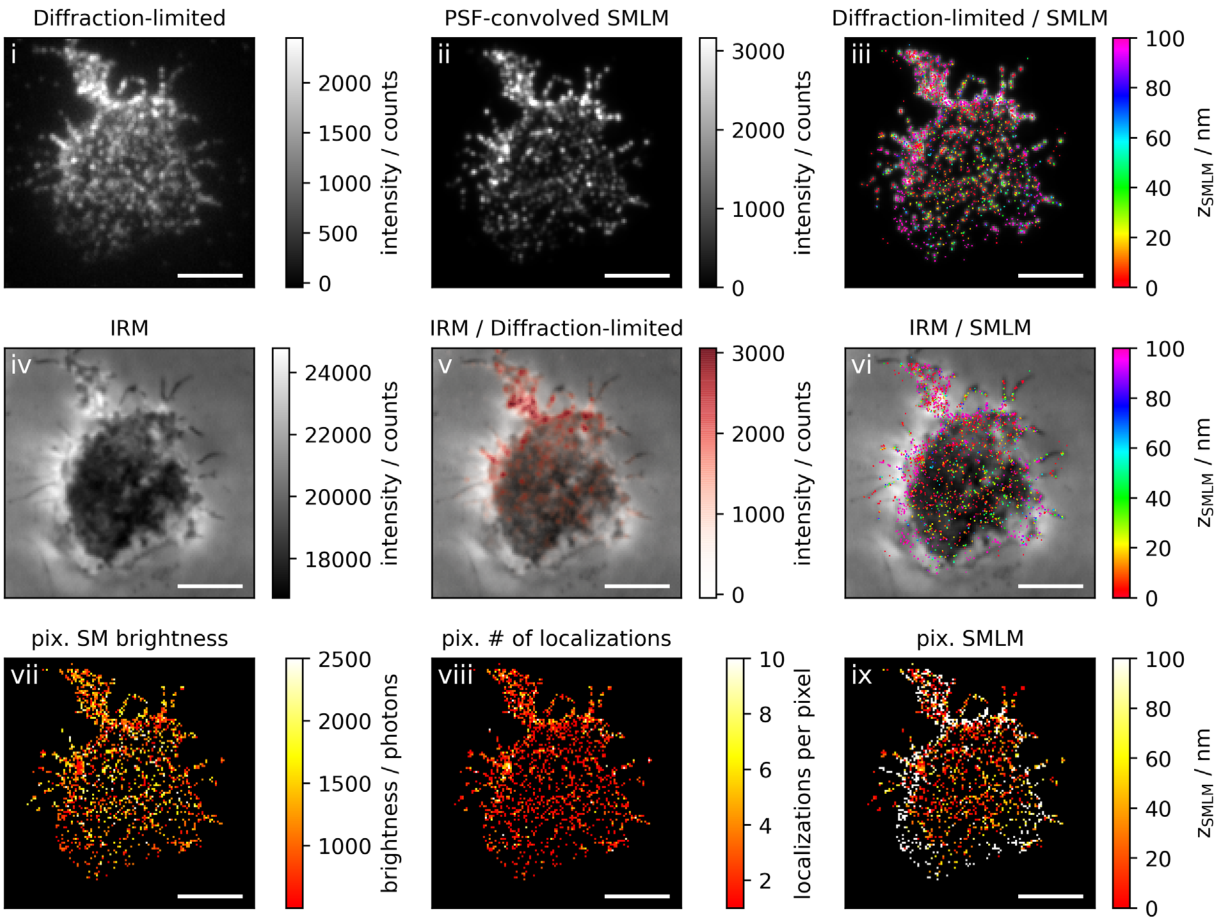
Diffraction-limited
i
intensity / counts
0
500
1000
1500
2000
PSF-convolved SMLM
ii
intensity / counts
0
1000
2000
3000
Diffraction-limited / SMLM
iii
$z_{SMLM}$ / nm
0
20
40
60
80
100
IRM
iv
intensity / counts
18000
20000
22000
24000
IRM / Diffraction-limited
v
intensity / counts
0
1000
2000
3000
IRM / SMLM
vi
$z_{SMLM}$ / nm
0
20
40
60
80
100
pix. SM brightness
vii
brightness / photons
1000
1500
2000
2500
pix. # of localizations
viii
localizations per pixel
2
4
6
8
10
pix. SMLM
ix
$z_{SMLM}$ / nm
0
20
40
60
80
100

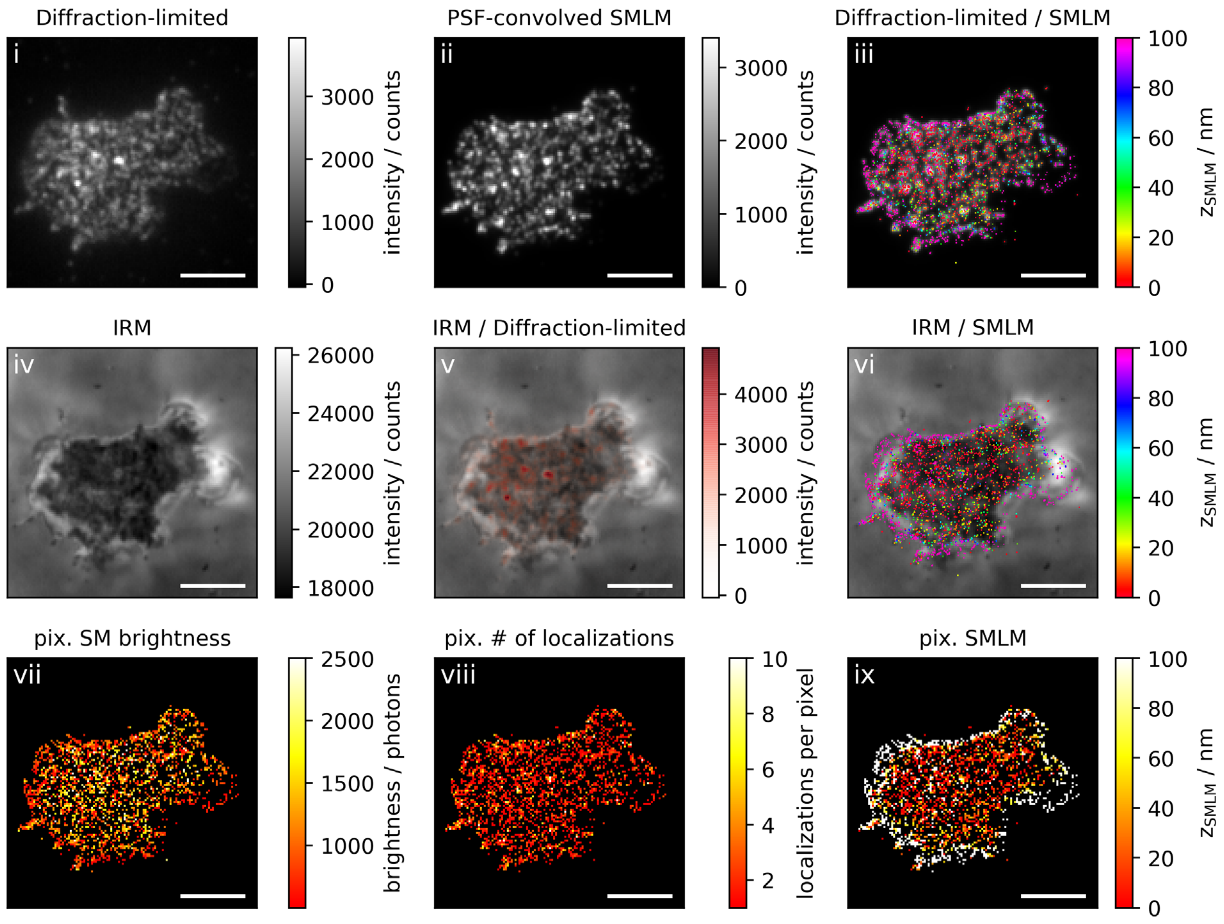
Diffraction-limited
i
intensity / counts
PSF-convolved SMLM
ii
intensity / counts
Diffraction-limited / SMLM
iii
$z_{SMLM}$ / nm
IRM
iv
intensity / counts
IRM / Diffraction-limited
v
intensity / counts
IRM / SMLM
vi
$z_{SMLM}$ / nm
pix. SM brightness
vii
brightness / photons
pix. # of localizations
viii
localizations per pixel
pix. SMLM
ix
$z_{SMLM}$ / nm

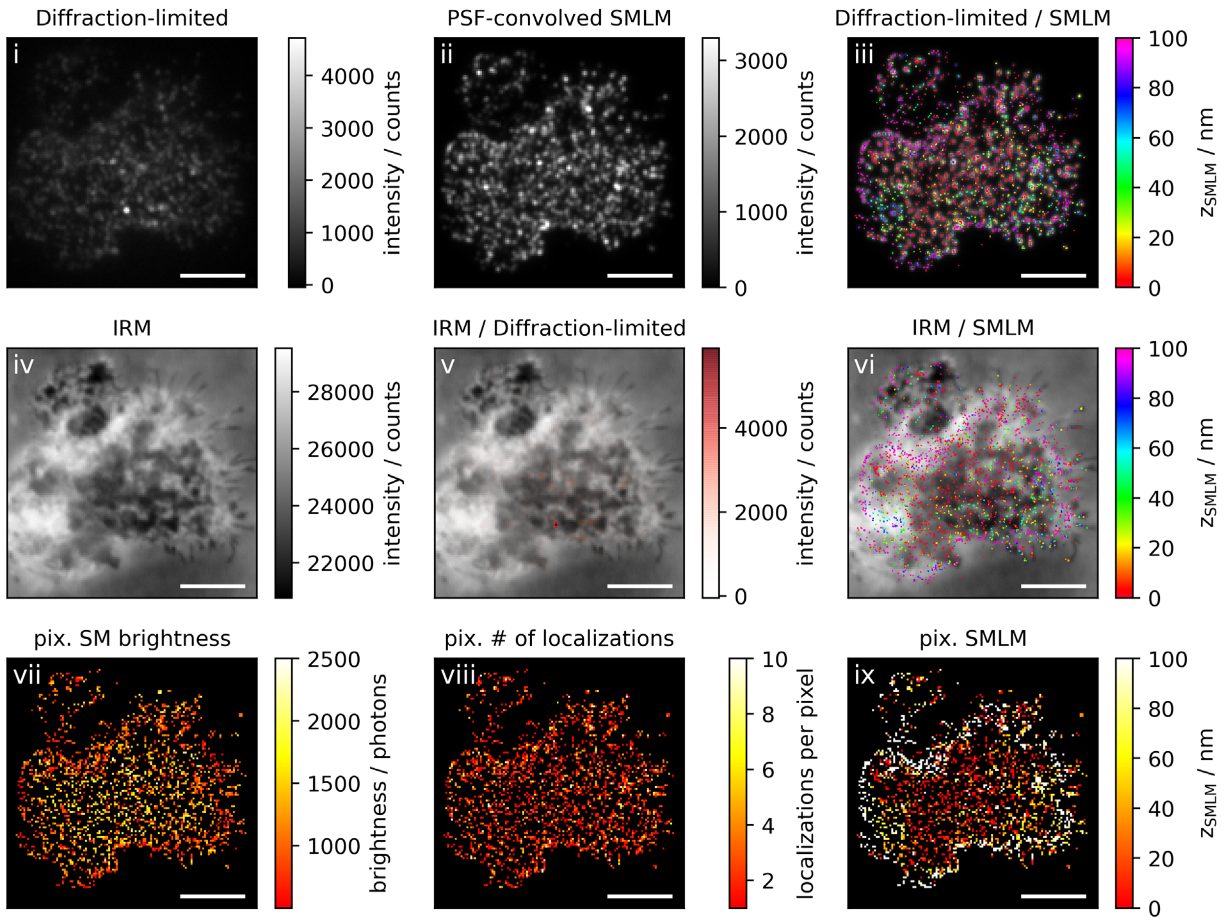
Diffraction-limited
i
intensity / counts
PSF-convolved SMLM
ii
Diffraction-limited / SMLM
iii
$z_{SMLM}$ / nm
IRM
iv
IRM / Diffraction-limited
v
IRM / SMLM
vi
pix. SM brightness
vii
brightness / photons
pix. # of localizations
viii
localizations per pixel
pix. SMLM
ix

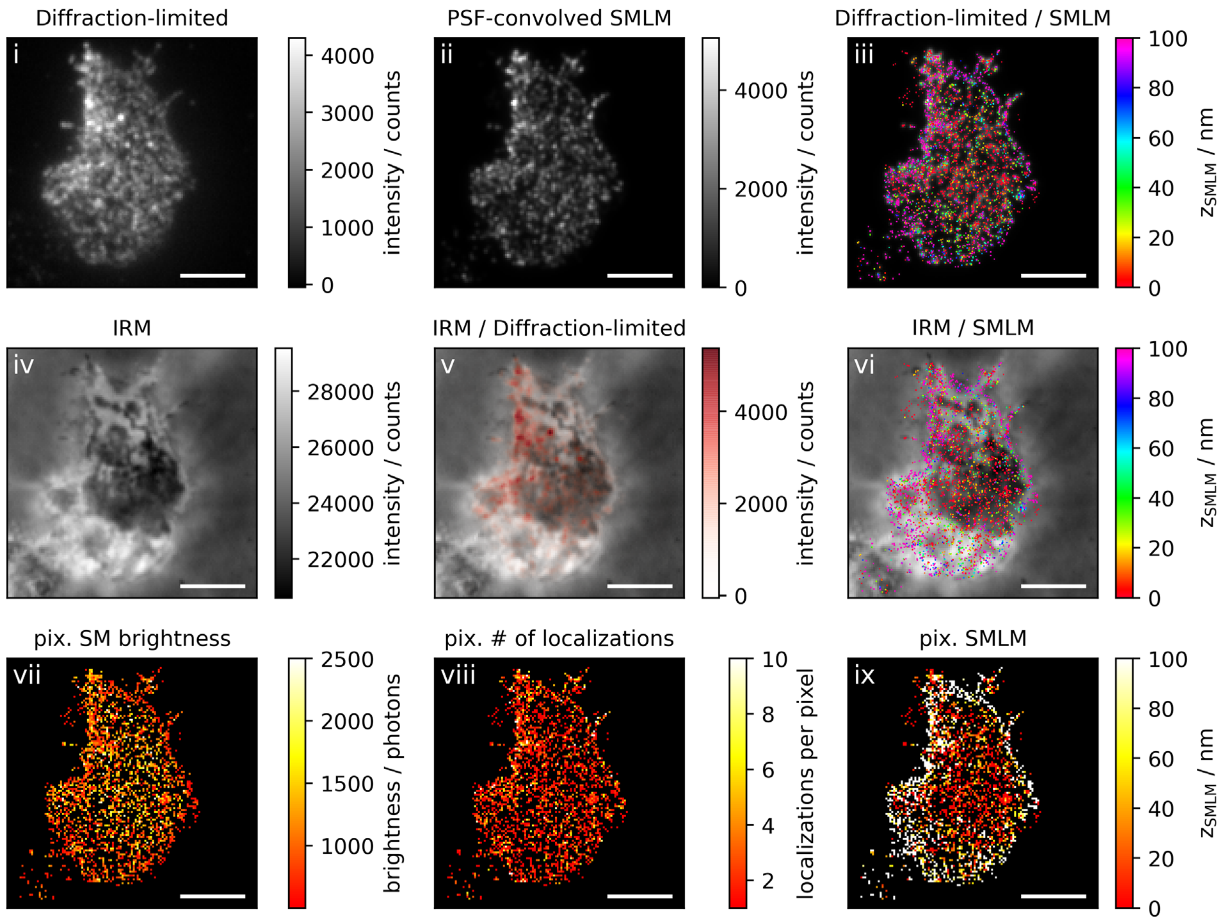
Diffraction-limited
i
intensity / counts
4000
3000
2000
1000
0
PSF-convolved SMLM
ii
intensity / counts
4000
2000
0
Diffraction-limited / SMLM
iii
$z_{SMLM}$ / nm
100
80
60
40
20
0
IRM
iv
intensity / counts
28000
26000
24000
22000
IRM / Diffraction-limited
v
intensity / counts
4000
2000
0
IRM / SMLM
vi
$z_{SMLM}$ / nm
100
80
60
40
20
0
pix. SM brightness
vii
brightness / photons
2500
2000
1500
1000
pix. # of localizations
viii
localizations per pixel
10
8
6
4
2
pix. SMLM
ix
$z_{SMLM}$ / nm
100
80
60
40
20
0

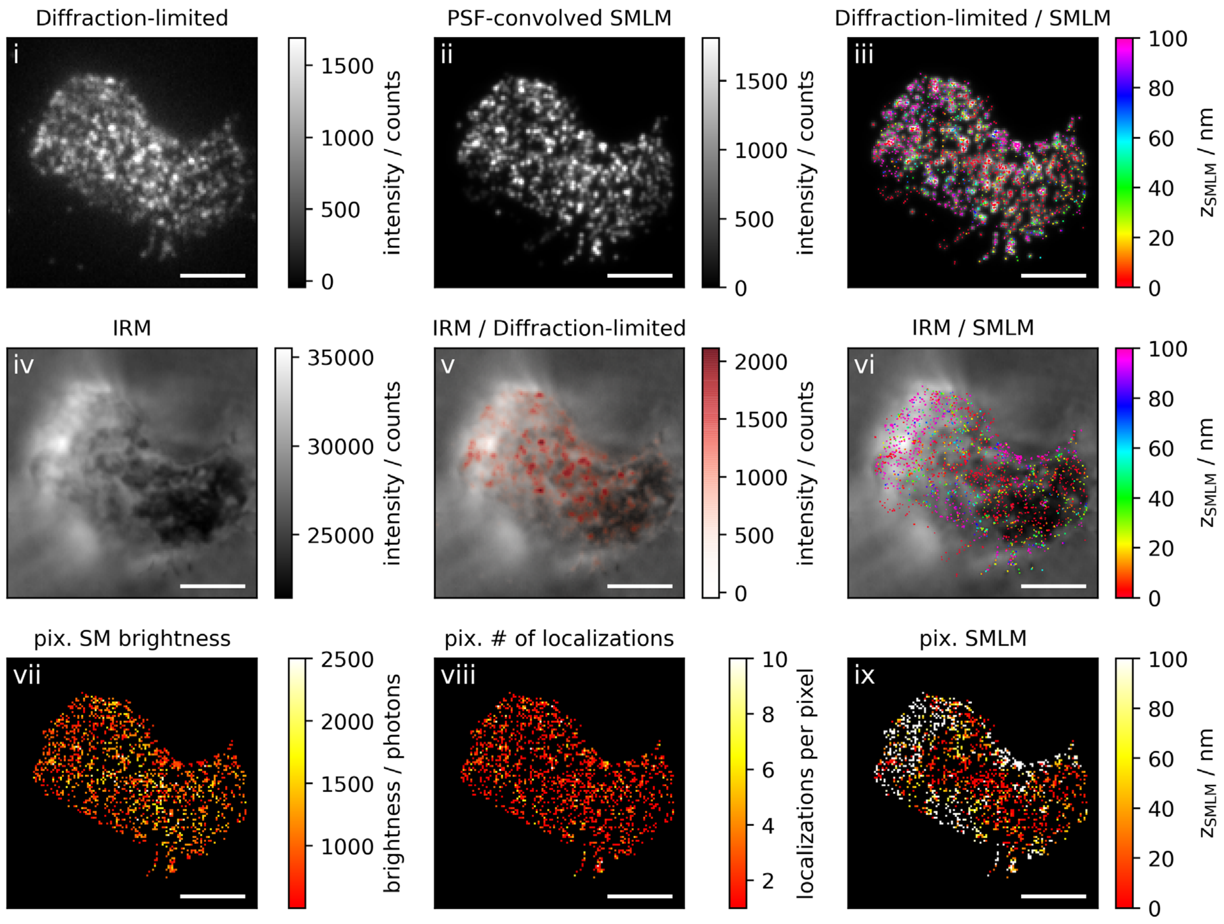

Diffraction-limited
i
intensity / counts
PSF-convolved SMLM
ii
Diffraction-limited / SMLM
iii
$z_{SMLM}$ / nm
IRM
iv
IRM / Diffraction-limited
v
IRM / SMLM
vi
pix. SM brightness
vii
brightness / photons
pix. # of localizations
viii
localizations per pixel
pix. SMLM
ix

**Figure 3c: Non-activating conditions, high ICAM-1 density, fixation: 10-15 min post seeding**

**Correlative 3D-SMLM, IRM, and diffraction-limited TIR microscopy of the immunological synapse.** T cells were seeded on an SLB functionalized with high density of ICAM-1, and fixed 10-15 minutes post seeding. The T cell was imaged with IRM and fluorescence microscopy: (i) Diffraction-limited TIR image of the T cell. (ii) Reconstruction of the diffraction-limited image by convolving the 3D-SMLM image with the corresponding psf. (iii) Overlay of the diffraction-limited TIR image with the 3D-SMLM image. Color-code indicates distance to the coverslip $z_{SMLM}$. (iv) IRM image. (v) Overlay of the IRM image with the diffraction-limited image. (vi) Overlay of the IRM and the 3D-SMLM image. Bottom row images were generated by calculating the pixel-wise average of the 3D-SMLM images (pixel size of 146nm is consistent with diffraction-limited image) according to pixelated mean single molecule (SM) intensity (vii), pixelated number of localizations (viii) and pixelated mean $z_{SMLM}$ (ix). Scale bars 5 µm.

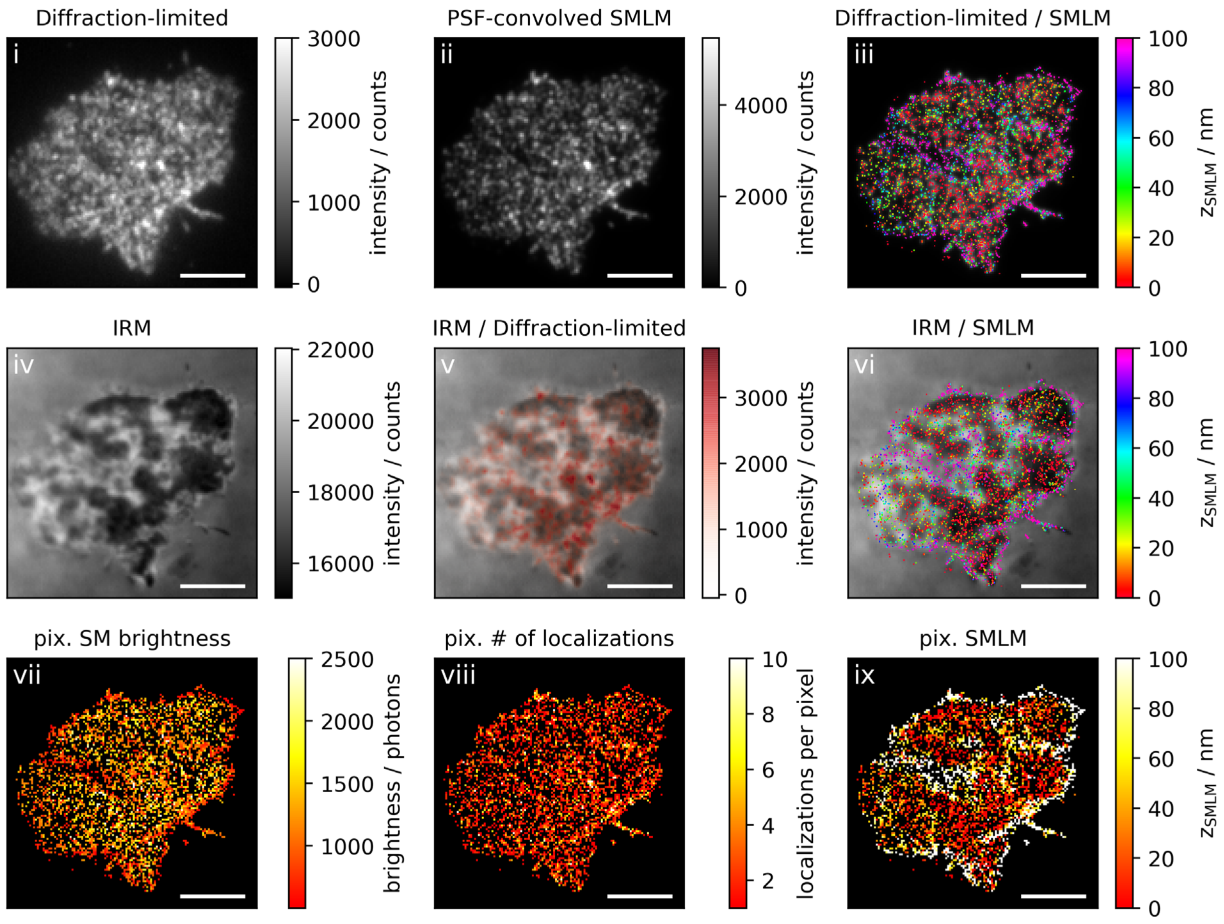

Diffraction-limited
i
intensity / counts
PSF-convolved SMLM
ii
Diffraction-limited / SMLM
iii
$z_{SMLM}$ / nm
IRM
iv
IRM / Diffraction-limited
v
IRM / SMLM
vi
pix. SM brightness
vii
brightness / photons
pix. # of localizations
viii
localizations per pixel
pix. SMLM
ix

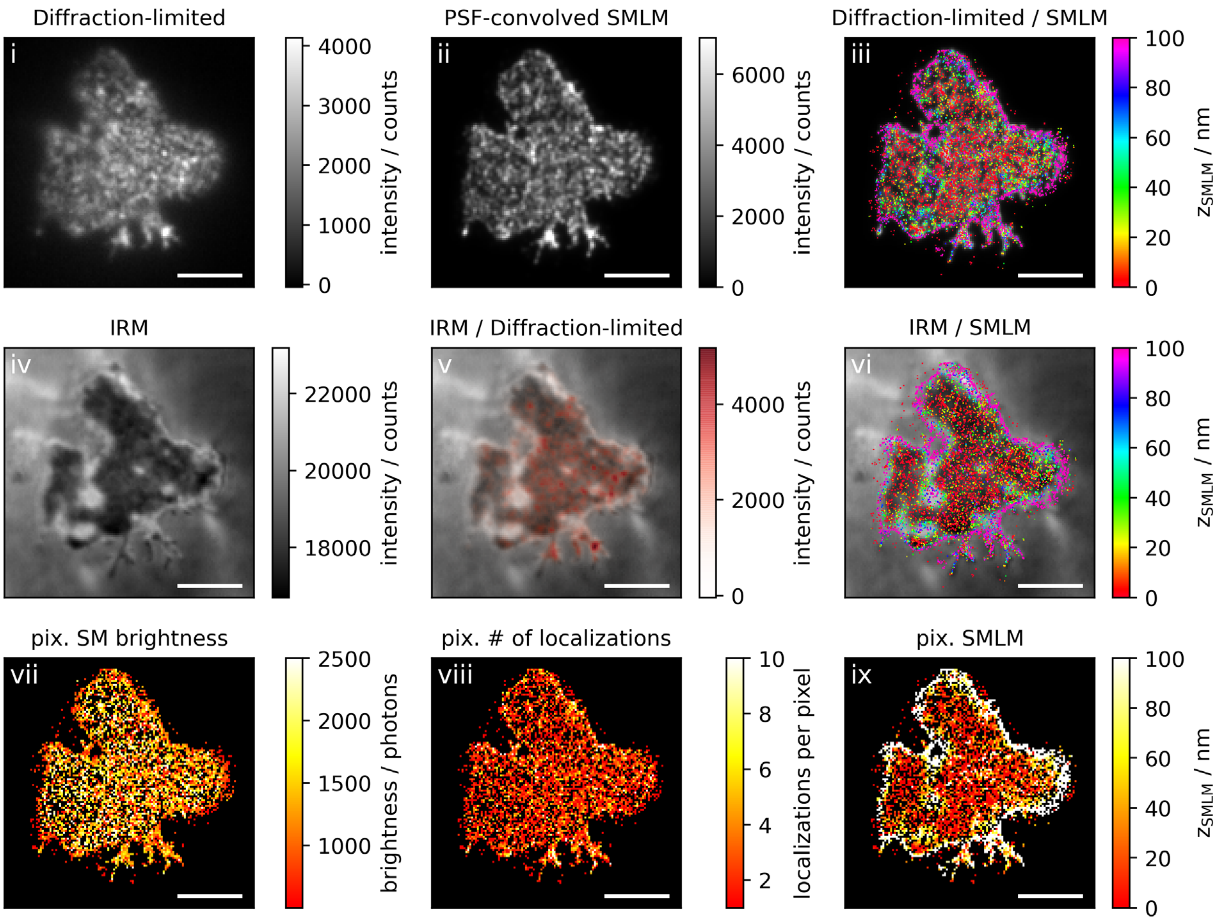
Diffraction-limited
i
4000
3000
2000
1000
0
intensity / counts
PSF-convolved SMLM
ii
6000
4000
2000
0
intensity / counts
Diffraction-limited / SMLM
iii
100
80
60
40
20
0
$z_{SMLM}$ / nm
IRM
iv
22000
20000
18000
intensity / counts
IRM / Diffraction-limited
v
4000
2000
0
intensity / counts
IRM / SMLM
vi
100
80
60
40
20
0
$z_{SMLM}$ / nm
pix. SM brightness
vii
2500
2000
1500
1000
brightness / photons
pix. # of localizations
viii
10
8
6
4
2
localizations per pixel
pix. SMLM
ix
100
80
60
40
20
0
$z_{SMLM}$ / nm

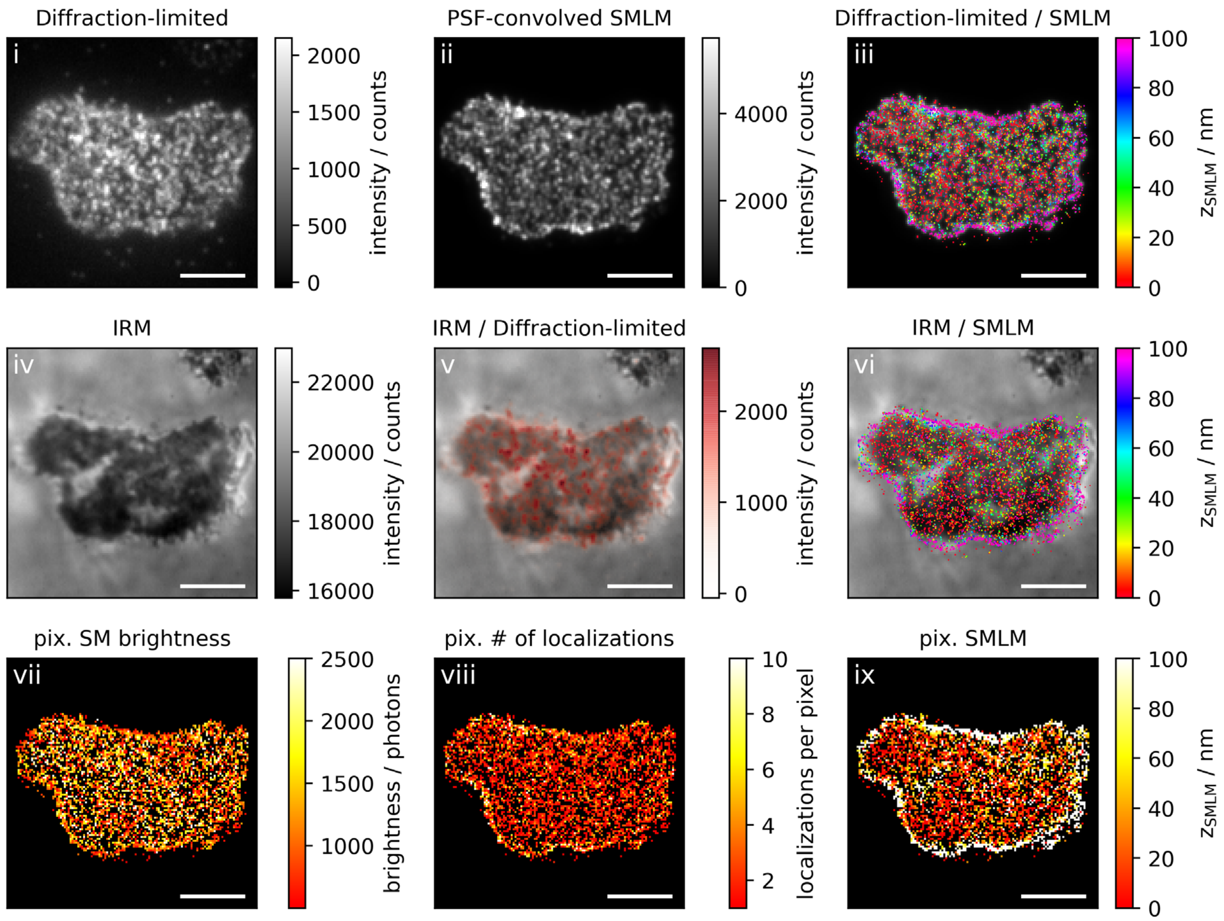
Diffraction-limited
i
intensity / counts
PSF-convolved SMLM
ii
intensity / counts
Diffraction-limited / SMLM
iii
$z_{SMLM}$ / nm
IRM
iv
intensity / counts
IRM / Diffraction-limited
v
intensity / counts
IRM / SMLM
vi
$z_{SMLM}$ / nm
pix. SM brightness
vii
brightness / photons
pix. # of localizations
viii
localizations per pixel
pix. SMLM
ix
$z_{SMLM}$ / nm

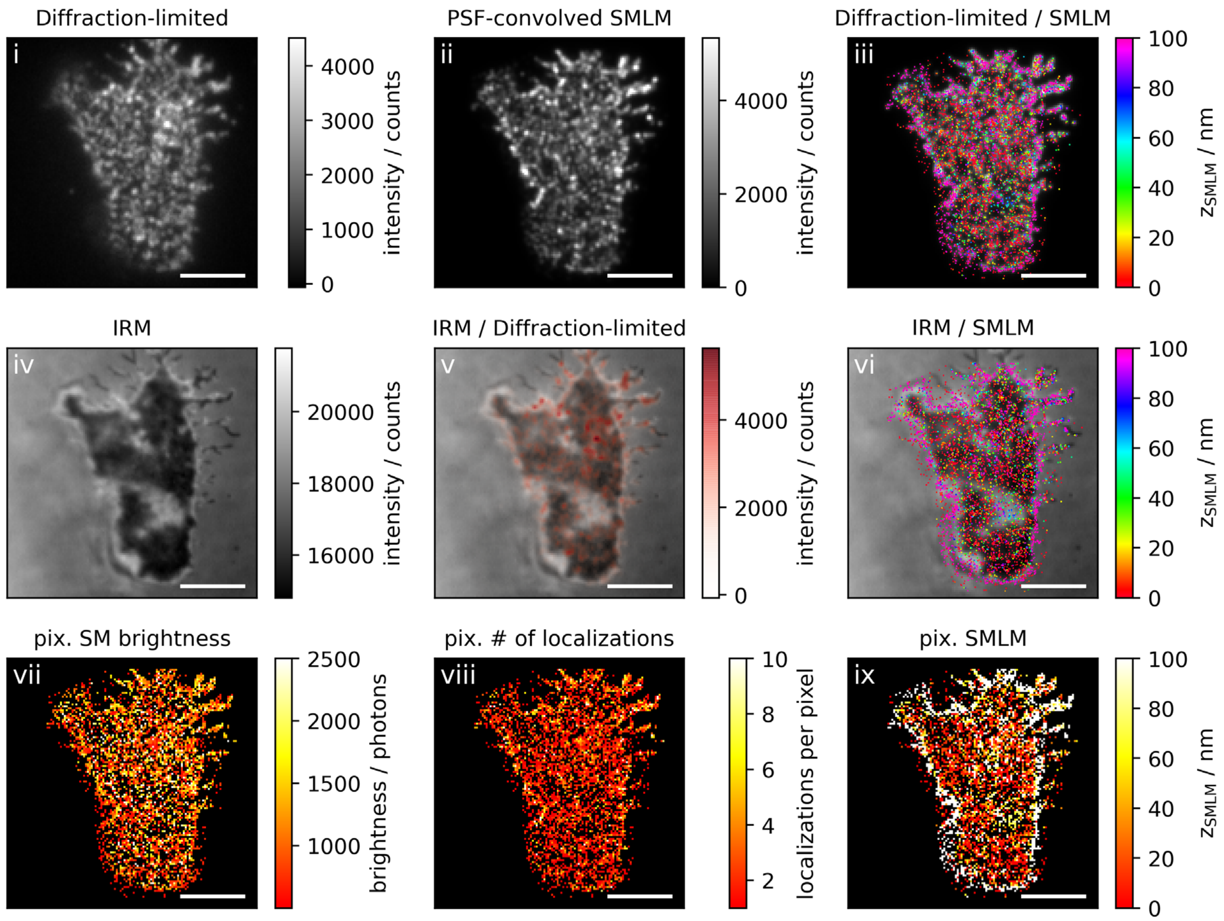

Diffraction-limited
i
4000
3000
2000
1000
0
intensity / counts
PSF-convolved SMLM
ii
4000
2000
0
intensity / counts
Diffraction-limited / SMLM
iii
100
80
60
40
20
0
$z_{SMLM}$ / nm
IRM
iv
20000
18000
16000
intensity / counts
IRM / Diffraction-limited
v
4000
2000
0
intensity / counts
IRM / SMLM
vi
100
80
60
40
20
0
$z_{SMLM}$ / nm
pix. SM brightness
vii
2500
2000
1500
1000
brightness / photons
pix. # of localizations
viii
10
8
6
4
2
localizations per pixel
pix. SMLM
ix
100
80
60
40
20
0
$z_{SMLM}$ / nm

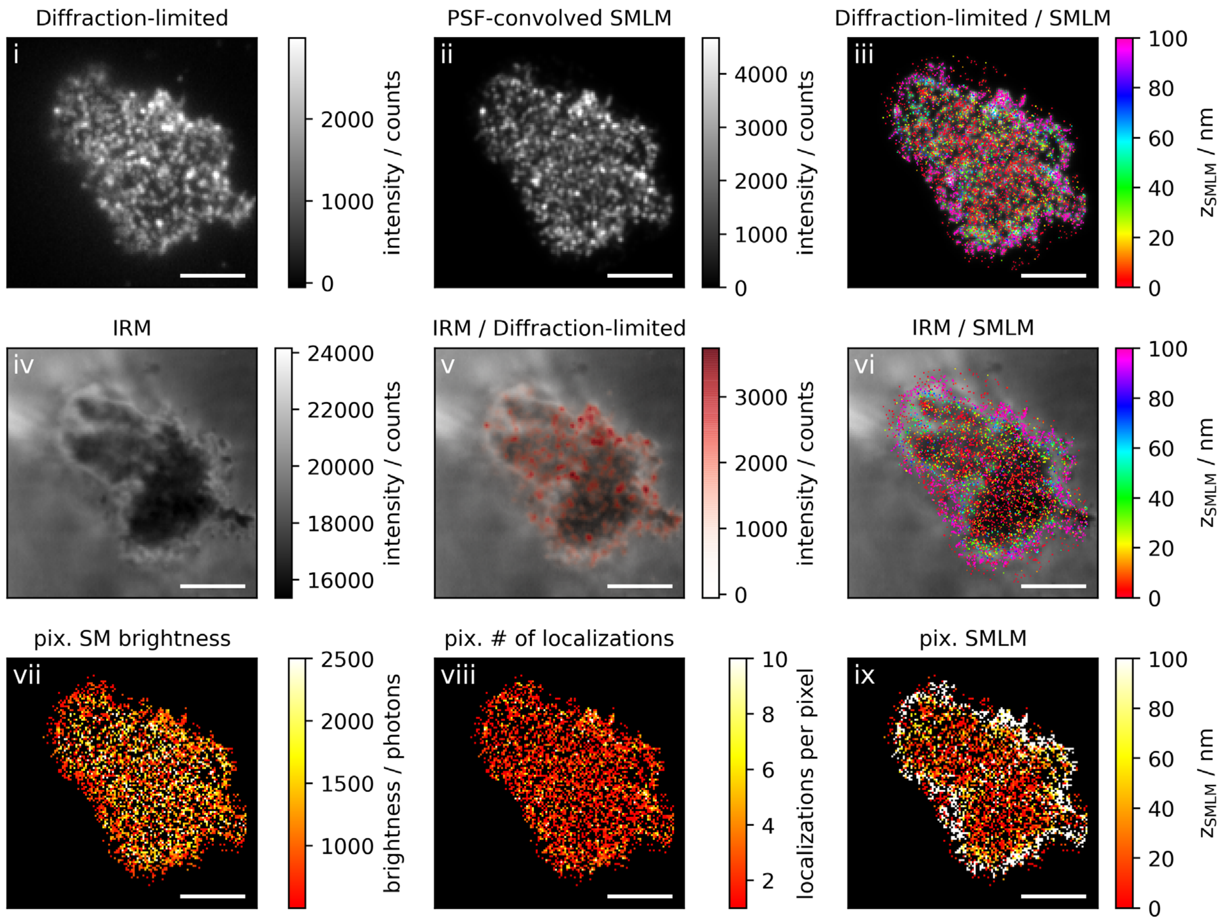
Diffraction-limited
i
intensity / counts
PSF-convolved SMLM
ii
intensity / counts
Diffraction-limited / SMLM
iii
$z_{SMLM}$ / nm
IRM
iv
intensity / counts
IRM / Diffraction-limited
v
intensity / counts
IRM / SMLM
vi
$z_{SMLM}$ / nm
pix. SM brightness
vii
brightness / photons
pix. # of localizations
viii
localizations per pixel
pix. SMLM
ix
$z_{SMLM}$ / nm

**Figure 4a: Non-activating conditions, low ICAM-1 density, fixation: 5-10 min post seeding**

**Correlative 3D-SMLM, IRM, and diffraction-limited TIR microscopy of the immunological synapse.** T cells were seeded on an SLB functionalized with low density of ICAM-1, and fixed 5-10 minutes post seeding. The T cell was imaged with IRM and fluorescence microscopy: (i) Diffraction-limited TIR image of the T cell. (ii) Reconstruction of the diffraction-limited image by convolving the 3D-SMLM image with the corresponding psf. (iii) Overlay of the diffraction-limited TIR image with the 3D-SMLM image. Color-code indicates distance to the coverslip $z_{SMLM}$. (iv) IRM image. (v) Overlay of the IRM image with the diffraction-limited image. (vi) Overlay of the IRM and the 3D-SMLM image. Bottom row images were generated by calculating the pixel-wise average of the 3D-SMLM images (pixel size of 146nm is consistent with diffraction-limited image) according to pixelated mean single molecule (SM) intensity (vii), pixelated number of localizations (viii) and pixelated mean $z_{SMLM}$ (ix). Scale bars 5 µm.

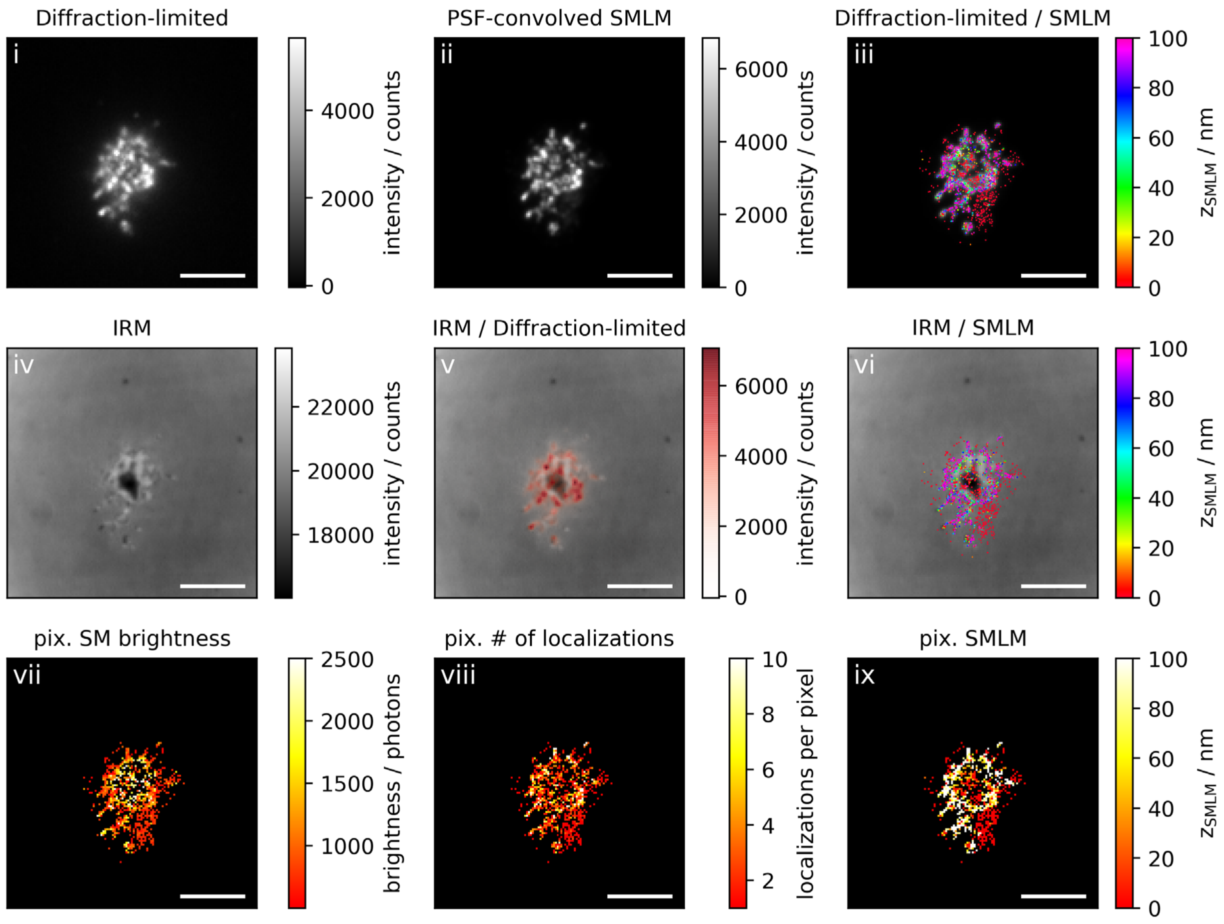

Diffraction-limited
i
intensity / counts
0
2000
4000
PSF-convolved SMLM
ii
intensity / counts
0
2000
4000
6000
Diffraction-limited / SMLM
iii
$z_{SMLM}$ / nm
0
20
40
60
80
100
IRM
iv
intensity / counts
18000
20000
22000
IRM / Diffraction-limited
v
intensity / counts
0
2000
4000
6000
IRM / SMLM
vi
$z_{SMLM}$ / nm
0
20
40
60
80
100
pix. SM brightness
vii
brightness / photons
1000
1500
2000
2500
pix. # of localizations
viii
localizations per pixel
2
4
6
8
10
pix. SMLM
ix
$z_{SMLM}$ / nm
0
20
40
60
80
100

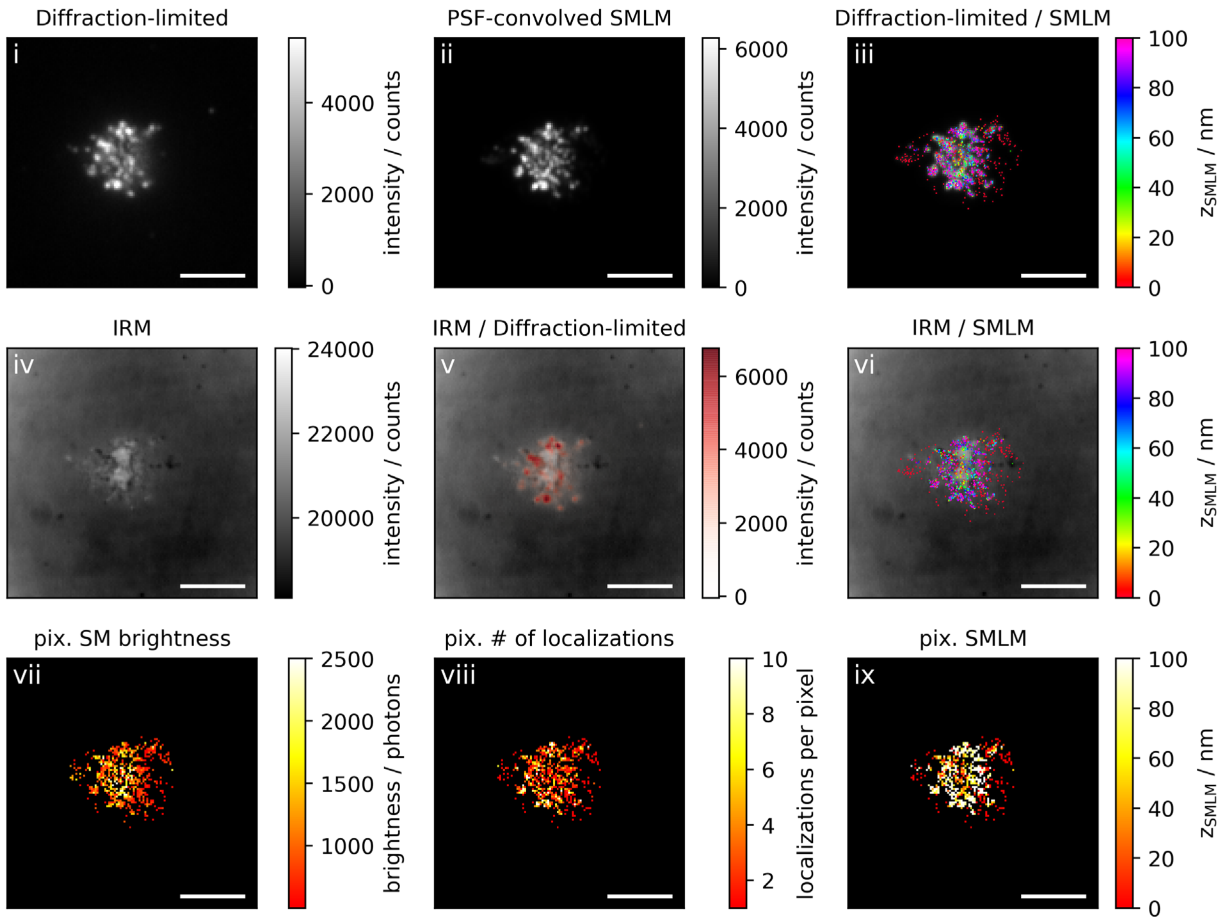

Diffraction-limited
i
4000
2000
0
intensity / counts
PSF-convolved SMLM
ii
6000
4000
2000
0
intensity / counts
Diffraction-limited / SMLM
iii
100
80
60
40
20
0
z_SMLM / nm
IRM
iv
24000
22000
20000
intensity / counts
IRM / Diffraction-limited
v
6000
4000
2000
0
intensity / counts
IRM / SMLM
vi
100
80
60
40
20
0
z_SMLM / nm
pix. SM brightness
vii
2500
2000
1500
1000
brightness / photons
pix. # of localizations
viii
10
8
6
4
2
localizations per pixel
pix. SMLM
ix
100
80
60
40
20
0
z_SMLM / nm

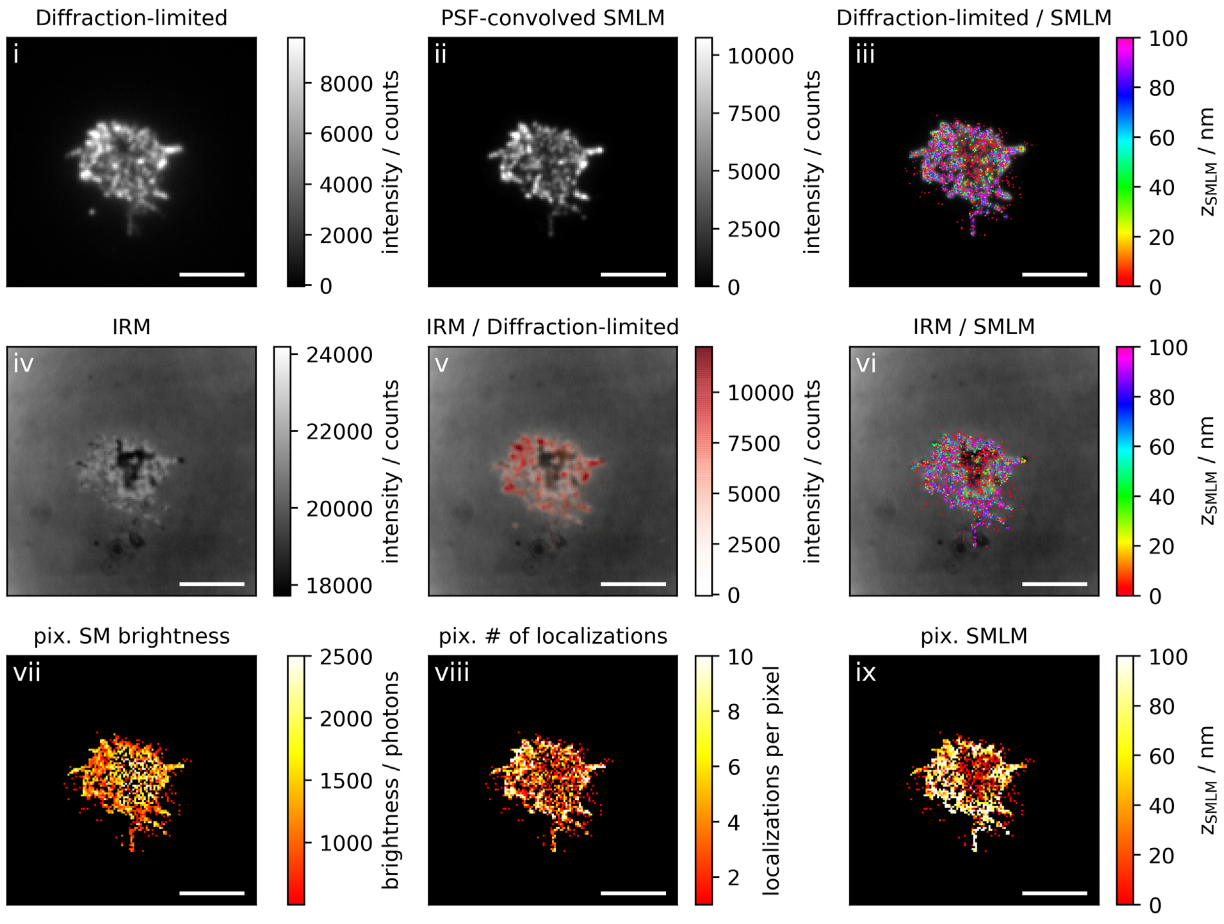
Diffraction-limited
i
intensity / counts
PSF-convolved SMLM
ii
Diffraction-limited / SMLM
iii
$z_{SMLM}$ / nm
IRM
iv
IRM / Diffraction-limited
v
IRM / SMLM
vi
pix. SM brightness
vii
brightness / photons
pix. # of localizations
viii
localizations per pixel
pix. SMLM
ix

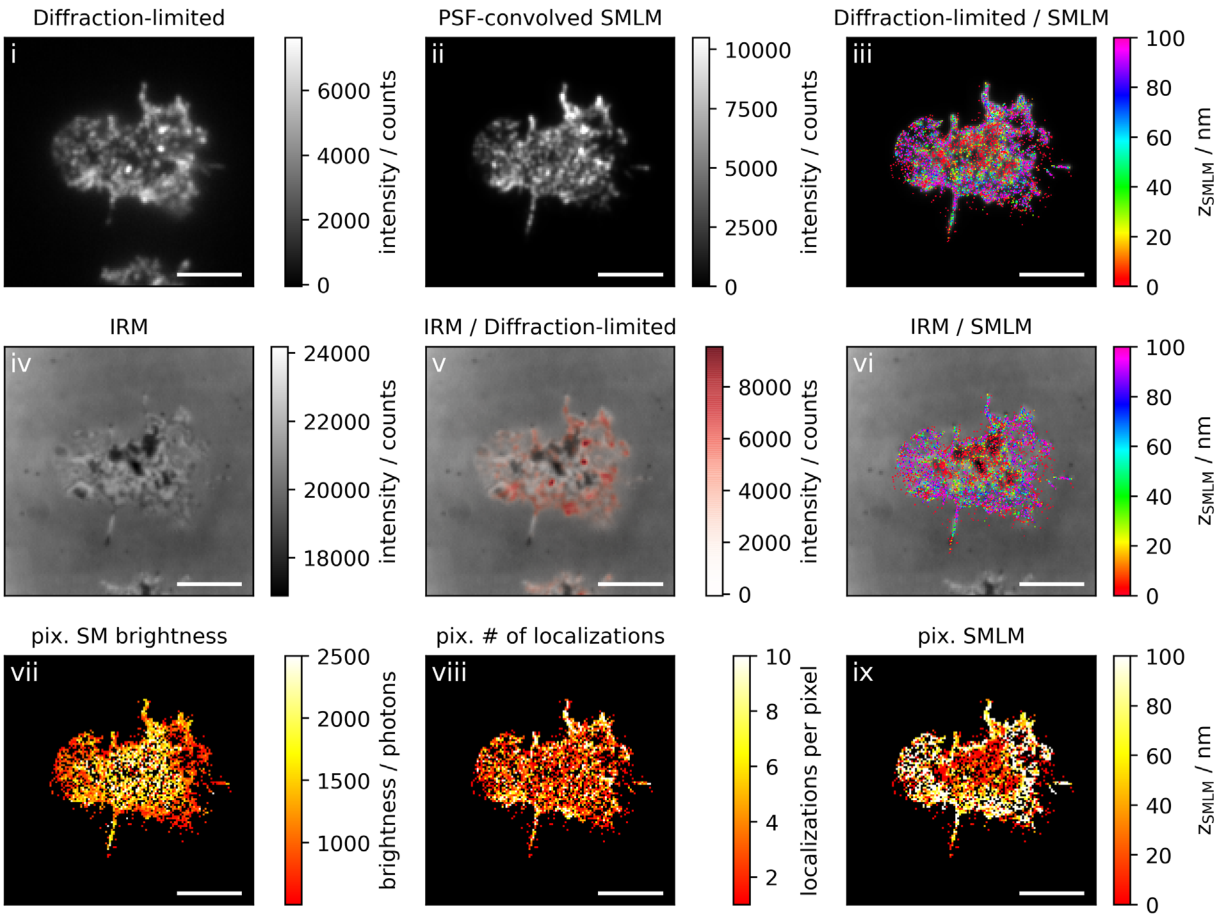
Diffraction-limited
i
6000
4000
2000
0
intensity / counts
PSF-convolved SMLM
ii
10000
7500
5000
2500
0
intensity / counts
Diffraction-limited / SMLM
iii
100
80
60
40
20
0
zSMLM / nm
IRM
iv
24000
22000
20000
18000
intensity / counts
IRM / Diffraction-limited
v
8000
6000
4000
2000
0
intensity / counts
IRM / SMLM
vi
100
80
60
40
20
0
zSMLM / nm
pix. SM brightness
vii
2500
2000
1500
1000
brightness / photons
pix. # of localizations
viii
10
8
6
4
2
localizations per pixel
pix. SMLM
ix
100
80
60
40
20
0
zSMLM / nm

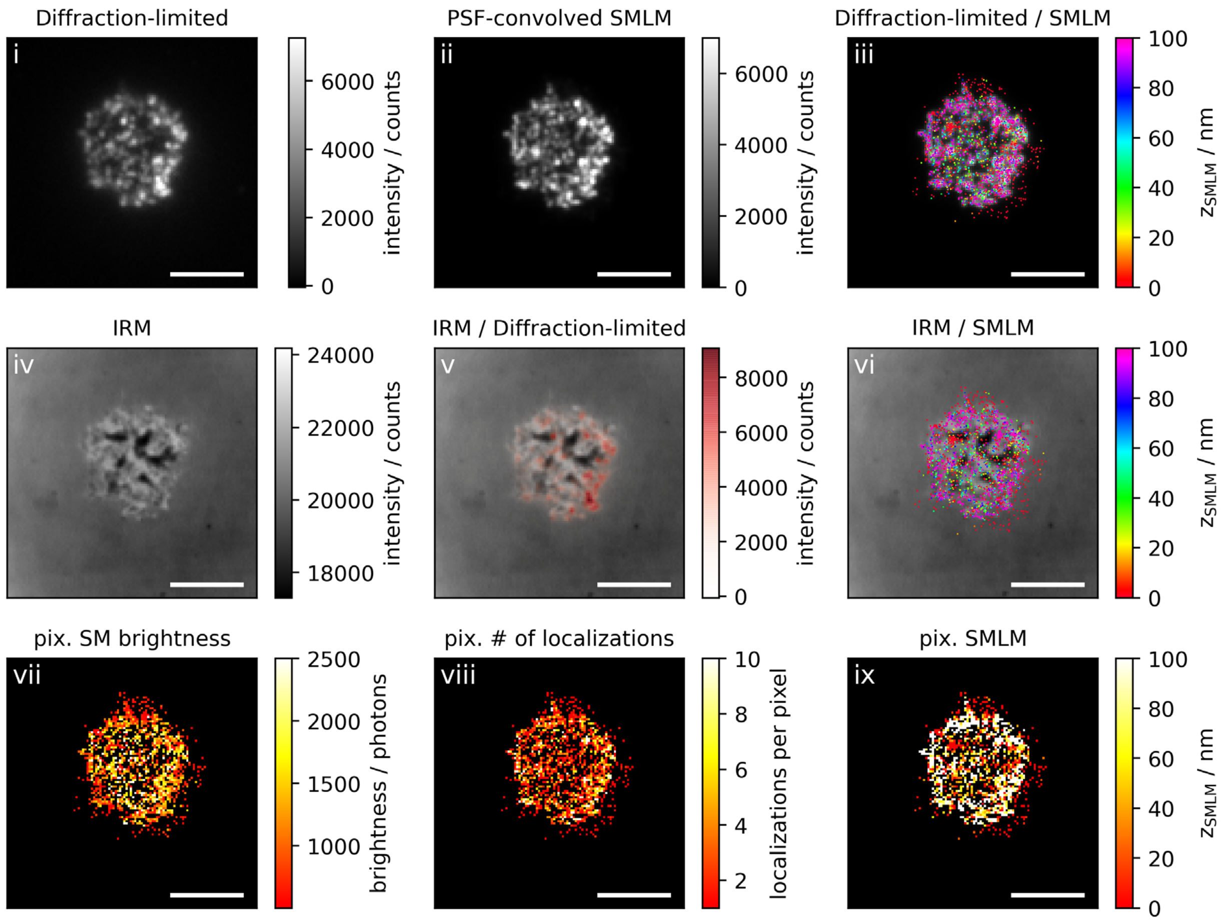

Diffraction-limited
i
intensity / counts
PSF-convolved SMLM
ii
Diffraction-limited / SMLM
iii
$z_{SMLM}$ / nm
IRM
iv
IRM / Diffraction-limited
v
IRM / SMLM
vi
pix. SM brightness
vii
brightness / photons
pix. # of localizations
viii
localizations per pixel
pix. SMLM
ix

**Figure 4b: Non-activating conditions, low ICAM-1 density, fixation: 10 min post seeding**

**Correlative 3D-SMLM, IRM, and diffraction-limited TIR microscopy of the immunological synapse.** T cells were seeded on an SLB functionalized with low density of ICAM-1, and fixed 10 minutes post seeding. The T cell was imaged with IRM and fluorescence microscopy: (i) Diffraction-limited TIR image of the T cell. (ii) Reconstruction of the diffraction-limited image by convolving the 3D-SMLM image with the corresponding psf. (iii) Overlay of the diffraction-limited TIR image with the 3D-SMLM image. Color-code indicates distance to the coverslip $z_{SMLM}$. (iv) IRM image. (v) Overlay of the IRM image with the diffraction-limited image. (vi) Overlay of the IRM and the 3D-SMLM image. Bottom row images were generated by calculating the pixel-wise average of the 3D-SMLM images (pixel size of 146nm is consistent with diffraction-limited image) according to pixelated mean single molecule (SM) intensity (vii), pixelated number of localizations (viii) and pixelated mean $z_{SMLM}$ (ix). Scale bars 5 µm.

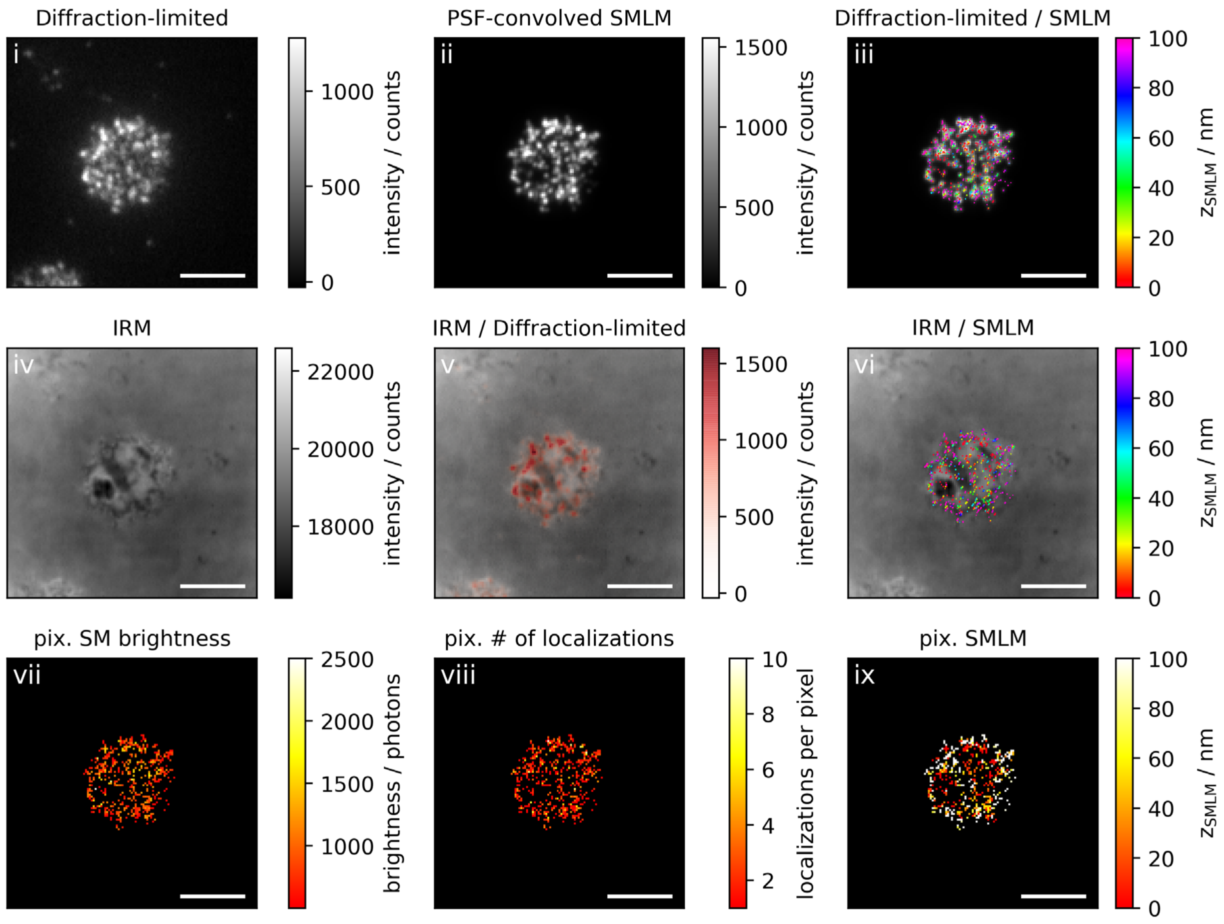

Diffraction-limited
i
1000
500
0
intensity / counts
PSF-convolved SMLM
ii
1500
1000
500
0
intensity / counts
Diffraction-limited / SMLM
iii
100
80
60
40
20
0
$z_{SMLM}$ / nm
IRM
iv
22000
20000
18000
intensity / counts
IRM / Diffraction-limited
v
1500
1000
500
0
intensity / counts
IRM / SMLM
vi
100
80
60
40
20
0
$z_{SMLM}$ / nm
pix. SM brightness
vii
2500
2000
1500
1000
brightness / photons
pix. # of localizations
viii
10
8
6
4
2
localizations per pixel
pix. SMLM
ix
100
80
60
40
20
0
$z_{SMLM}$ / nm

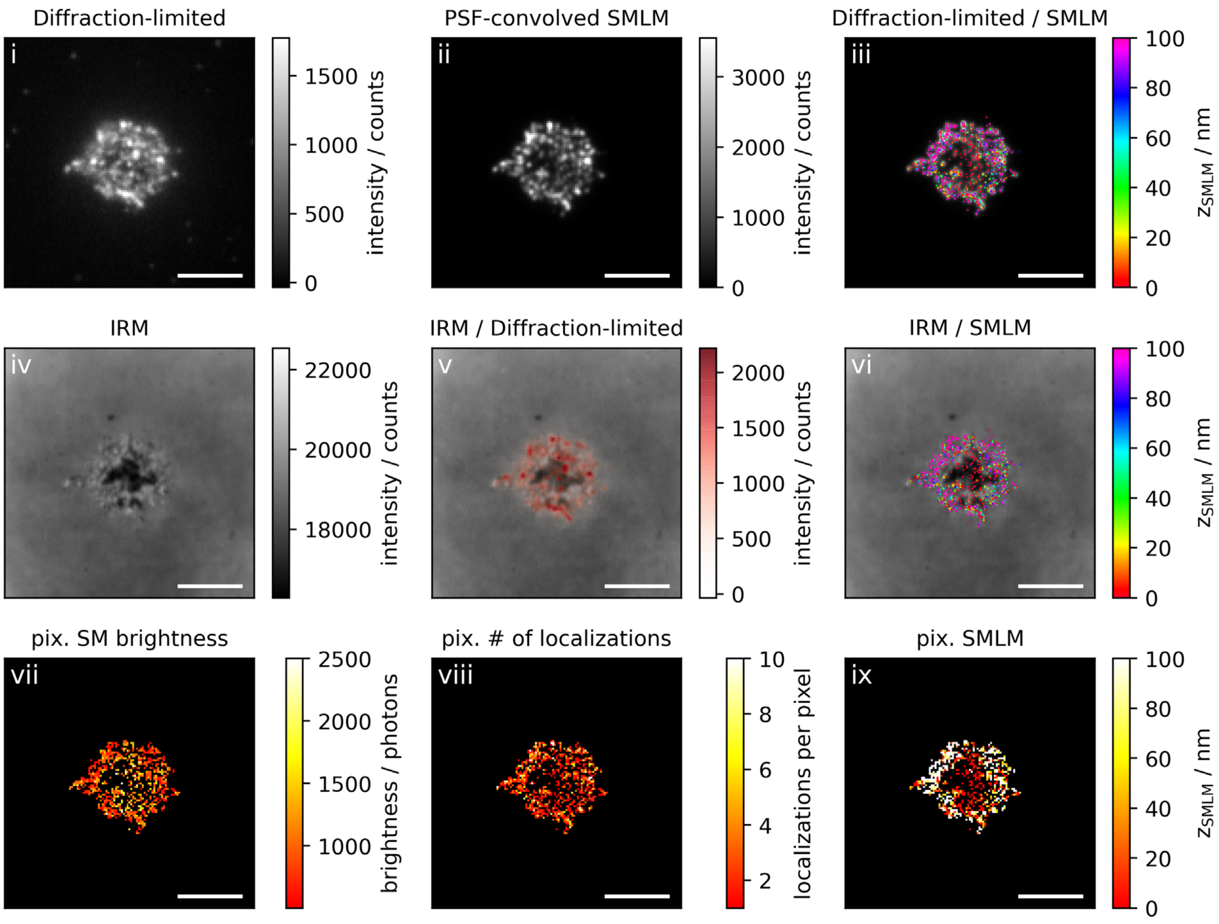

Diffraction-limited
PSF-convolved SMLM
Diffraction-limited / SMLM
i
ii
iii
1500
1000
500
0
intensity / counts
3000
2000
1000
0
intensity / counts
100
80
60
40
20
0
z_SMLM / nm
IRM
IRM / Diffraction-limited
IRM / SMLM
iv
v
vi
22000
20000
18000
intensity / counts
2000
1500
1000
500
0
intensity / counts
100
80
60
40
20
0
z_SMLM / nm
pix. SM brightness
pix. # of localizations
pix. SMLM
vii
viii
ix
2500
2000
1500
1000
brightness / photons
10
8
6
4
2
localizations per pixel
100
80
60
40
20
0
z_SMLM / nm

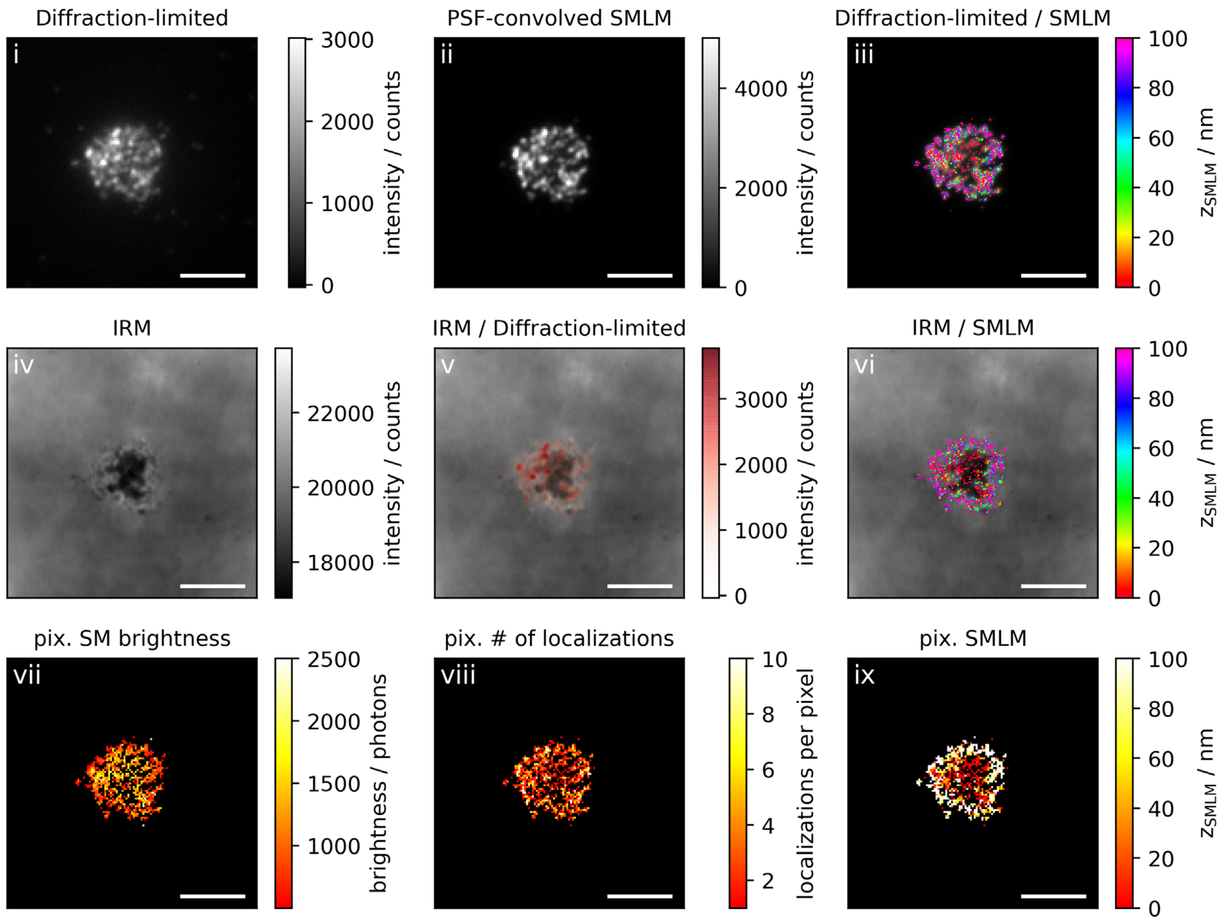
Diffraction-limited
i
3000
2000
1000
0
intensity / counts
PSF-convolved SMLM
ii
4000
2000
0
intensity / counts
Diffraction-limited / SMLM
iii
100
80
60
40
20
0
$z_{SMLM}$ / nm
IRM
iv
22000
20000
18000
intensity / counts
IRM / Diffraction-limited
v
3000
2000
1000
0
intensity / counts
IRM / SMLM
vi
100
80
60
40
20
0
$z_{SMLM}$ / nm
pix. SM brightness
vii
2500
2000
1500
1000
brightness / photons
pix. # of localizations
viii
10
8
6
4
2
localizations per pixel
pix. SMLM
ix
100
80
60
40
20
0
$z_{SMLM}$ / nm

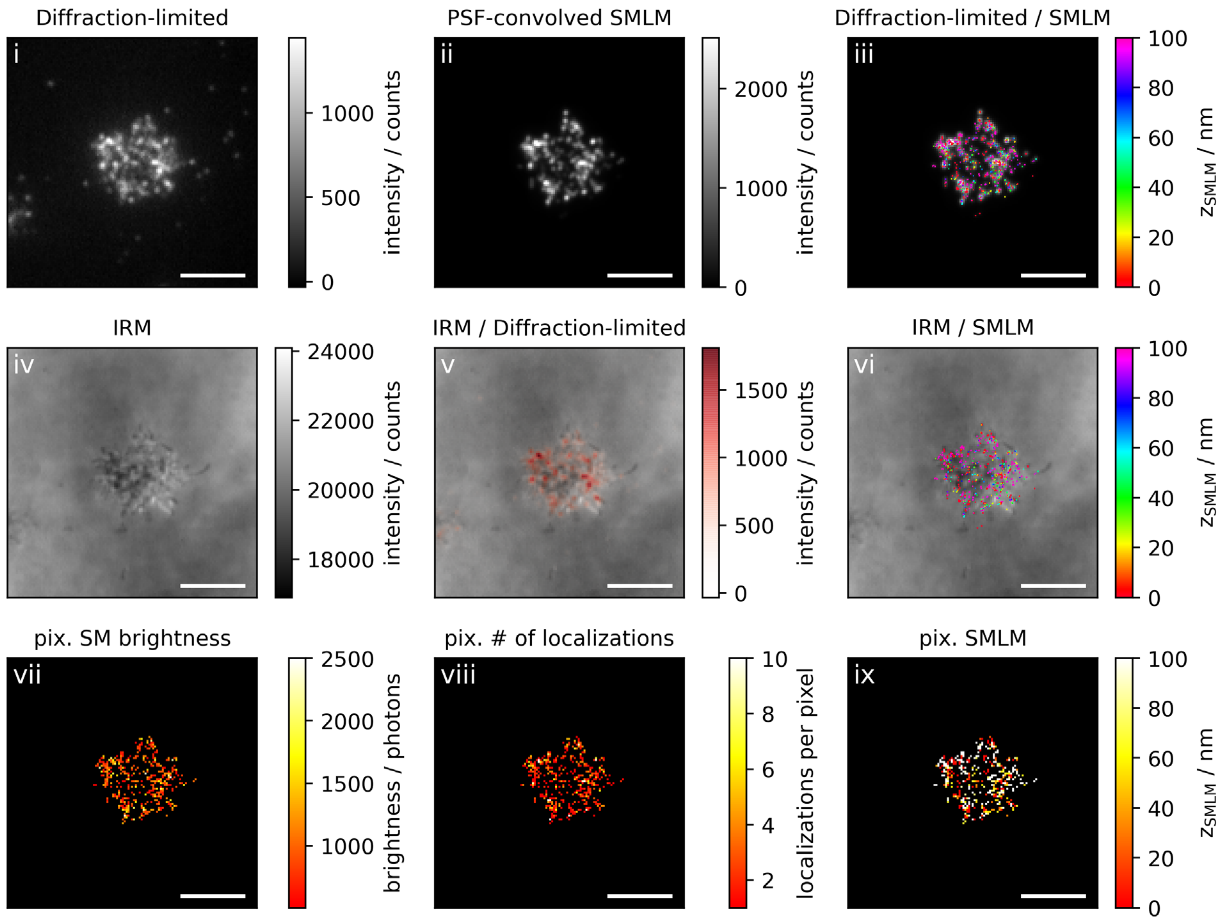
Diffraction-limited
i
1000
500
0
intensity / counts
PSF-convolved SMLM
ii
2000
1000
0
intensity / counts
Diffraction-limited / SMLM
iii
100
80
60
40
20
0
$z_{SMLM}$ / nm
IRM
iv
24000
22000
20000
18000
intensity / counts
IRM / Diffraction-limited
v
1500
1000
500
0
intensity / counts
IRM / SMLM
vi
100
80
60
40
20
0
$z_{SMLM}$ / nm
pix. SM brightness
vii
2500
2000
1500
1000
brightness / photons
pix. # of localizations
viii
10
8
6
4
2
localizations per pixel
pix. SMLM
ix
100
80
60
40
20
0
$z_{SMLM}$ / nm

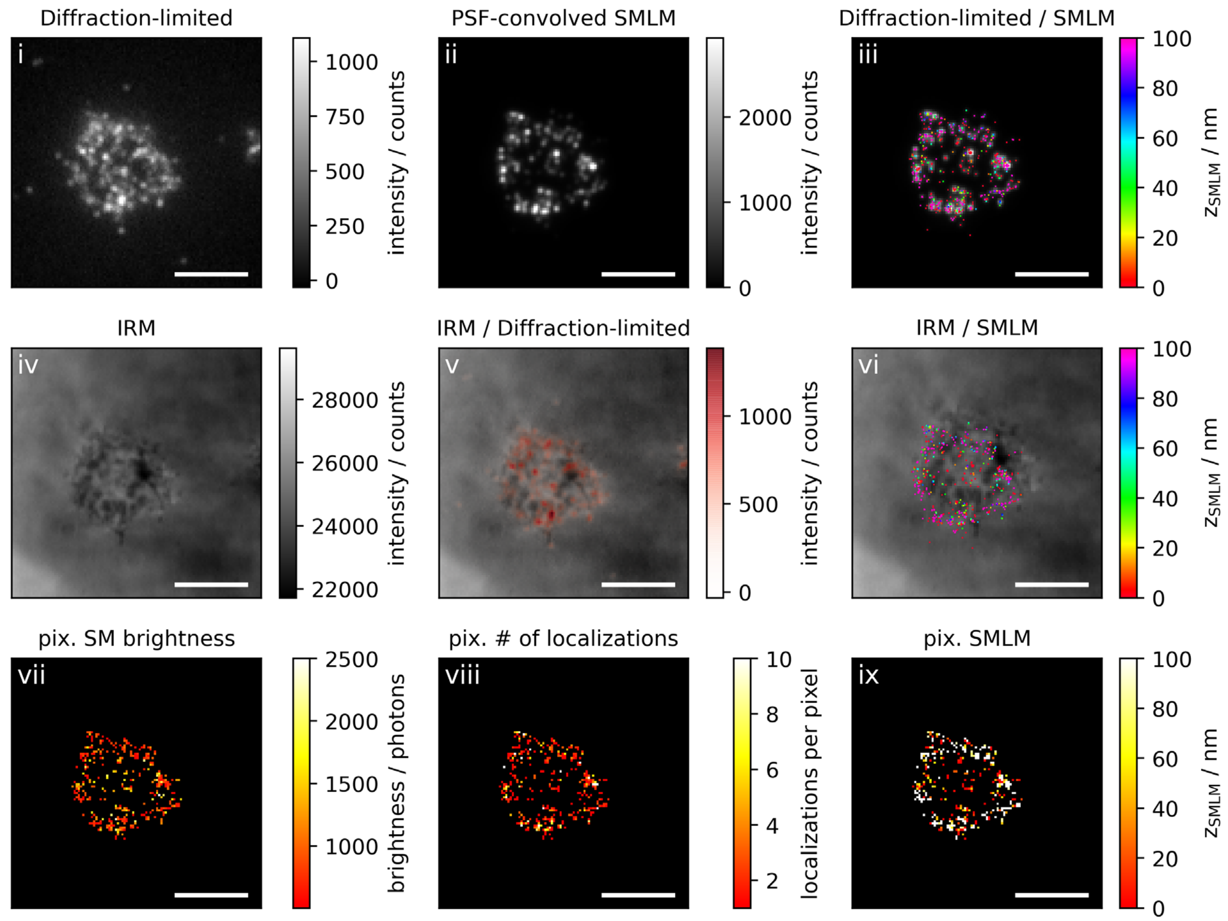
Diffraction-limited
i
1000
750
500
250
0
intensity / counts
PSF-convolved SMLM
ii
2000
1000
0
intensity / counts
Diffraction-limited / SMLM
iii
100
80
60
40
20
0
$z_{SMLM}$ / nm
IRM
iv
28000
26000
24000
22000
intensity / counts
IRM / Diffraction-limited
v
1000
500
0
intensity / counts
IRM / SMLM
vi
100
80
60
40
20
0
$z_{SMLM}$ / nm
pix. SM brightness
vii
2500
2000
1500
1000
brightness / photons
pix. # of localizations
viii
10
8
6
4
2
localizations per pixel
pix. SMLM
ix
100
80
60
40
20
0
$z_{SMLM}$ / nm

**Figure 4c: Non-activating conditions, low ICAM-1 density, fixation: 10-15 min post seeding**

**Correlative 3D-SMLM, IRM, and diffraction-limited TIR microscopy of the immunological synapse.** T cells were seeded on an SLB functionalized with low density of ICAM-1, and fixed 10-15 minutes post seeding. The T cell was imaged with IRM and fluorescence microscopy: (i) Diffraction-limited TIR image of the T cell. (ii) Reconstruction of the diffraction-limited image by convolving the 3D-SMLM image with the corresponding psf. (iii) Overlay of the diffraction-limited TIR image with the 3D-SMLM image. Color-code indicates distance to the coverslip $z_{SMLM}$. (iv) IRM image. (v) Overlay of the IRM image with the diffraction-limited image. (vi) Overlay of the IRM and the 3D-SMLM image. Bottom row images were generated by calculating the pixel-wise average of the 3D-SMLM images (pixel size of 146nm is consistent with diffraction-limited image) according to pixelated mean single molecule (SM) intensity (vii), pixelated number of localizations (viii) and pixelated mean $z_{SMLM}$ (ix). Scale bars 5 µm.

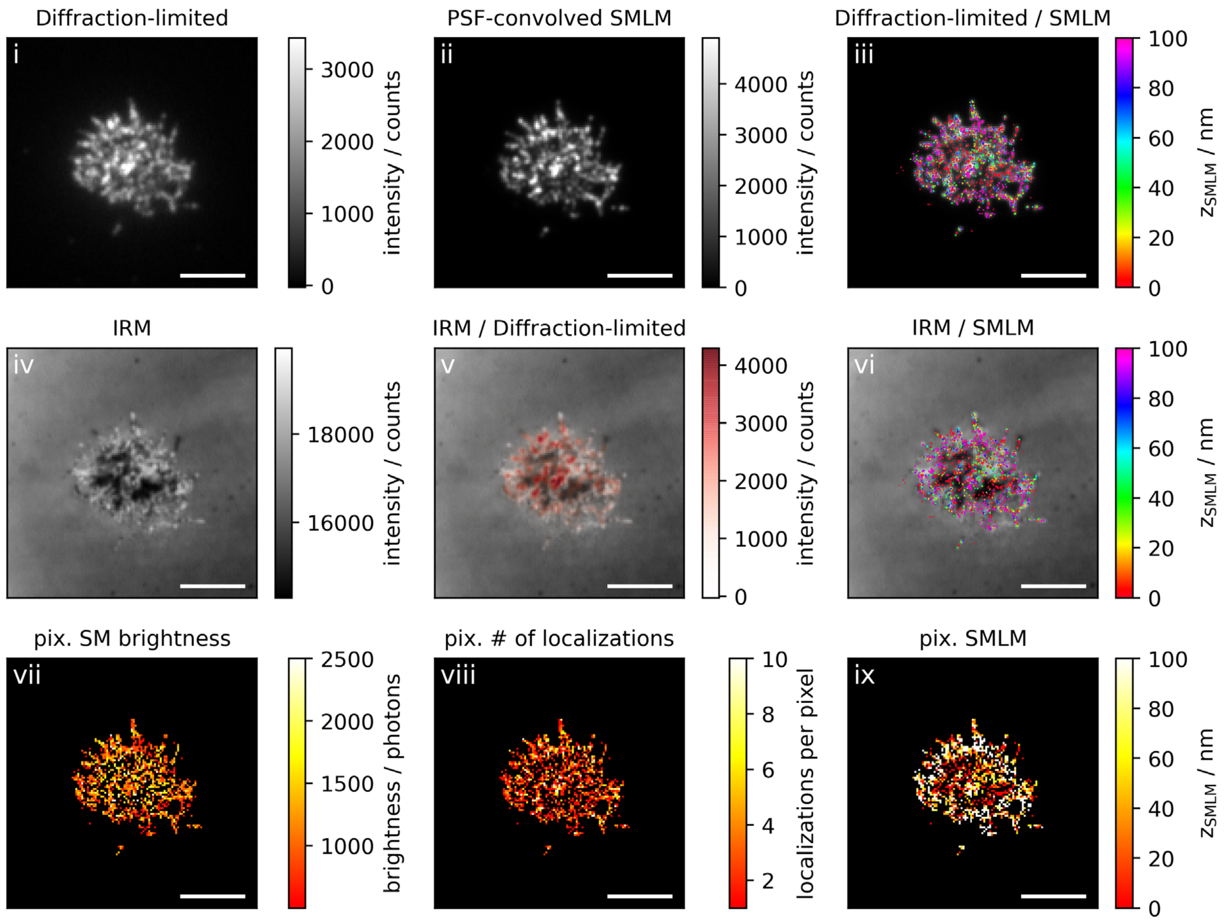
Diffraction-limited
i
intensity / counts
PSF-convolved SMLM
ii
Diffraction-limited / SMLM
iii
$z_{SMLM}$ / nm
IRM
iv
IRM / Diffraction-limited
v
IRM / SMLM
vi
pix. SM brightness
vii
brightness / photons
pix. # of localizations
viii
localizations per pixel
pix. SMLM
ix

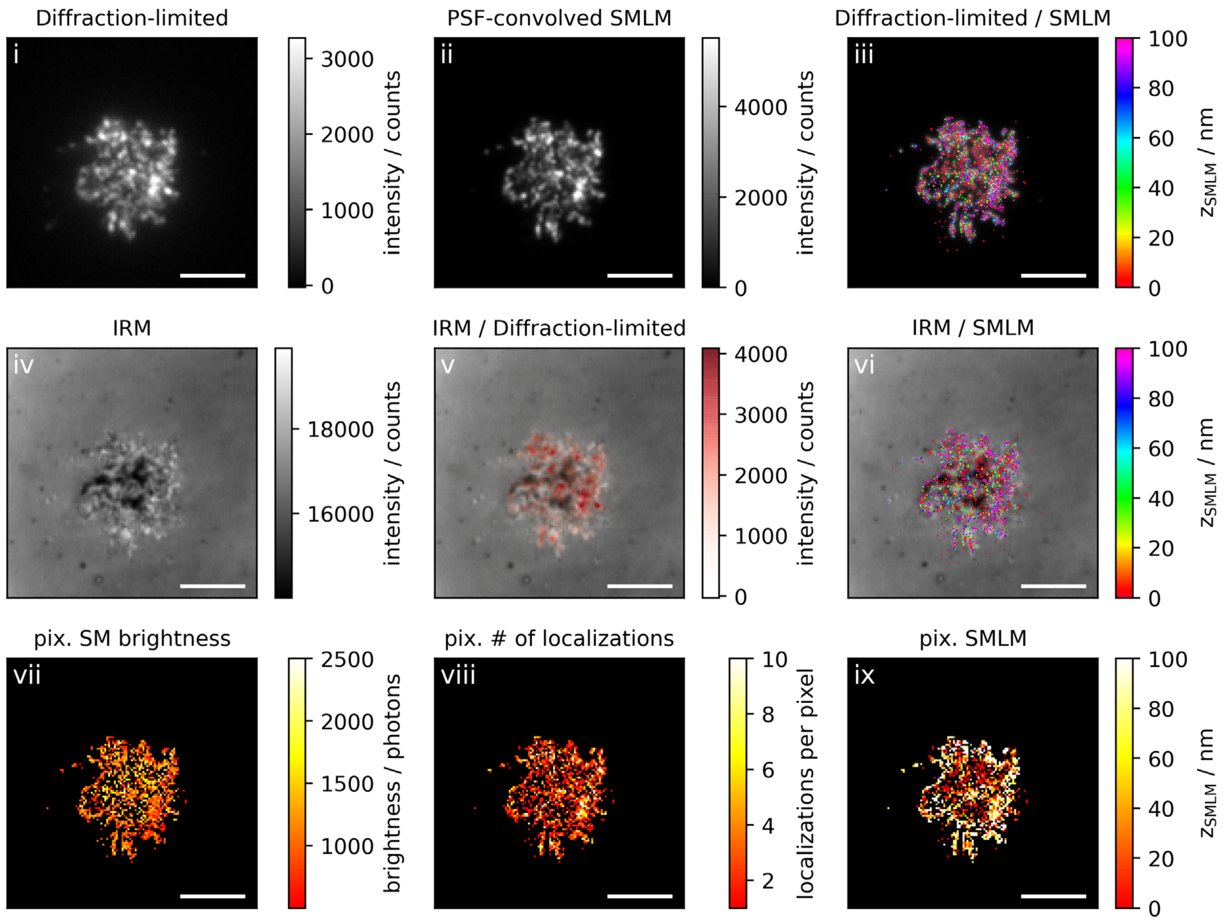

Diffraction-limited
i
intensity / counts
PSF-convolved SMLM
ii
Diffraction-limited / SMLM
iii
$z_{SMLM}$ / nm
IRM
iv
IRM / Diffraction-limited
v
IRM / SMLM
vi
pix. SM brightness
vii
brightness / photons
pix. # of localizations
viii
localizations per pixel
pix. SMLM
ix

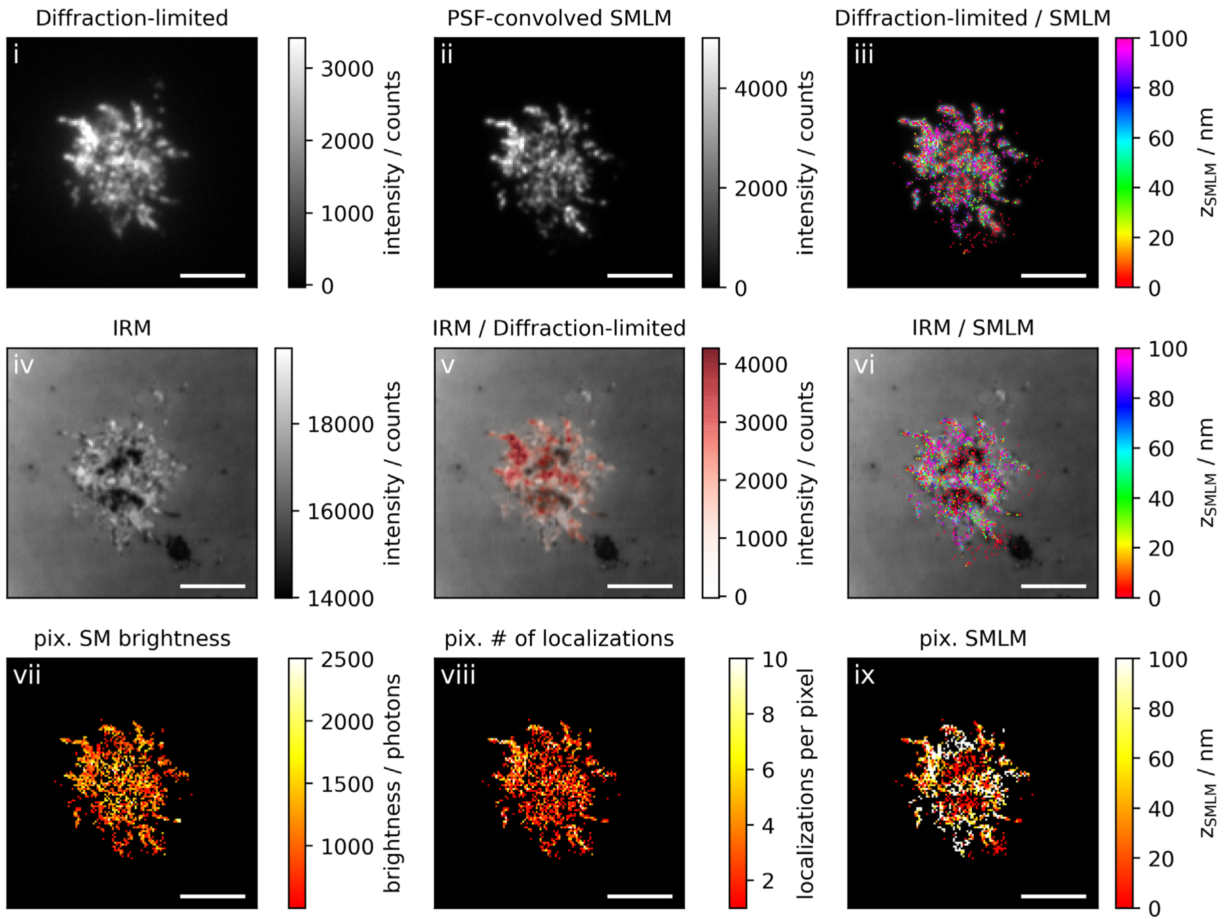
Diffraction-limited
i
intensity / counts
PSF-convolved SMLM
ii
Diffraction-limited / SMLM
iii
$z_{SMLM}$ / nm
IRM
iv
IRM / Diffraction-limited
v
IRM / SMLM
vi
pix. SM brightness
vii
brightness / photons
pix. # of localizations
viii
localizations per pixel
pix. SMLM
ix

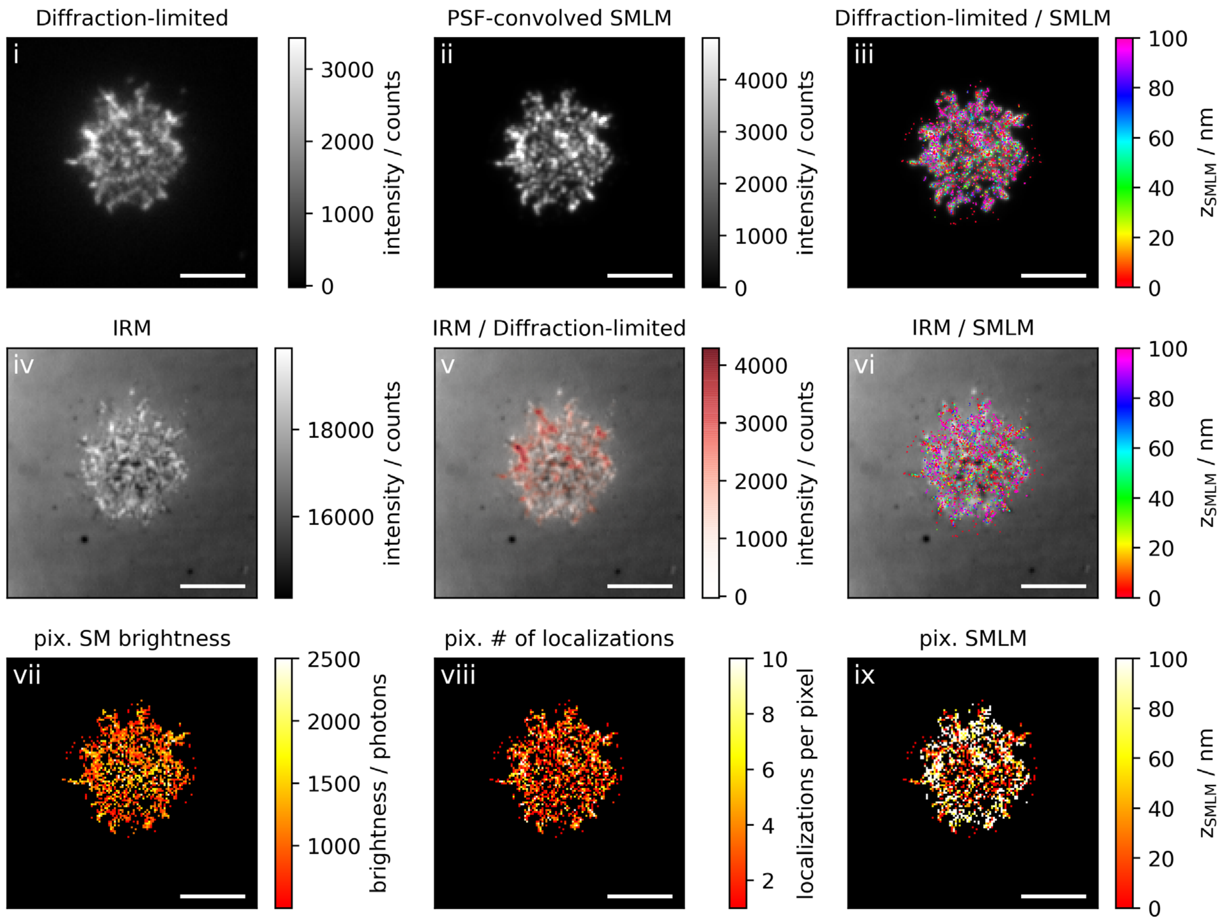
Diffraction-limited
i
intensity / counts
PSF-convolved SMLM
ii
Diffraction-limited / SMLM
iii
$z_{SMLM}$ / nm
IRM
iv
IRM / Diffraction-limited
v
IRM / SMLM
vi
pix. SM brightness
vii
brightness / photons
pix. # of localizations
viii
localizations per pixel
pix. SMLM
ix

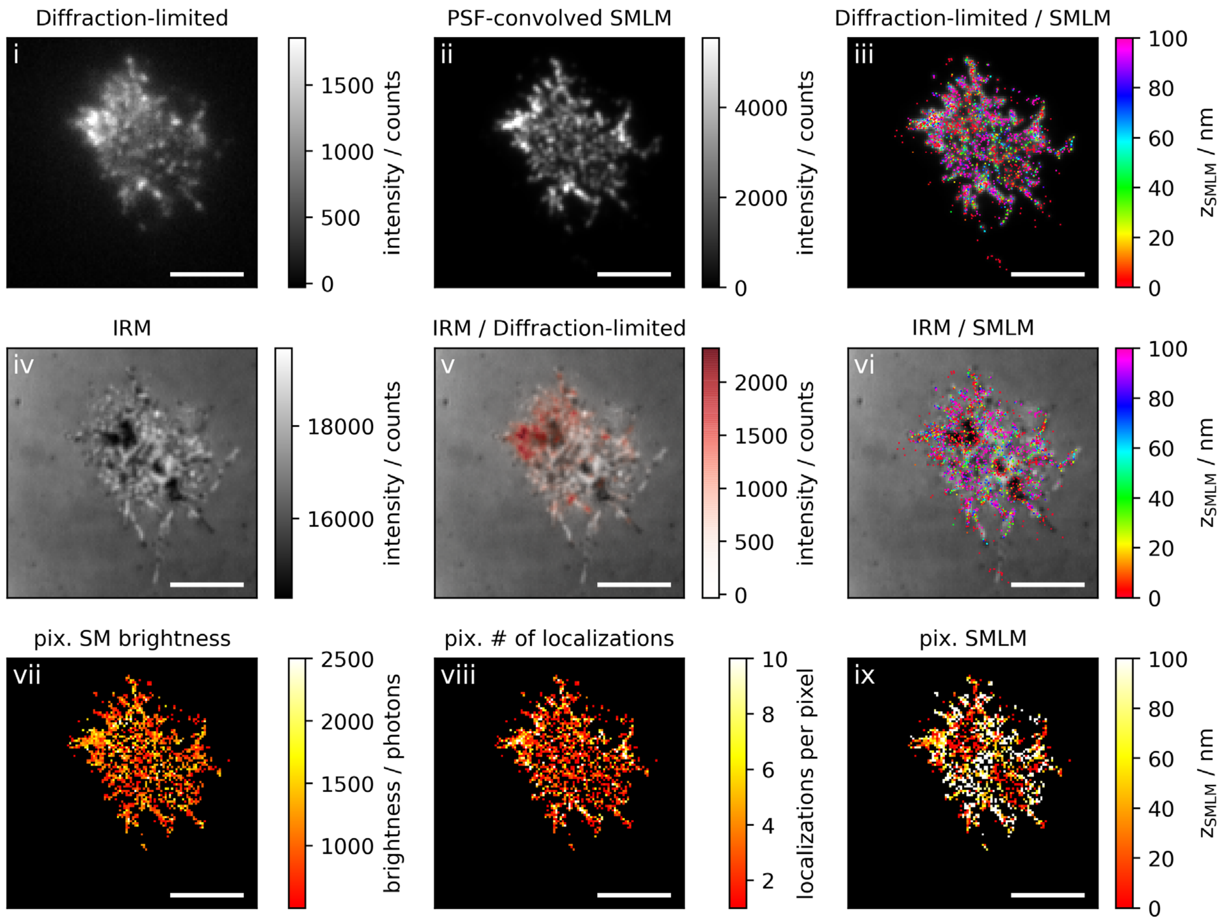
Diffraction-limited
i
1500
1000
500
0
intensity / counts
PSF-convolved SMLM
ii
4000
2000
0
intensity / counts
Diffraction-limited / SMLM
iii
100
80
60
40
20
0
$z_{SMLM}$ / nm
IRM
iv
18000
16000
intensity / counts
IRM / Diffraction-limited
v
2000
1500
1000
500
0
intensity / counts
IRM / SMLM
vi
100
80
60
40
20
0
$z_{SMLM}$ / nm
pix. SM brightness
vii
2500
2000
1500
1000
brightness / photons
pix. # of localizations
viii
10
8
6
4
2
localizations per pixel
pix. SMLM
ix
100
80
60
40
20
0
$z_{SMLM}$ / nm
